# 2,2-Di-Fluoro-Derivatives of Fucose Can Inhibit Cell Surface Fucosylation Without Causing Slow Transfer to Acceptors

**DOI:** 10.1101/2024.07.30.605762

**Authors:** Yanyan Liu, Igor Sweet, Geert-Jan Boons

## Abstract

Fucosyltransferases (FUTs) are enzymes that transfer fucose (Fuc) from GDP-Fuc to acceptor substrates resulting in fucosylated glycoconjugates that are involved in myriad of physiological and disease processes. Previously, it has been shown that per-*O*-acetylated 2-F-Fuc can be taken up by cells and converted into GDP-2-F-Fuc which is a competitive inhibitor of FUTs. Furthermore, it can act as a feedback inhibitor of *de-novo* biosynthesis of GDP-Fuc resulting in reduced glycoconjugate fucosylation. GDP-2-F-Fuc and several other reported analogs are slow substrates, which can result in unintended incorporation of unnatural fucosides. Here, we describe the design, synthesis, and biological evaluation of GDP-2,2-di-F-Fuc and corresponding prodrugs as inhibitor of FUTs. This compound lacks the slow transfer activity observed for the mono-fluorinated counterpart. Furthermore, it was found that GDP-2-F-Fuc and GDP-2,2-di-F-Fuc have similar K_i_ values for the various human fucosyl transferases while the corresponding phosphate prodrugs exhibit substantial differences in inhibition of cell surface fucosylation. Quantitative sugar nucleotides analysis by LC-MS indicates that the 2,2-di-F-Fuc prodrug has substantial greater feedback inhibitory activity. It was also found that by controlling the concentration of the inhibitor, varying degrees of inhibition of the biosynthesis of different types of fucosylated *N-*glycan structures can be achieved. These findings open new avenues for the modulation of fucosylation of cell surface glycoconjugates.

## INTRODUCTION

Fucosylation of glycoproteins and glycolipids is important for many biological processes such as cell signalling, cell adhesion, cell differentiation, immune response modulation and pathogen recognition fucosyltransferases (FUTs) catalyze the transfer of fucose (Fuc) from guanosine diphosphate β-L-fucose (GDP-Fuc) to an accepter glycoprotein or glycolipid.^1–3^ FUT1 and FUT2 are α1,2-fucosyltransferases responsible for the synthesis of H-type blood group antigen and related structures, whereas FUT3, FUT4, FUT5, FUT6, FUT7, and FUT9 synthesize α1,3- and α1,4-fucosylated glycans which are part of Lewis and blood group antigens.^1, 4^ FUT8 is the only α1,6-fucosyltransferase responsible for core fucosylation of *N*-glycans. Despite substantial interest, inhibition of FUTs for research or therapeutic purposes remains challenging.^5–8^ Two main approaches have been pursued for the development of inhibitors for FUTs. The first approach involves high-throughput screening of compounds that structurally are not related to fucosyl-transfer substrates,^9, 10^ some of which exhibit K_i_ values in the nanomolar range.^11^ Due to a lack of structural similarity to fucose, these compounds may exhibit off target effects. A second direction focuses on substrate analogues as competitive inhibitors of GDP-Fuc.^5, 7, 8^ Notable examples include GDP-2-F-Fuc,^6^ GDP-6-F-Fuc,^6^ and GDP-carba-Fuc.^12^ However, the high negative charge of nucleotide sugar analogs hampers efficient penetration of cell membranes, limiting utility in living cells. Various approaches have been explored to overcome this limitation, which include the development of per-acetylated-Fuc derivatives and phosphate prodrugs.^13–17^ A key discovery was that per-*O*-acetylated 2-F-Fuc (1, Fig. 1) can be taken up by cells and be converted into GDP-2-F-Fuc (**5**) by enzymes of the salvage pathway.^13^ The resulting compound is a competitive inhibitor of GDP-Fuc thereby blocking the activity of FUTs. Furthermore, it is a feedback inhibitor of *de-novo* synthesis of GDP-Fuc which in turn reduces glycan fucosylation.

**Figure 1.**
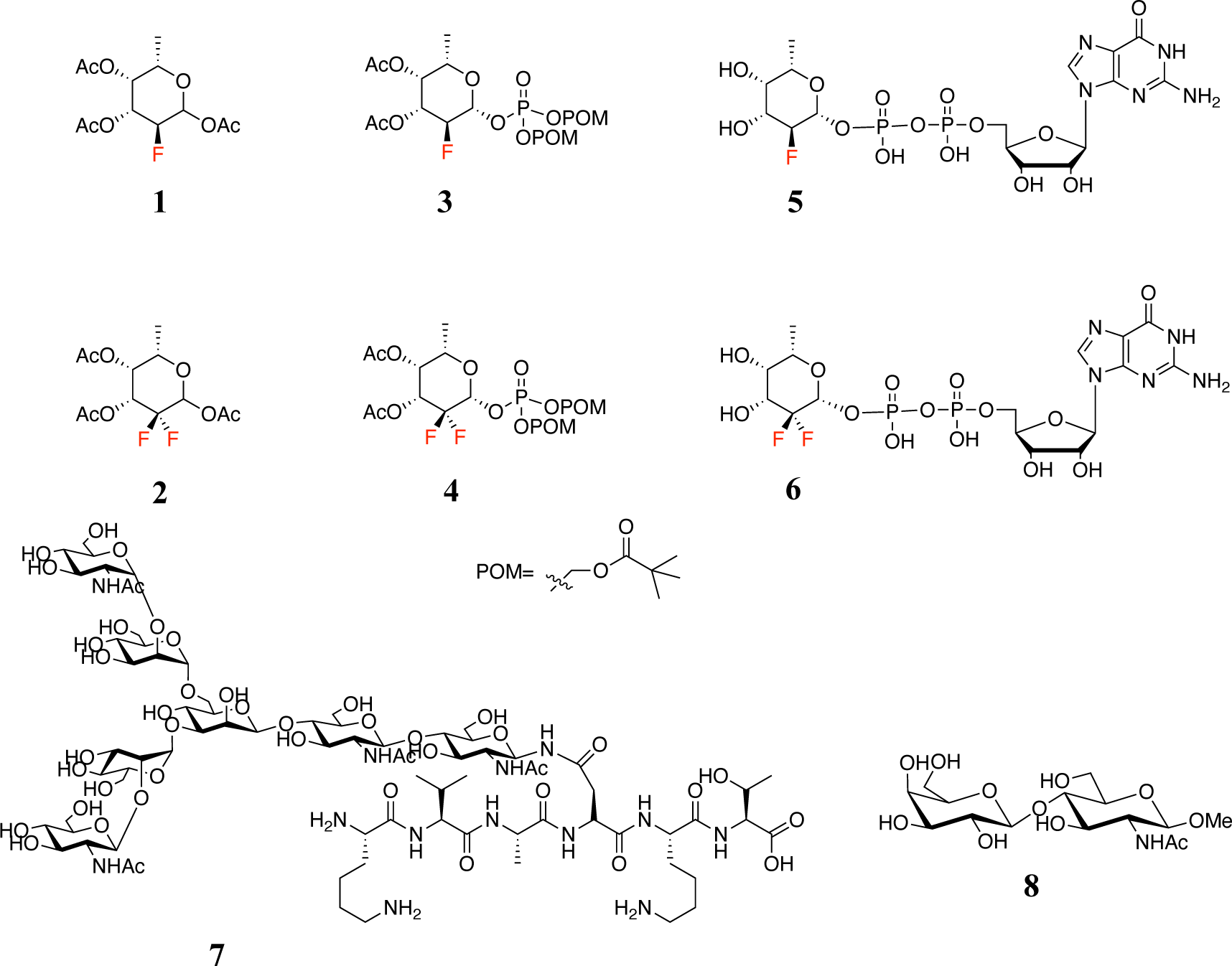
Chemical structures of inhibitors of FUTs and substrates for testing their inhibitory activity.

The potency of GDP-2-F-Fuc is modest often requiring high concentrations (∼200 µM) for substantial reduction in cell surface fucosylation. Moreover, GDP-2-F-Fuc acts as a slow substrate for several FUTs,^6, 13^ which may result in unintentional incorporation of the modified fucoside into glycan acceptors. Other fucose derivatives that can be taken up by cells and converted into GDP-Fuc analogs also exhibit slowly transferred to glycan acceptors.^14, 15, 18^

We reasoned that GDP-2,2-di-F-Fuc (**6**, Fig. 1) may be a competitive inhibitor of fucosyltransferases. It was expected that this compound would be resistant to slow transfer to glycan acceptors because the additional electron withdrawing fluorine atom at C2 further destabilizes the partial positive charge that is developed in the transition state of transfer of Fuc of GDP-Fuc to an acceptor substrate. For intracellular application, prodrugs **2** and **4** were prepared and evaluated. The incorporation of the bis(pivaloyloxymethyl) (POM) group was inspired by the successful application of several FDA-approved antiviral drugs such as adefovir dipivoxil.^19^ The POM carbonate can be removed by intracellular by esterases resulting in an unstable carboxylate intermediate that undergoes chemical degradations to a phosphate monoester. The POM moiety has been successfully applied as a prodrug for 6-CF_3_-Fuc.^15^ 2-F-Fuc derivatives **1** and **3** were also synthesized for comparison purposes. Compounds **7** and **8** were prepared to function as acceptors of the various human FUTs.

We found that GDP-2,2-di-F-Fuc (**6**) is a potent inhibitor of various human FUTs and can substantially reduce cellular GDP-Fuc concentrations by inhibition of *de novo* biosynthesis. Peracetyl-2,2-di-F-Fuc (**2**) did not reduce cellular fucosylation while prodrug **4** exhibited potent inhibition of fucosylation of cell surface glycoproteins. Furthermore, it was found that this compound is much more potent compared to mono-fluorinated **1** and **3**. Importantly, glycosyl acceptors **7** could not be modified by GDP-2,2-di-F-Fuc (**6**) when exposed FUT8 for extended periods of time. Collectively, the results demonstrate that di-fluorination of fucose eliminates the risk of unintended incorporation of the fucose analog into glycoproteins and gives compound of higher inhibitory activity in cell-based studies.

## RESULTS AND DISCUSSION

### Preparation of Fucosylation Inhibitors

First, we synthesized the series of 2-F-Fuc derivatives (Scheme 1A).^6, 13, 20^ Compound **1** was treated with hydrogen bromide in acetic acid resulting in the formation of fucosyl bromide which was reacted with di-benzyl phosphate in the presence of silver carbonate resulting in S_N_2 displacement of the bromide to afford **9** as only the β-anomer in a yield of 70% over 2 steps. Compound **9** was deprotected by a two-step procedure entailing hydrogenation over Pd/C and deacetylation by Et_3_N, resulting in the formation of **10** in a yield of 88%. Compound **10** was coupled with GMP-morpholidate in pyridine and catalyzed by 1H-tetrazole leading to the desired GDP-2-F-Fuc (**5**) in a yield of 18%. The low yield of this step is common for this type of condensation and proceeds slow with substantial starting materials remaining even after a prolonged reaction time of 5-7 days which is followed by a challenging purification. 2-F-Fuc-prodrug **3** was synthesized from **9** by a two-step procedure involving removal of the benzyl esters of the phosphate by catalytic hydrogenation over Pd/C to give a mono-phosphate that was protected by POM by reaction with POMCl in acetonitrile at 60 °C (66% yield, two steps).

**Scheme 1.**
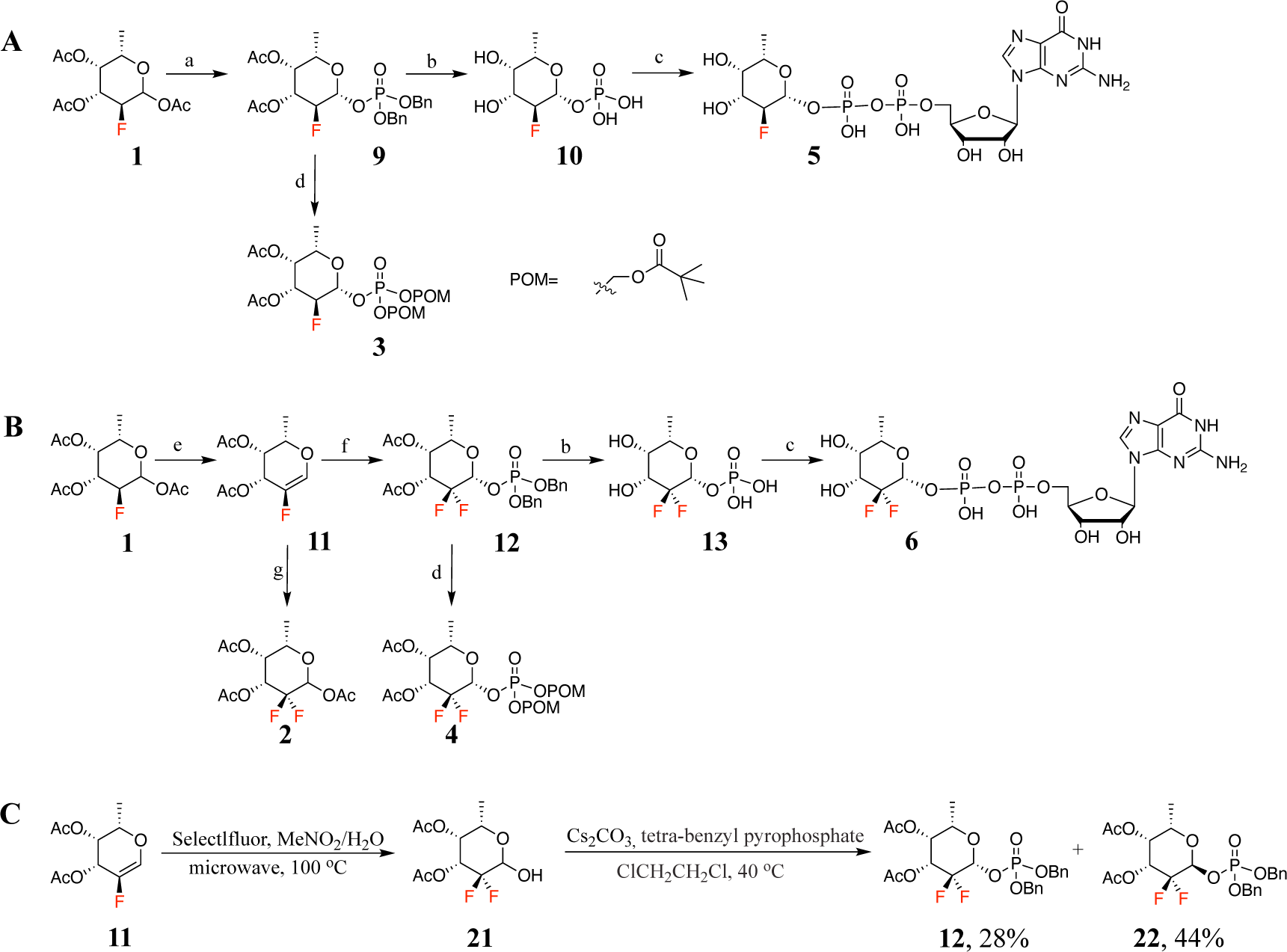
Synthesis of target compounds. a) i. HBr/AcOH, DCM, ii. Ag_2_CO_3_, (BnO)_2_POOH, 3Å MS, CH_3_CN, 70%. b) i. H_2_, Pd/C, MeOH, ii. MeOH, H_2_O, Et_3_N, 88%. c) GMP-morpholidate, 1H-tetrazole, pyridine, 18%. d) i. H_2_, Pd/C, MeOH, ii. Ag_2_CO_3_, POMCl, CH_3_CN, 60 °C, 66%. e) i. HBr/AcOH, ii. Et_3_N, CH_3_CN, 82°C, 52%. f) i. Selectfluor, MeNO_2_/H_2_O, microwave heating, 100°C, ii. Cs_2_CO_3_, tetra-benzyl pyrophosphate, ClCH_2_CH_2_Cl, 40 °C, 28% β-anomer. g) i. Selectfluor, MeNO_2_/H_2_O, microwave heating, 100°C, ii. Ac_2_O, pyridine, 85%.

The synthesis of the 2,2-di-F-Fuc derivatives is outlined in Scheme 1B and involved a vinyl fluoride as key intermediate for the introduction of the second fluorine atom at C2.^21, 22^ We explored several methodologies, such as constructing of the fucose ring from a four-carbon difluoro derivatives.^23^ However, these approaches were ineffective, and we focused on an approach involving a vinyl fluoride intermediate. Thus, treatment of **1** with hydrogen bromide in acetic acid resulted in the formation of an intermediate anomeric bromide that was treated with triethylamine in acetonitrile to induce elimination and formation of vinyl fluoride **11** in a yield of 52%.^24^ Electrophilic fluorination of **11** with Selectfluor in a mixture of nitromethane and H_2_O, followed by acetylation of the resulting anomeric hydroxyl produced per-*O*-acetyl-2,2-di-F-Fuc (**2**) in a yield of 85%. To achieve this result, the reaction condition for this transformation was optimized. Heating the reaction mixture in an oil bath ranging from 40 °C to 80 °C for several hours to overnight resulted in product decomposition while the starting material was not fully consumed. The use of a microwave reactor instead of a traditional heating reduced the reaction time to just 5-10 min and greatly enhancing the reaction efficiency and yield.

Next, attention was focused on the preparation of compound **12** which is a key intermediate for the synthesis of the target compounds **4** and **6** (Schemes 1B and 1C). The generation of the β-anomeric phosphate linkage of **12** proved challenging and most of the examined reaction conditions gave only the α-anomer. Although there are many glycosylation methodologies available for constructing β-anomeric phosphate linkages, the difluoro modification at C2 greatly increases the stability of the chemical bond at C1, making the installation of a halogen or displacement of a leaving group at the C1 challenging. Instead of generating an oxocarbenium ion intermediate for anomeric phosphorylation, 2,2-di-F-Fuc lactol **21** was employed as nucleophile for attack on a phosphate anhydride in the presence of a base to form compound **12**. Thus, compound **11** was subjected to electrophilic fluorination using Selectfluor in a mixture of nitromethane and H_2_O which resulted in the formation of compound **21**, which is unstable and decomposed during purification and therefore was immediately employed in the next step (Scheme 1C). Various reaction conditions were explored to convert **21** into **12**, including different bases, phosphorylation reagents and temperatures, which gave the α-anomer with negligible β-anomer formation. Based on these results, we presumed that the α-anomer is the thermodynamic while the β-anomer is the kinetic product. Therefore, we expected that the use of a relative weak base and a phosphorylation reagent of lower activity may enhance the β-anomeric selectivity.^25–27^ By utilizing cesium carbonate as the base and tetrabenzyl pyrophosphate as the phosphorylation reagent at 40 °C, a separable mixture of the α-anomer (44%, **22**) and β-anomer was obtained (28%, **12**).

Compound **12** was deprotection by a two-step process involving hydrogenation over Pd/C followed by deacetylation with Et_3_N yielding compound **13** in 88% yield. Subsequently, compound **13** was condensed with GMP-morpholidate in pyridine and catalyzed by 1H-tetrazole, resulting in the production of the desired GDP-2,2-diF-Fuc (**6**). 2,2-diF-Fuc prodrug **4** was synthesized from **12** by removal of the benzyl protecting groups by hydrogenation over Pd/C, resulting a mono-phosphate group intermediate that was acylated with POMCl resulting in **4** in a yield of 66% over 2 steps.

The preparation of glycosyl acceptors **7** and **8** is described in the supporting information. (Fig. S1 and Scheme S1).

We also investigated enzymatic approaches to synthesize compounds **5** and **6** (Scheme S2). Bacterial L-fucokinase/GDP-fucose pyrophosphorylase (FKP) is a bifunctional enzyme that catalyzes the conversion of fucose to Fuc-1-P and subsequently to GDP-Fuc. FKP exhibits promiscuity towards L-fucose derivatives including 2-F-Fuc^13, 28^. Compound **5** could easily be synthesized by treatment of **10** with GTP in the presenec of FKP. Unfortunately, a similar transformation using 2,2-difluoro-Fuc failed. Interestingly, FKP could readily convert 2,2-di-fluoro-Fuc-1-P (**14**) obtained by chemical into **6**, which indicated that the first enzymatic step is most sensitive to chemical modifications.

### Inhibition of Six Human FUTs by Fluorinated GDP-Fuc Analogs 5 and 6

Next, attention was focused on determining the inhibitory activity of the GDP-mono- (**5**) and di- (**6**) fluoro-Fuc derivatives. The disaccharide LacNAc **8** was used as the acceptor for FUT1, 3, 5, 6, and 9 while the bi-antennary glycan **7** was employed as the acceptor for FUT8.^8, 12, 29–36^ Kinetic parameters were obtained by keeping the acceptors (**7** and **8**) at saturating concentrations while varying the concentrations of GDP-Fuc (3 to100 µM) and fluorinated inhibitors **5** and **6** (3 to 300 µM). Following an incubation period of 1 h, reaction conversions were determined using a commercially available GDP-Glo glycosyltransferase assay kit. The data was analyzed using Prism and fitted by nonlinear regression. The results are summarized in Table 1.

**Table 1.**
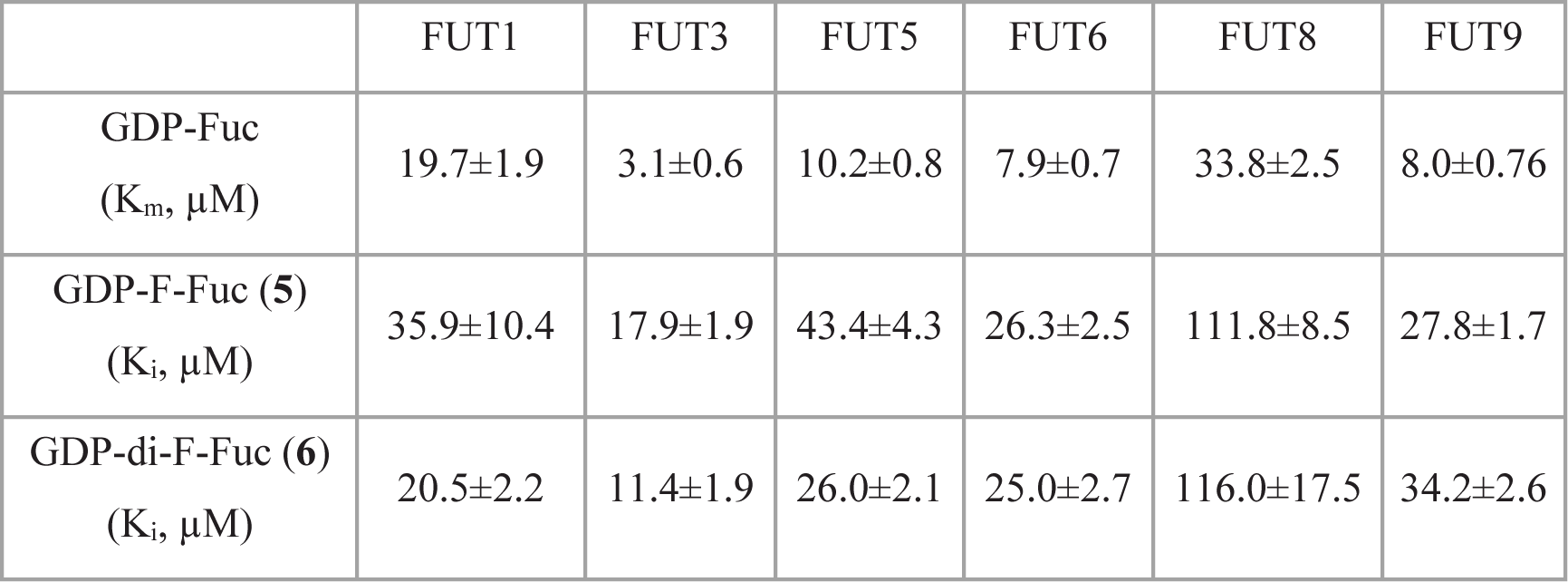
Inhibition of human FUTs by GDP-2-F-Fuc and GDP-2,2-di-F-Fuc.

|  | FUT1 | FUT3 | FUT5 | FUT6 | FUT8 | FUT9 |
| --- | --- | --- | --- | --- | --- | --- |
| GDP-Fuc<br>($K_m$, $\mu M$) | 19.7±1.9 | 3.1±0.6 | 10.2±0.8 | 7.9±0.7 | 33.8±2.5 | 8.0±0.76 |
| GDP-F-Fuc (**5**)<br>($K_i$, $\mu M$) | 35.9±10.4 | 17.9±1.9 | 43.4±4.3 | 26.3±2.5 | 111.8±8.5 | 27.8±1.7 |
| GDP-di-F-Fuc (**6**)<br>($K_i$, $\mu M$) | 20.5±2.2 | 11.4±1.9 | 26.0±2.1 | 25.0±2.7 | 116.0±17.5 | 34.2±2.6 |

The K_m_ values for the various FUTs differ considerably and FUT3 has the smallest value and FUT8 the largest while the others have intermediate inhibitory activities. Furthermore, the results demonstrate that GDP-2,2-di-F-Fuc (**6**) is a competitive inhibitor for all examined FUTs. Interestingly, **5** and **6** had similar inhibitory activities for FUT1, 3, 5, 6, and 8 whereas the K_i_ value for FUT8 is substantially higher.

A limitation of compound **5** is its slow activity as a substrate for certain FUTs, particularly FUT8.^13^ One of the aims for introducing the second fluorine atom at **6** is to increase the stability of the glycosidic bond preventing transfer. To validate this hypothesis, we set up two parallel transfer reactions catalyzed by FUT8 using *N*-glycan **7** as acceptor (300 μM) and compounds **5** and **6** (750 μM) as donor. The possible transfer of fluorinated fucose was monitored using MALDI-TOF MS at 0, 3, and 7 days (Fig. 2). After 3 days, a substantial amount of **5** had been transferred to 7 while after 7 days, the reaction had gone to near completion. In contrast, no product formation was observed for compound **6** and even after 7 days, no observable signals emerged for the fucosylated product indicating that GDP-di-F-Fuc does not act as a donor.

**Figure 2.**
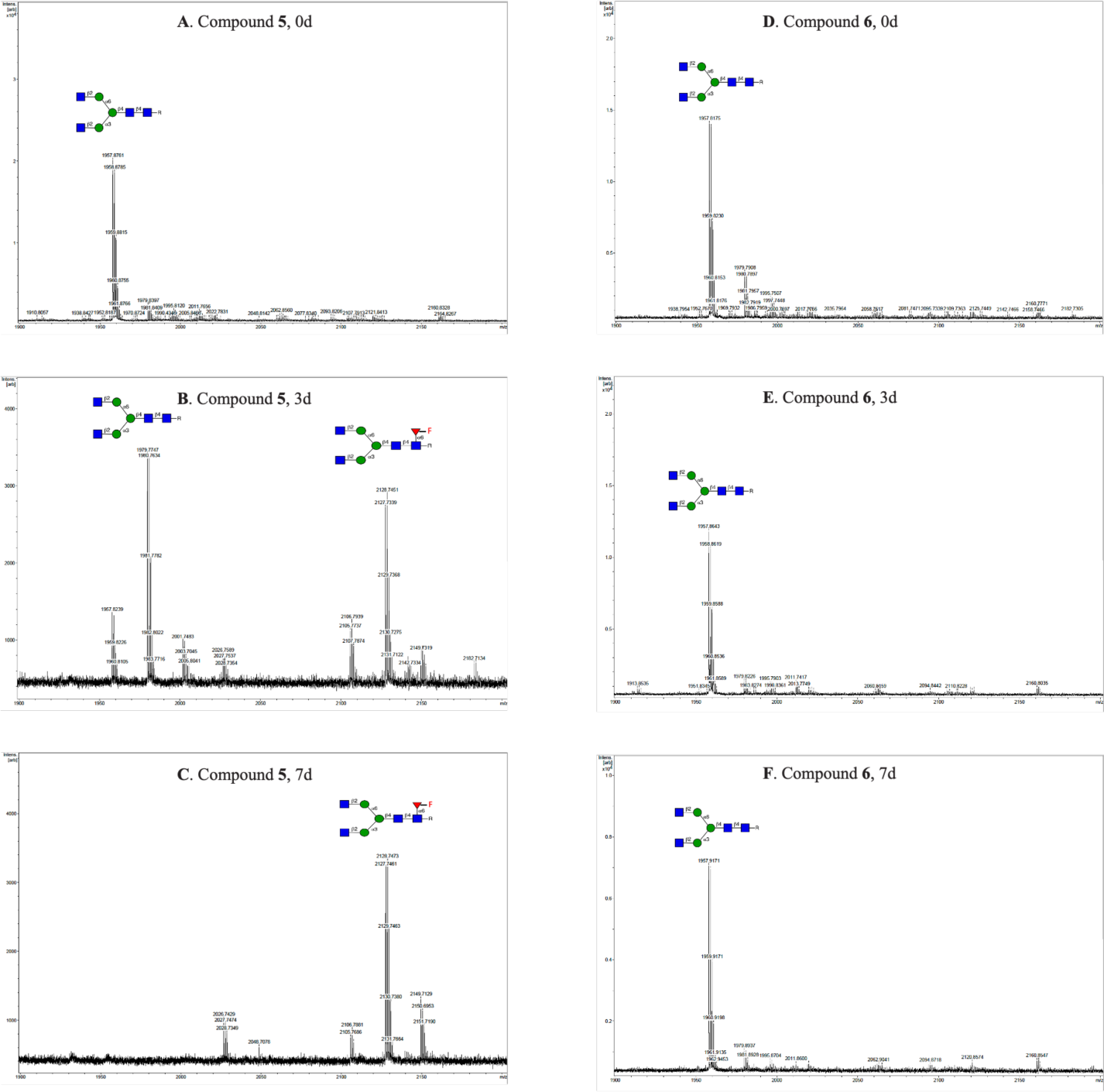
MALDI-TOF analysis of two parallel FUT8-catalyzed transfer reactions in the presence of 100 mM Tris buffer and 10 mM MgCl_2_. **A**-**C**, Time points for compound **5** as donor substrate. **D**-**F**, Time points for compound **6** as donor.

### Inhibition of Cell Surface Fucosylation

Although fluorinated GDP-Fuc analogues exhibit potent inhibitory activity, these compounds cannot be employed for cell-based inhibition studies due to poor cell permeability. A prodrug strategies based on the biosynthetic pathway of nucleotide sugars has been developed to improve the cell permeability of Fuc analogs.^37–40^ In this respect, GDP-Fuc can be biosynthesized by *de novo* and salvage pathway (Fig. 2).^1, 41–44^ Quantitative examination of the fucose metabolism in HeLa cells has shown that over 90% of GDP-Fuc originates from *de novo* biosynthesis.^45^ It starts by conversion of GDP-mannose into GDP-4-keto-6-deoxymannose catalyzed by GDP-mannose-4,6-dehydratase (GMD) which is converted into GDP-Fuc by a dual functional epimerase-reductase known as the FX protein.^44^ GDP-Fuc acts as an inhibitor of GMD thereby blocking the *de novo* pathway and regulating GDP-Fuc levels. In the salvage pathway, GDP-Fuc is biosynthesized from free fucose derived from extracellular or lysosomal sources by anomeric phosphorylation by fucose kinase to give Fuc-1-phosphate that is converted into GDP-Fuc by GDP-Fuc pyrophosphorylase (GFPP).

The salvage pathway has been exploited to convert fucose analogs that can penetrate the cell membrane into the corresponding GDP-Fuc derivatives.^13–17^ Here, we tested compounds **1** and **2** as inhibitors of fucosylation of cell surface glycoconjugates. After cell uptake, esterases can remove the acetyl esters and the resulting compounds can then be converted into the corresponding GDP-Fuc derivatives by fucose kinase and GFPP (Fig. 3). Fucose kinase has restricted substrate specificity and therefore protected monophosphate fucose derivatives have been developed that can cross the cell membrane ^13, 15, 46^ Therefore, we also examined POM carbonates of 2-F-Fuc-1-P (**3**) and 2-di-F-Fuc-1-P (**4**) as precursors of the corresponding mono-phosphates.

**Figure 3.**
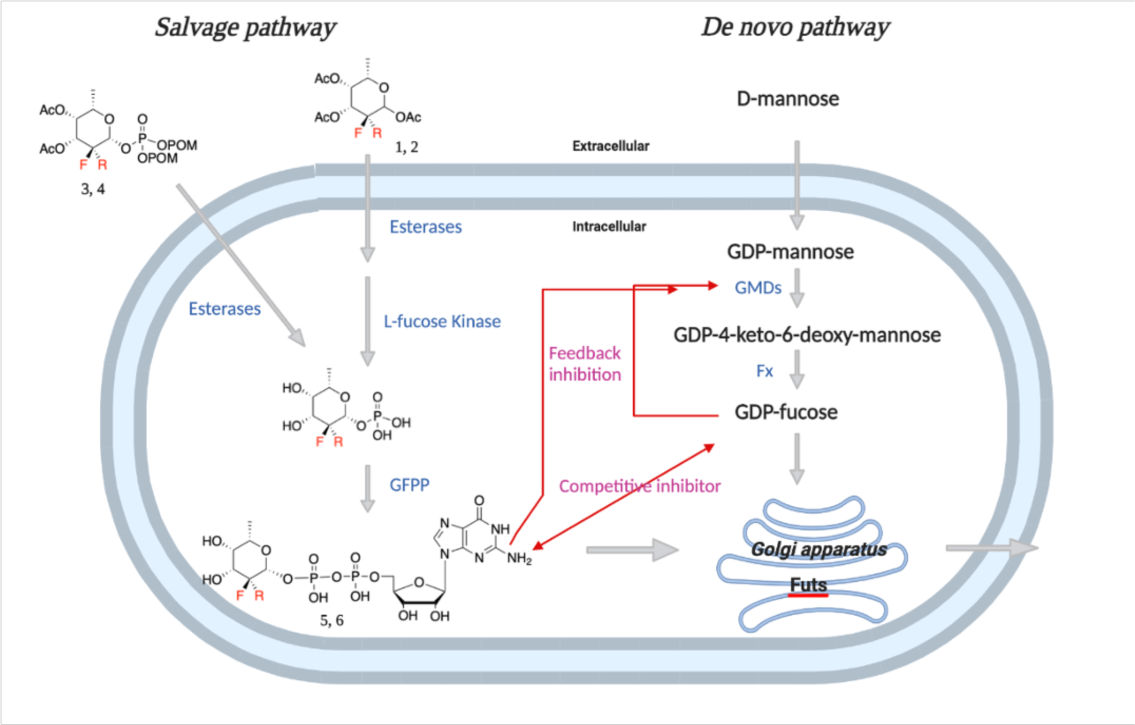
Metabolic pathway of fluorinated inhibitors in cells.

The inhibitory activity of the compounds **1**-**4** was evaluated in HL60, U-2 OS, and HEK293 cell lines.^47^ Cells were treated with prodrugs at concentrations ranging from 0.5 to 250 μM or in DMSO control. After 3 days, cells were harvested and analyzed using flow cytometry to evaluate the expression of cell surface fucosides by employing *Aleuria aurantia* lectin (AAL) and *Lens culinaris* agglutinin (LCA) and anti-Le^x^ and SLe^x^ antibodies (Fig. 4).^13, 18, 48–50^ AAL has broad binding properties and preferentially binds to α1,3-, α1,2-, α1,4-, and α1,6-linked fucosides of *N*-acetyl-lactosamine. Per-*O*-acetyl-2,2-di-F-Fuc (**2**) did not show any reduction in cell surface fucosylation while 2,2-di-F-Fuc-1-P-POM (**4**) exhibited potent inhibition. This observation indicates that fucose kinase cannot phosphorylate 2,2-di-F-Fuc whereas 2-F-Fuc is a proper substrate for this enzyme. At a concentration of 32 μM, compound **4** suppressed all most all cell surface fucosylation as demonstrated by a lack of binding of AAL and LCA in the two cell lines. In contrast, a comparable reduction in cell surface fucosylation by compounds **1** and **3** required concentrations higher than 125 μM. It is worth noting that for both HL60 and U-2 OS, a lower concentration of compound **4** was required to inhibit Le^x^/SLe^x^ expression (8 μM) compared to core fucosylation which was detected by LCA. This observation agrees with the relatively high K_m_ value for FUT8 which is responsible for core fucosylation. We also investigated the effects of the compounds on cell proliferation and viability^51, 52^ and no effects were observed at concentrations lower than 125 μM (Fig. S4).

**Figure 4.**
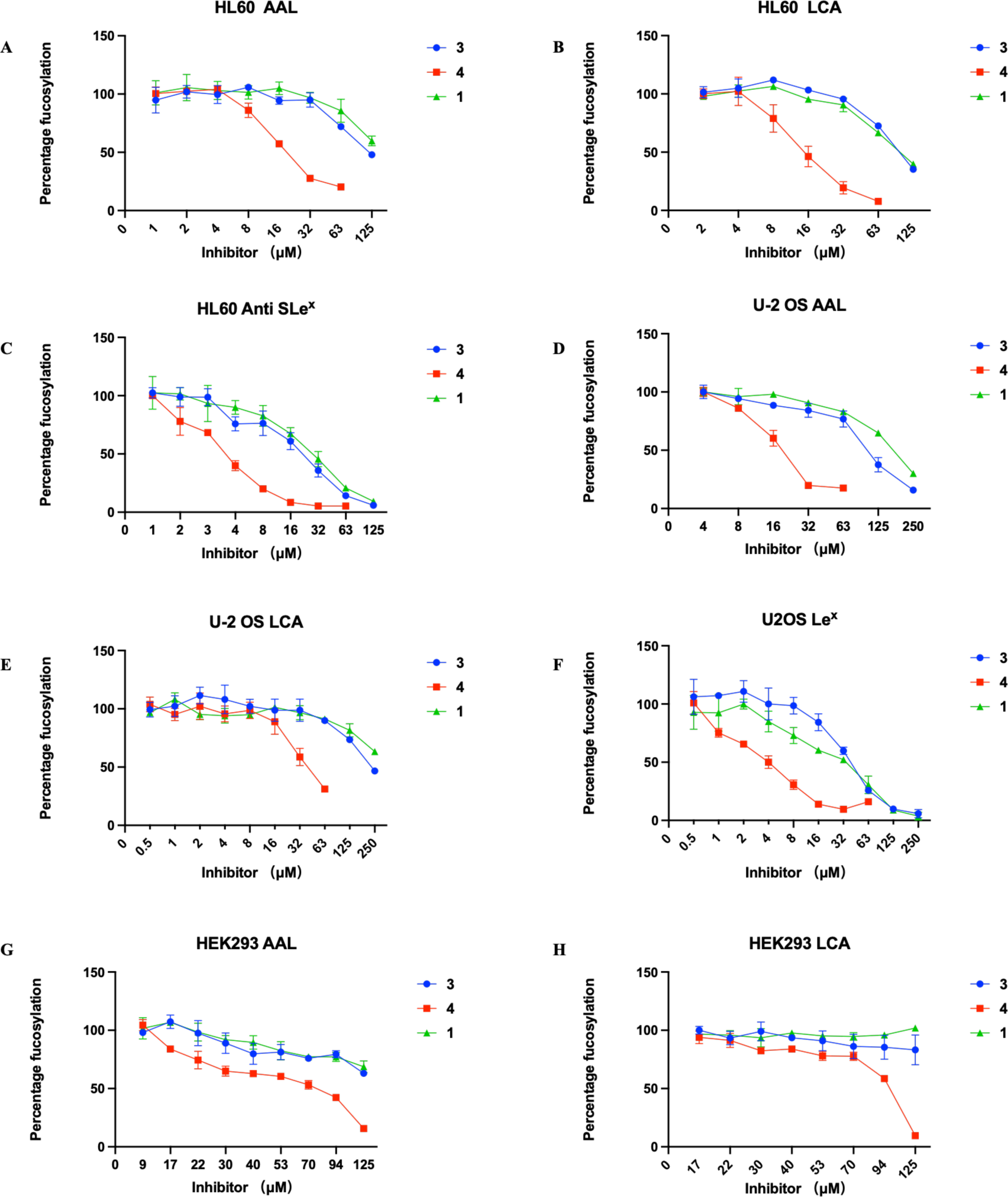
Fluorinated Fuc prodrugs act as fucosyl transferase inhibitors of cells. Cells were subjected to prodrugs at concentrations ranging from 0.5 μM to 250 μM or in DMSO control for 3 days. The data were normalized to cells treated with DMSO only as 100% fucosylation and unstained cells as 0%. The data presented are indicative of three independent experiments conducted in triplicate, illustrating the mean ± s.d. values. Error bars are displayed for each dataset.

### Glycan profiling of cells treated with inhibitor

To further investigate the impact of the fluorinated fucose derivatives on cell surface fucosylation, we performed LC-MS analysis of *N*-glycans released from HL-60 cells.^53^ Thus, the cells were treated with 8 and 32 μM of compounds **1**, **3**, and **4** and DMSO control. After 3 days, the cells were harvested and subjected to PNGase F treatment to liberate the *N*-glycans, followed by labelling with procainamide for positive-mode detection. The top 20 most abundant *N*-glycans for each sample are shown in Fig. 5 to illustrate the change in the *N*-glycan profile. To highlight the fucosylated species, only for these glycans a putative compositional structure is depicted.

**Figure 5.**
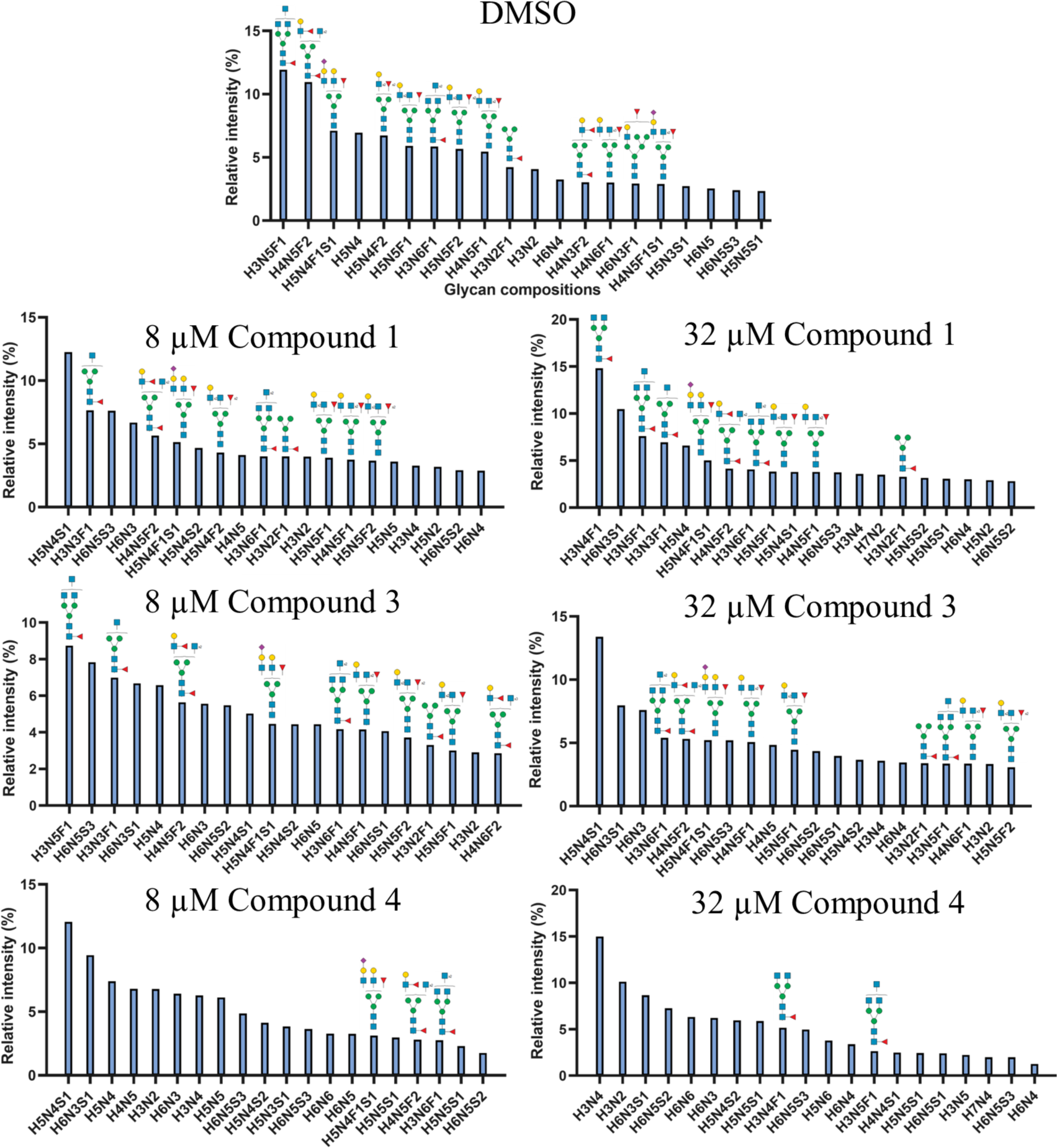
*N*-glycosylation profiles of HL-60 cells treated with compounds **1**, **3**, and **4**. The proposed fucosylated *N*-glycan structures are putative, based on monosaccharide composition (H: Hexose; N: N-acetylhexosamine; F: deoxyhexose; S: N-acetylneuraminic acid).

At a low concentration of inhibitors, compounds **1** and **3** had minimal influence on the HL-60 cell *N*-glycan profile. In contrast, compound **4** displayed a notable relative decrease in fucosylated structures with only low levels of core fucosylation detected at the highest concentration of 32 µM.

In addition to the compositional analysis, comparison of in-source fragment ions of fucosylated terminal *N*-glycan epitopes showed a drastic decrease in terminal fucosyl-LacNAc (i.e. Le^X^, H-antigen) and SLe^x^. This difference was most evident in the compound **4**-treated cells, see Fig. S5. Noteworthy is that the abundance of sialyl-LacNAc in-source fragments showed a slight increase compared to the DMSO control.

### Nucleotide Sugar Analysis of Cells Treated with 1 and 3

The observation that GDP-2-F-Fuc (**5**) and GDP-2,2-di-F-Fuc (**6**) exhibit small differences in K_i_ values for the various human fucosyl transferases, while the corresponding phosphate prodrugs **3** and **4** showed relatively large differences in inhibition of cell surface fucosylation, indicates that they have different feedback inhibition activities. To validate this hypothesis, we performed quantitative sugar nucleotides analysis by LC-MS.^54^ Different concentrations of **1**, **3**, and **4** were administered to HL60 cells in culture medium and after three days, cell lysates were collected for mass spectrometry analysis. GDP-Fuc, GDP-2-F-Fuc (**5**), and GDP-2,2-di-F-Fuc (**6**) were used as internal standards to quantify the levels of corresponding sugar nucleotides in the cells (Fig. 6). As anticipated, the intracellular concentration of GDP-Fuc decreases with increasing concentrations of inhibitors **1**, **3**, and **4** (Fig. 6A). The most substantial reduction was observed with **4** and at 8 μM it already induced a noticeable impact on the GDP-Fuc level, while at 32 μM, it reduced the GDP-Fuc concentration to 20% compared to control and at 64 μM, the presence of GDP-Fuc was almost undetectable. GDP-2-F-Fuc (**5**) was detected in cell samples treated with **1** and **3** and GDP-2,2-di-F-Fuc (**6**) was detected in cell samples exposed to **4** (Fig. 6B,C). Across the concentration range of 8 to 64 μM, the concentration of **5** increased with increasing concentrations of **1** and **3**. The data also indicates that **4** is more readily converted into GDP-Fuc analog **6** compared to the conversion of **1** and **3** into GDP-Fuc derivative **5** and exhibits more potent feedback inhibition of GDP-Fuc biosynthesis.

**Figure 6.**
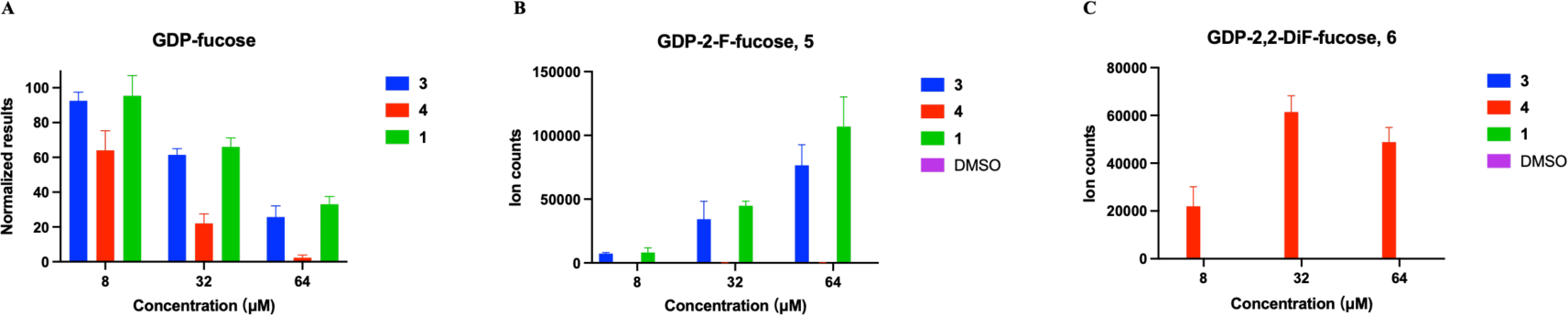
Sugar nucleotide quantification of HL60 cells treated with compounds **1**, **3**, and **4**. HL60 cells were treated with **1**, **3** or **4** and DMSO control for 3 days. **A,** Relative concentrations of GDP-Fuc in cell lysates. The data was normalized to cells incubated with DMSO at 100%. **B** and **C,** Ion counts of GDP-2-F-Fuc and GDP-2,2-di-F-Fuc detected by LC-MS in lysates of cells treated with **1**, **3**, **4** and DMSO as control. The data presented are of three independent experiments conducted as triplicates, illustrating the mean ± s.d. values. Error bars are displayed for each dataset.

## CONCLUSIONS

A synthetic methodology has been developed for the preparation of GDP-2,2-di-F-Fuc (**6**) and a corresponding prodrug (**4**), and the resulting compounds have been examined for the inhibition of fucosylation of cell surface glycoconjugates. The synthesis of the compounds was challenging due to the presence of two fluorine atoms at C-2 of fucose. The most efficient approach entailed the preparation of a lactol of 2,2-di-Fuc that was employed as a nucleophile in the presence of a weak base to react with tetra-benzyl pyrophosphate to provide the corresponding anomeric phosphate. After removal of the protecting groups, the resulting anomeric mono-phosphate could be converted into GDP-2,2-di-F-Fuc (**6**) by condensation with GMP-morpholidate. In addition, a phosphate prodrug (**4**) was prepared by the conversion of the mono-phosphate into a bis(pivaloyloxymethyl) (POM) ester. Similar compounds were prepared having only one F atom of C-2 of fucose (**3** and **5**). It was found that GDP-2-F-Fuc (**5**) and GDP-2,2-di-F-Fuc (**6**) have similar inhibitory activities for various human fucosyl transferases. While slow transfer of GDP-2-F-Fuc (**5**) was observed, the corresponding GDP-2,2-di-F-Fuc (**6**) could not modify glycosyl acceptors even after a prolonged period of incubation. This is due to the strong electronegativity of the two-fluorine atom that stabilizes the anomeric phosphate. Interestingly, the mono-fluorinated (**3**) and di-fluorinated (**4**) phosphate prodrugs showed substantial differences in potency of inhibiting cell surface fucosylation and di-fluorinated derivative **4** is more activity at lower concentrations. Examination of the concentrations of GDP-Fuc by LC-MS indicated that **4** has greater feedback inhibitory activity compared to the mono-fluorinated analog. These studies also confirmed that intracellularly, **4** is converted into GDP-2,2-di-F-Fuc (**6**). Another surprising observation was that lower concentrations of **4** are required to block the biosynthesis of Le^x^ and SLe^x^ compared to core-fucosylation of *N*-glycans. This is attributed to differences in K_m_ values of the various FUTs and in particular FUT8 which is responsible for core fucosylation has the highest K_m_ value of the examined fucosyl transferases and thus is most sensitive to lower levels of GDP-Fuc caused by the application of the inhibitors. The outcome of the described studies holds promise for exploration of the biological activities of fucosylated glycans.

## ASSOCIATED CONTENT

### Supporting Information

The Supporting Information is available free of charge at https://pubs.acs.org.

**S**ynthetic protocols, compound characterization, cell experimental, and NMR spectra (PDF).

## AUTHOR INFORMATION

### Author Contributions

Y.L. and G.J.B. designed the project. G.J.B. was responsible for overall project management. Y.L. performed all chemical and enzymatic synthesis and cellular assays. I.S. performed the MS analysis. The manuscript was written by Y.L and G.J.B. All authors reviewed the final manuscript.

### Notes

The authors declare no competing financial interest.

## ACKNOWLEDGMENTS

Dr. Kelley Moremen (University of Georgia, USA) provided the expression vectors for the various fucosyl transferases, which were expressed by Dr. Gerlof P Bosman (Utrecht Uiversity) and Mrs Linda H.C. Quarles van Ufford (Utrecht University) by reported procedures. Dr. Monique van Scherpenzeel (GlycoMScan B.V) performed quantitative analysis of the nucleotide sugar. Y.L. was supported by the Chinese Scholarship Council.

## Table of Contents Graphic

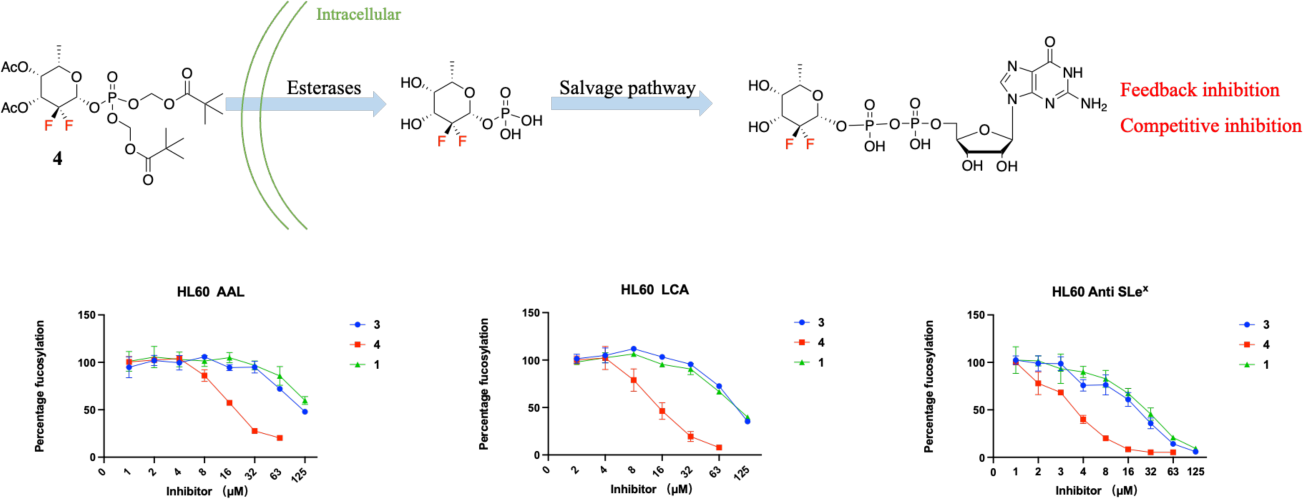

## Table of Contents Text

We have designed, synthesized, and biologically evaluated GDP-2,2-di-F-Fucose and corresponding prodrugs as inhibitor of cell surface fucosylation. The 2,2-Di-F-derivatives lack slow transfer activity to acceptors, are much more potent than the mono-fluorinated counterpart, and display some selectivity for specific FUTs.

