## Supplementary Information for "2,2-Di-Fluoro-Derivatives of Fucose Can Inhibit Cell Surface Fucosylation Without Causing Slow Transfer to Acceptors"

**Table of contents**


| | page |
|---|---|
| **1) Supplementary schemes and figures** | S3 |
| Scheme S1. Synthesis of substrate **8** | S3 |
| Scheme S2. Enzymatic synthesis of compounds **5** and **6** | S3 |
| Figure S1. Preparation of substrate **7** | S4 |
| Figure S2. $^{1}$H NMR spectrum of compound **7** | S5 |
| Figure S3. Anomeric region $^{13}$C -$^{1}$H HSQC spectrum (assignment **7**) | S5 |
| Figure S4. Proliferation and viability HL60 and U-2 OS cells | S6 |
| Figure S5. Terminal *N*-glycan epitopes LC-MS glycomics HL-60 cells | S7 |
| **2) Experimental procedures** | S8 |
| General procedures | S8 |
| Synthetic protocols and compound characterization | S8 |
| Kinetic studies | S18 |
| Cell culture | S18 |
| Reagents (antibodies and lectins) | S18 |
| Flow cytometry | S19 |
| Cell proliferation and viability | S19 |
| Sugar nucleotide analysis | S19 |
| Glycan profiling | S20 |
| **3) References** | S22 |
| **4) NMR spectra** | S23 |

### 1) Supplementary schemes and figures

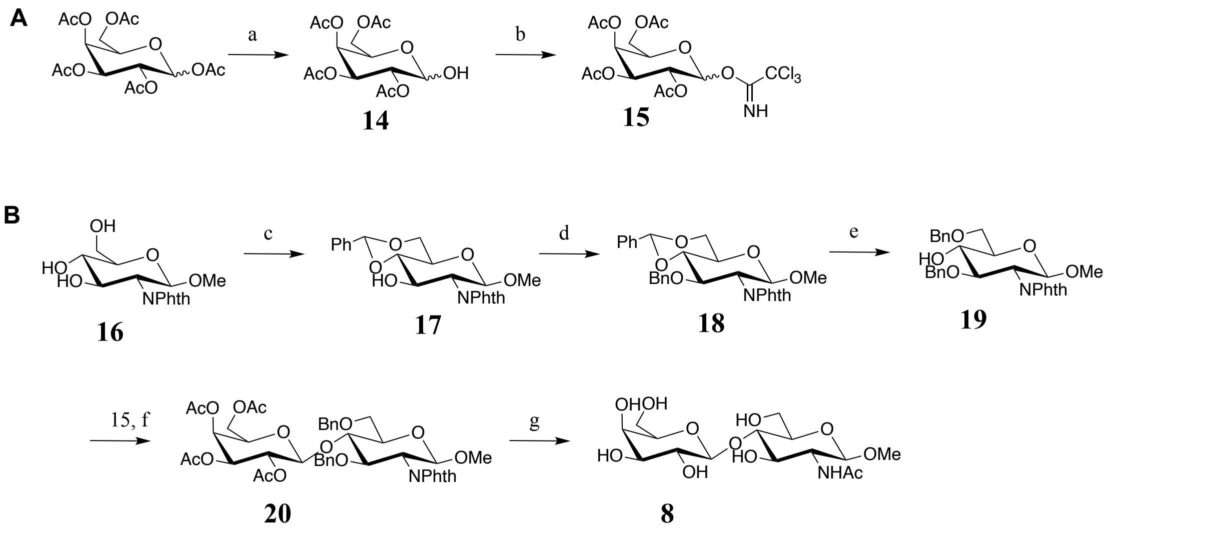


**Scheme S1.** Synthesis of substrate **8**. **A** Synthesis of building block **15**. a) Benzylamine, THF, 84%. b) Trichloroacetonitrile, Potassium carbonate, DCM, 77%. **B** Synthesis of LacNAc substrate **8**. c) Benzaldehyde dimethyl acetal, p-TsOH, $CH_3CN$, 95%. d) BnBr, NaH, DMF, 50%. e) Triethyl silane, trifluoroacetic acid, DCM, 44%. f) TMSOTf, DCM, 4Å ms, 55%. g) i. MeONa/MeOH, ii. ethylenediamine, EtOH, iii. MeOH, $Ac_2O$, iv. $H_2$, Pd/C, MeOH, $H_2O$, 95%.

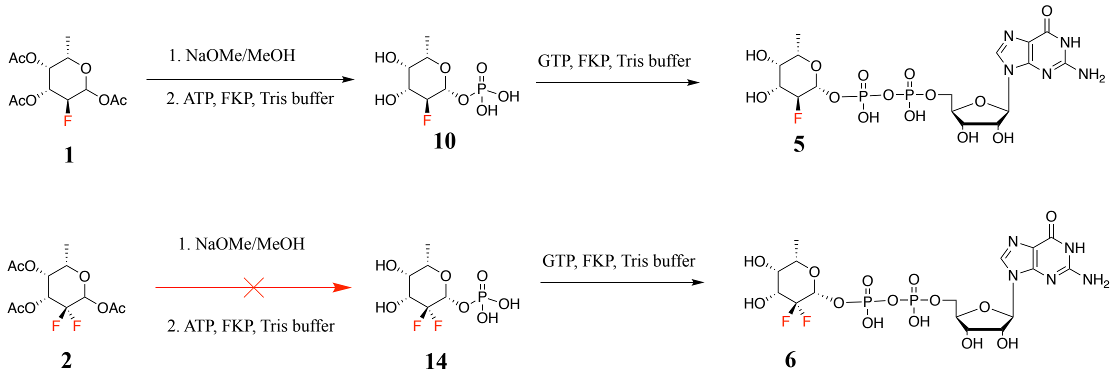


**Scheme S2.** Enzymatic synthesis of target fluorinated GDP-Fuc compounds (**5** and **6**).

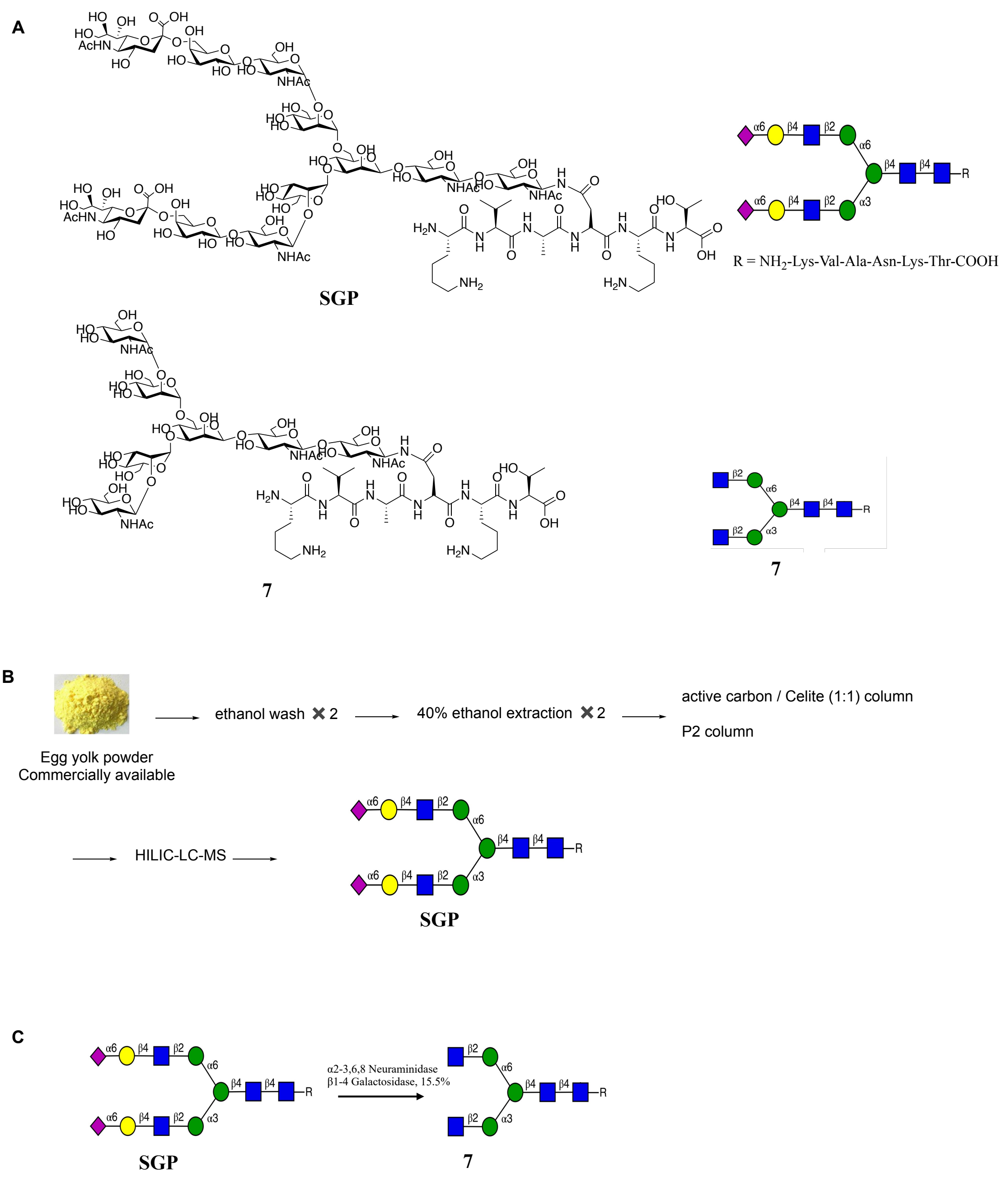


**Figure S1.** Preparation of substrate **7**.

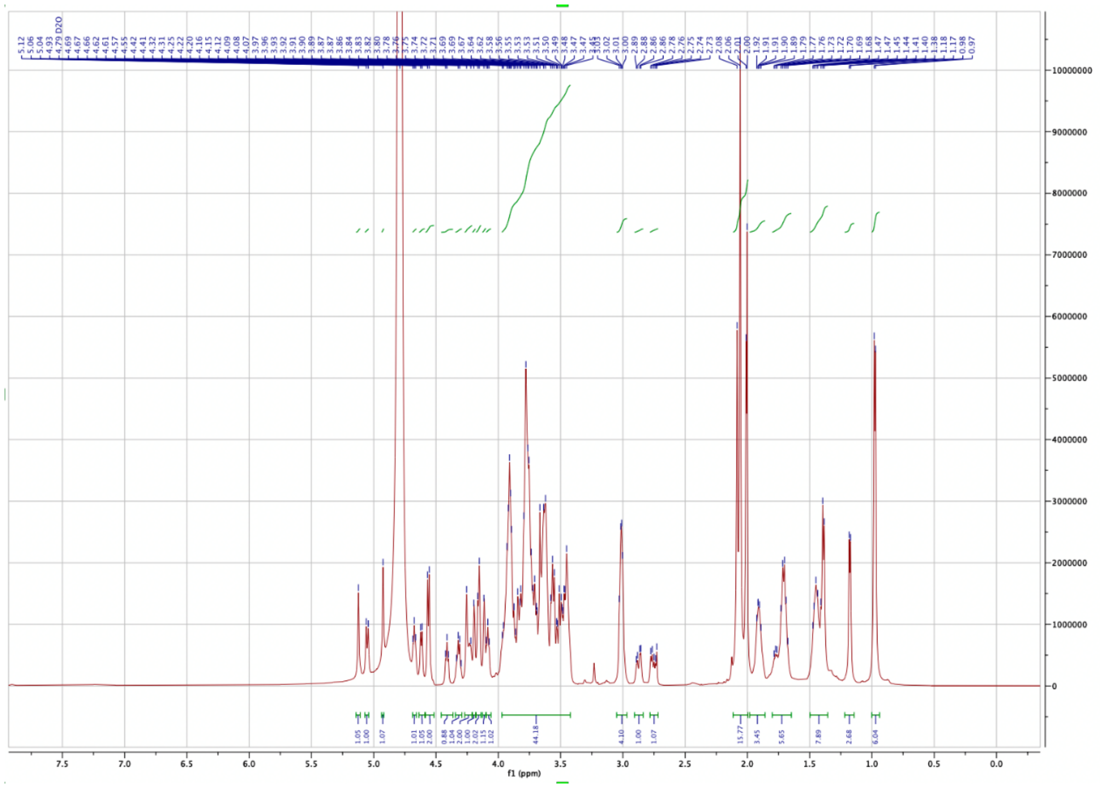


**Figure S2.** $^{1}$H NMR spectrum of compound **7** (600 MHz, deuterium oxide).

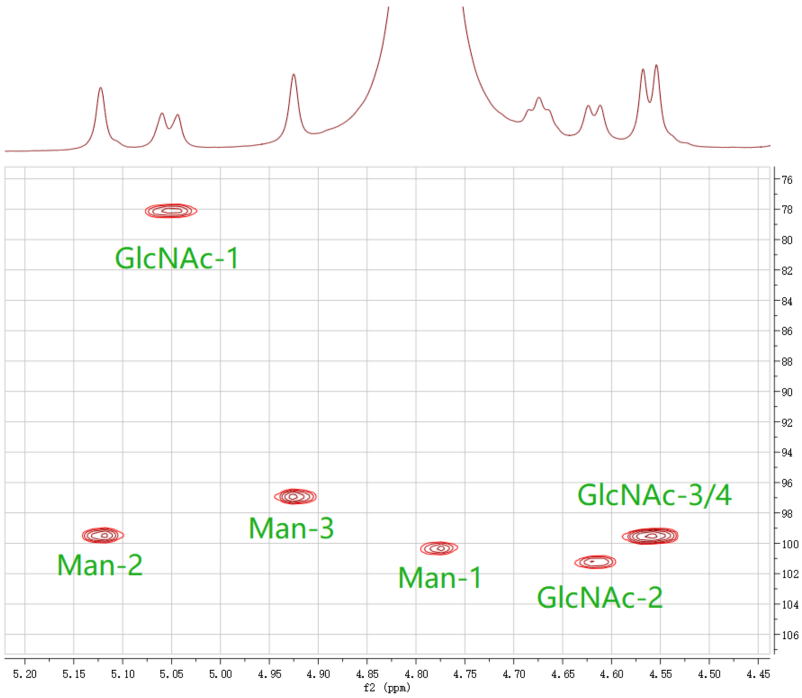


**Figure S3.** Anomeric region of the $^{13}$C -$^{1}$H HSQC spectrum (complete anomeric assignment of **7**).

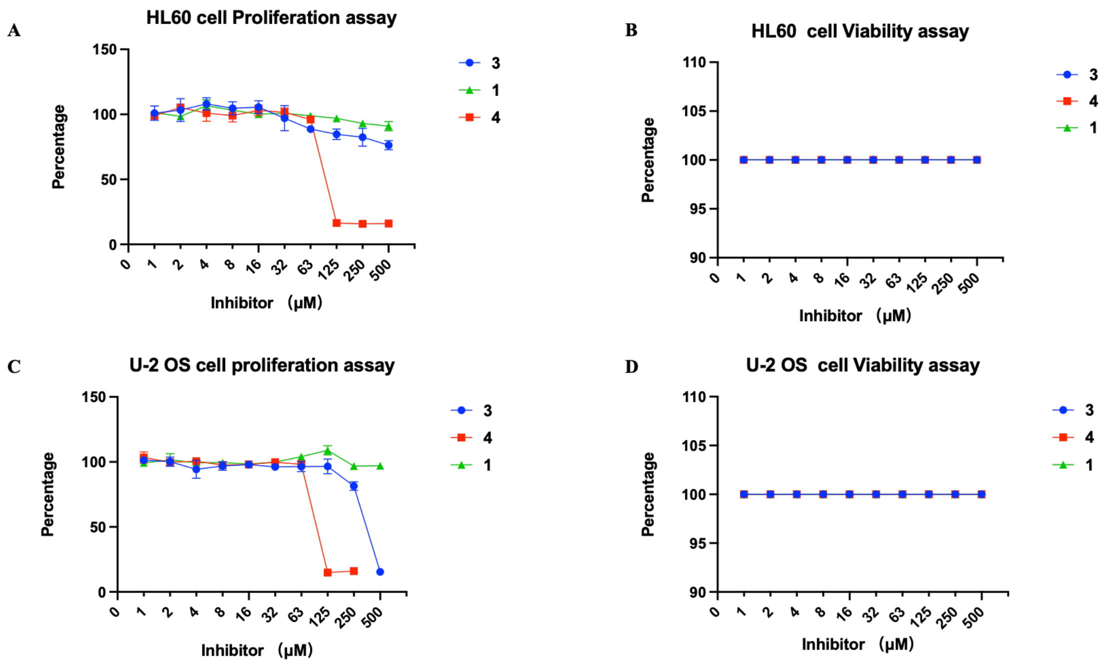


**Figure S4.** Proliferation and viability results with HL60 (**A, B**) and U-2 OS (**C, D**) cells.

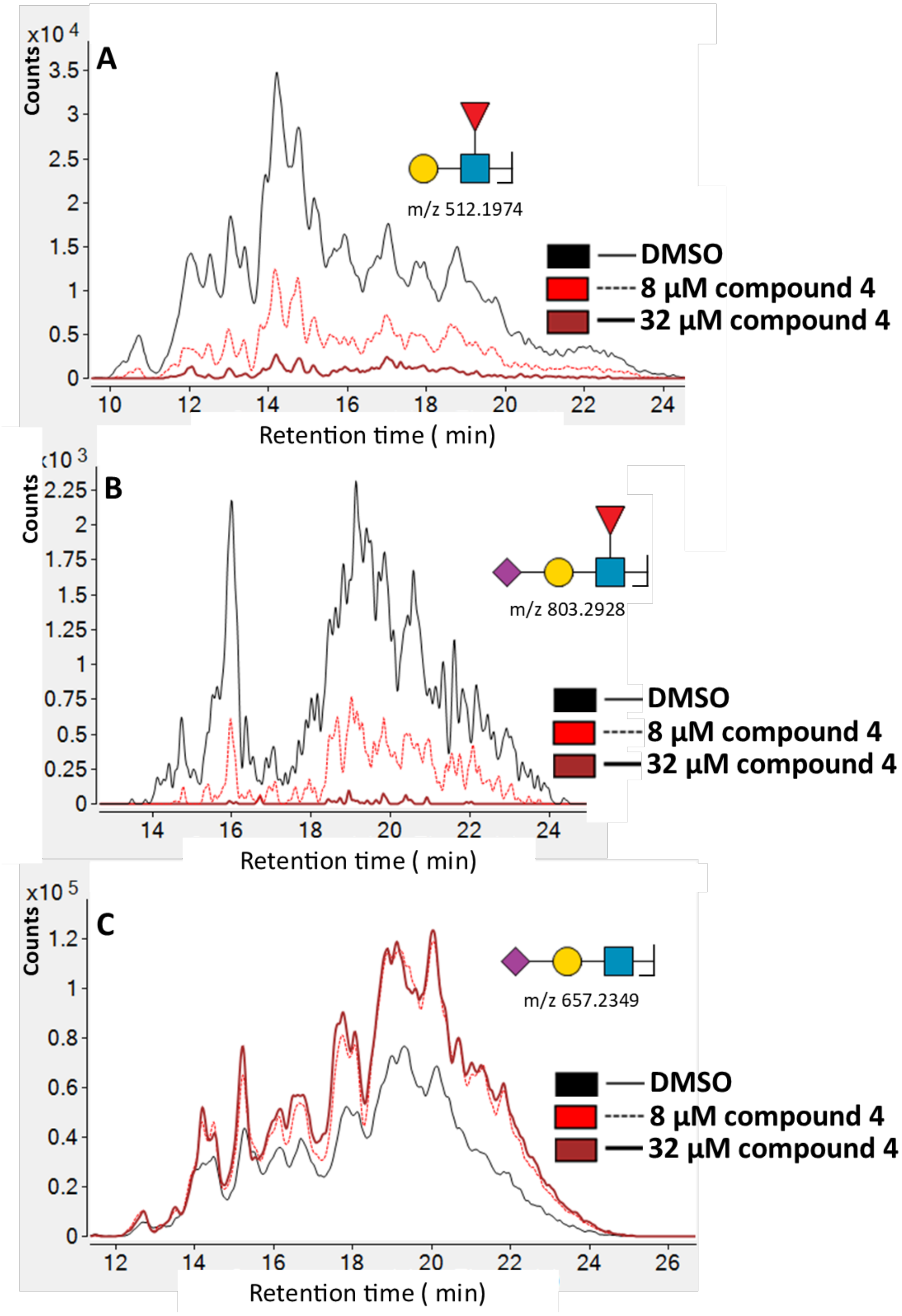


**Figure S5.** In-source fragments of terminal *N*-glycan epitopes from the LC-MS glycomic analysis of HL-60 cells incubated with DMSO control and 8 or 32 µM of compound **4**. Overlaid fragment B-ion extracted ion count overlays (5 ppm symmetrical expansion) are shown for (A) Fucosyl-LacNAc, (B) Sialyl Lewis$^{X}$ and (C) Sialyl LacNAc.

### 2) Experimental procedures

#### General procedures

All chemicals were purchased from Sigma-Aldrich, Fisher Scientific or Biosynth Carbosynth. NMR spectra were recorded with Agilent 400 or Bruker 600. 1H NMR data are presented in the order: Chemical shift, multiplicity (s = singlet, d = doublet, t = triplet, dd = doublet of doublets, m = multiplet) and coupling constants (J) are reported in Hertz (Hz). High-resolution mass spectrometry (HRMS) was recorded on an Agilent technologies 6560 Ion mobility Q-TOF. Column chromatography was performed on silica gel G60 (Silicycle, 60-200 μm, 60 Å), and size exclusion chromatography was performed on Bio-Gel P-2 (45-90 μm) by using 10-100 mM $NH_4HCO_3$ in Milli-Q $H_2O$ as eluent. Thin layer chromatography (TLC) analysis was conducted on Silica gel 60 F254 (EMD Chemicals Inc.) coated aluminum sheets and detected by using UV light (254 nm), staining by 5 % sulfuric acid in ethanol or p-anisaldehyde solution. Acid washed molecular sieves were flame activated under vacuum prior to reactions. Ion-exchange resin was purchased from Sigma-Aldrich. Before use, the $H^+$ resin (Amberlite IRC 120H, hydrogen form) was washed and activated, sequentially washed with Milli-Q $H_2O$, 1N NaOH, Milli-Q $H_2O$, 1N HCL and Milli-Q $H_2O$. MALDI-TOF MS analyses were performed on an Ultraflex II mass spectrometer (Bruker Daltonics, Bremen, Germany) equipped with a Smartbeam laser. Spectra were acquired in positive reflectron mode, using a method with a range of 700-3500 m/z. Samples were reconstituted in 50% ACN, 50% H2O and 1 μL was spotted onto the target plate and let dry under vacuum. 10 mg/mL solution of 2,5-dihydroxybenzoic acid matrix was dissolved in 50% ACN, 50% H2O, 0.1% TFA. The matrix solution (1 μL) was spotted onto the sample spot and dried at room temperature.

#### Synthetic protocols and compound characterization

**3,4-di-*O*-acetyl-fucal**

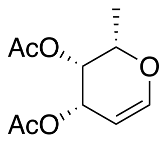


L-fucose (15 g, 90 mmol) was dissolved in acetic anhydride (75 mL) followed by the dropwise addition of $HClO_4$ (0.75 mL) at 0 °C. The reaction mixture was stirred at room temperature (RT) for 2 h, and then cooled (0 °C) and HBr (130 mL, 33% in AcOH) was added. After stirring at RT for 1 h, the reaction mixture was concentrated under reduced pressure. The resulting residue was dissolved in ethyl acetate and saturated $NaH_2PO_4$ (1:2, 390 mL), and zinc dust (66 g) was added. The mixture was stirred for 12 h at RT and then filtered. The filtrate was subsequently extracted with ethyl acetate, dried ($Na_2SO_4$) and filtered. The filtrate was concentrated under reduced pressure and the residue was purification by silica gel column chromatography affording fucal as colorless oil

(14.3 g, 73%).[1]. 1H NMR (600 MHz, Chloroform-*d*) δ 6.45 (dd, *J* = 6.3, 1.9 Hz, 1H, H-1), 5.56 (m, 1H, H-3), 5.27 (dt, *J* = 4.7, 1.3 Hz, 1H,H-4), 4.63 (dt, *J* = 6.3, 2.0 Hz, 1H, H-2), 4.20 (q, *J* = 6.2 Hz, 1H, H-5), 2.15 (s, 3H, $CH_3$-Ac), 2.00 (s, 3H, $CH_3$-Ac), 1.26 (d, *J* = 6.6 Hz, 3H, $CH_3$-Fuc). $^{13}$C NMR (151 MHz, Chloroform-*d*) δ 170.82, 170.52, 146.24, 98.39, 71.65, 66.38, 65.16, 20.98, 20.83, 16.65.

**2-Deoxy-2-fluoro-per-*O*-acetyl-fucose (1)**

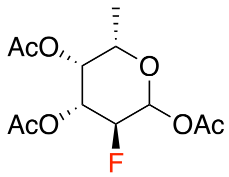


To a solution of Fucal (6 g, 28 mmol) in DMF/$H_2O$ (1:1, 100 mL), Selectfluor (16 g, 46 mmol) was added, and the mixture was stirred at 50 °C for 5 h. Then the mixture was diluted with ethyl acetate and washed with sat. aqueous $NaHCO_3$. The organic layer was dried ($Na_2SO_4$), filtered and the filtrate concentrated *in vacuo*. The resulting residue was dissolved in pyridine (50 mL) and acetic anhydride (50 mL). After stirring for 2 h at RT, the reaction mixture was concentrated in vacuo. Purification by silica gel column chromatography afforded 1 as a colorless oil (5.9 g, 73%, α: β = 1.4:1). $^1$H NMR (600 MHz, Chloroform-*d*) δ 6.42 (t, *J* = 4.1 Hz, 1.3H, αH-1), 5.77 (m, 1H, βH-1), 5.40 (m, 1.4H, αH-3), 5.36 (m, 1.3H, αH-4), 5.30 (m, 1H, βH-4), 5.16 (m, 1H, βH-3), 4.92 (dt, *J* = 10.2, 3.9 Hz, 0.7H, αH-2), 4.83 (dt, *J* = 10.2, 3.9 Hz, 0.7H, αH-2), 4.67 (ddd, *J* = 9.7, 8.0, 4.7 Hz, 0.5H, βH-2), 4.58 (ddd, *J* = 9.7, 8.0, 4.7 Hz, 0.5H, βH-2), 4.24 (m, 1.4H, αH-5), 3.98 (q, *J* = 6.5, 6.0 Hz, 1H, βH-5), 2.24 – 2.14 (m, 14H, α/β $CH_3$-Ac), 2.06 (m, 7H, α/β $CH_3$-Ac), 1.21 (dd, *J* = 6.5 Hz, 3H, β $CH_3$-Fuc), 1.14 (dd, *J* = 6.6, 4H, α $CH_3$-Fuc). $^{13}$C NMR (151 MHz, Chloroform-*d*) δ 170.45, 170.43, 170.24, 169.99, 169.23, 169.14, 91.75 (d, *J* = 24.3 Hz, βC-1), 89.35 (d, *J* = 22.3 Hz, αC-1), 86.93 (d, *J* = 188.0 Hz, βC-2), 84.37 (d, *J* = 190.7 Hz, αC-2), 71.49 (d, *J* = 18.4 Hz, βC-3), 71.21 (d, *J* = 7.7 Hz, αC-4), 70.72 (d, *J* = 8.2 Hz, βC-4), 70.44, 68.74 (d, *J* = 18.6 Hz, αC-3), 67.27, 21.06, 21.00, 20.81, 20.73, 20.67, 15.91, 15.88. ESI MS(m/z): $[M + Na]^+$ calcd for $C_{12}H_{17}FO_7Na$, 315.0856; found 315.0231.

**2-Deoxy-2-fluoro-3,4-di-*O*-acetyl-fucal (11)**

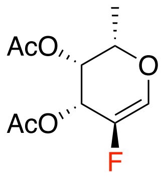


HBr (33% in AcOH, 10 mL) was added to compound **1** (6 g, 20 mmol) and after stirring for 0.5 h at RT, the mixture was concentrated *in vacuo*. The residue was dissolved in acetonitrile (185 mL), and triethylamine (11.1 mL, 84 mmol) was added. After stirring for 12 h at 82 °C, the reaction mixture was concentrated *in vacuo*. The residue was purified by silica gel chromatography to afford **11** (2.5 g, 52%) as colorless oil. $^1$H NMR (400 MHz, Chloroform-*d*) δ 6.74 (m, 1H, H-1), 5.87 (m, 1H, H-3), 5.30

(m, 1H, H-4), 4.17 (m, 1H, H-5), 2.17 (s, 3H, $CH_3$-Ac), 2.07 (s, 3H, $CH_3$-Ac), 1.29 (d, $J$ = 7.0 Hz, 3H, $CH_3$-Fuc). $^{13}$C NMR (101 MHz, Chloroform-*d*) δ 170.56, 170.29, 142.80 (d, $J_{C2-F}$ = 242.7 Hz), 133.08 (d, $J_{C1-F}$ = 39.7 Hz), 72.32, 66.78 (d, $J_{C4-F}$= 6.2 Hz), 63.48 (d, $J_{C3-F}$ = 20.2 Hz), 20.71, 16.02.

**2-Deoxy-2,2-di-fluoro-per-*O*-acetyl-fucose (2)**

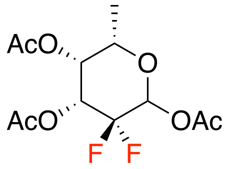


Compound **11** (50 mg, 0.2 mmol) was dissolved in $MeNO_2/H_2O$ (4:1, 1 mL). Selectfluor (114 mg, 0.3 mmol) was added, and the reaction mixture was placed in a microwave reactor and stirred for 5 min at 100 °C, after which it was allowed to cool to RT. The mixture was diluted with ethyl acetate and washed with water. The organic layer was dried ($Na_2SO_4$), filtered and the filtrate concentrated *in vacuo*. The residue was dissolved in a mixture of pyridine and acetic anhydride (1:1, 3 mL), and then stirred for 2 h at RT. It was then diluted with ethyl acetate and washed with brine and the organic layer was dried ($Na_2SO_4$), filtered and the filtrate concentrated *in vacuo*. The residue was purified by silica gel column chromatography to afford **2** (56 mg, 85%, α: β = 4:1) as colorless oil. $^1$H NMR (600 MHz, Chloroform-*d*) δ 6.20 (d, $J$ = 7.2 Hz, 4H, αH-1), 5.78 (d, $J$ = 15.6 Hz, 1H, βH-1), 5.42 (dt, $J$ = 23.1, 4.5 Hz, 4H, αH-3), 5.32 (d, $J$ = 4.3 Hz, 4H, αH-4), 5.29 (d, $J$ = 3.9 Hz, 1H, βH-4), 5.24 (dt, $J$ = 21.6, 4.8 Hz, 1H, βH-3), 4.32 (q, $J$ = 6.6 Hz, 4H, αH-5), 4.05 (dd, $J$ = 12.7, 6.5 Hz, 1H, βH-5), 2.18 (m, 30H, α/β $CH_3$-Ac), 2.11 (m, 15H, α/β $CH_3$-Ac), 1.28 (d, $J$ = 6.4 Hz, 3H, β $CH_3$-Fuc), 1.21 (d, $J$ = 6.5 Hz, 12H, α $CH_3$-Fuc). $^{13}$C NMR (151 MHz, Chloroform-*d*) δ 170.83, 170.76, 169.78, 169.57, 168.71, 168.18, 113.28, (t, $J_{C2-F}$ = 251.8 Hz, C-2), 90.01 (m, C-1), 89.74 (m, C-1), 70.85, 69.64 (d, $J$ = 7.6 Hz), 69.41 (d, $J$ = 6.6 Hz), 69.04 – 68.69 (m), 67.45, 66.39 (dd, $J$ = 21.6, 16.5 Hz), 20.92, 20.81, 20.66, 20.56, 20.50, 15.80. $^{19}$F NMR (565 MHz, Chloroform-*d*) δ -118.48 (ddd, $J$ = 256.2, 23.0, 7.4 Hz, $Fa_{ax}$), -120.76 (dt, $J$ = 256.2, 4.6 Hz, $Fa_{eq}$), -122.19 (dt, $J$ = 246.9, 4.9 Hz, $Fb_{eq}$), -133.27 (ddd, $J$ = 246.1, 21.6, 15.6 Hz, $Fb_{ax}$). ESI MS(m/z): $[M + Na]^+$ calcd for $C_{12}H_{16}F_2NaO_7$, 333.0762; found 333.0057

**2-Deoxy-2-fluoro-3,4-di-*O*-acetyl-β-1-(dibenzylphosphoryl)-L-fucopyranose (9)**

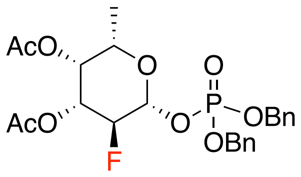


HBr (33% in AcOH, 1 mL) was added to compound **1** (200 mg, 0.68 mmol), and after stirring the resulting reaction mixture for 0.5 h at RT, it was concentrated *in vacuo*. The residue was dissolved in $CH_3CN$ (5 mL), 3Å molecular sieves (500 mg) were added. The mixture was stirred at RT for 10 min and then silver carbonate (375 mg, 1.36 mmol) and dibenzyl phosphate (375 mg, 1.36 mmol) were added. The reaction mixture was stirred at RT, under and atmosphere of nitrogen in the dark. After 12 h, the reaction mixture was diluted with ethyl acetate and then filtrated through celite, and the filtrate concentrated *in vacuo*. The residue was purified by silica gel chromatography affording **9** (244 mg, 70%) as colorless oil. $^{1}$H NMR (600 MHz, Chloroform-*d*) δ 7.44 – 7.31 (m, 10H, Ar), 5.37 (td, *J* = 7.5, 3.9 Hz, 1H, H-1), 5.28 (t, *J* = 3.2 Hz, 1H, H-4), 5.18 – 5.03 (m, 5H, $CH_2$, H-3), 4.65 (dd, *J* = 9.9, 7.7 Hz, 0.5H, H-2), 4.56 (dd, *J* = 9.9, 7.7 Hz, 0.5H, H-2), 3.92 (q, *J* = 6.4 Hz, 1H, H-5), 2.17 (s, 3H, $CH_3$-Ac), 2.06 (s, 3H, $CH_3$-Ac), 1.20 (d, *J* = 6.4 Hz, 3H, $CH_3$-Fuc). ESI MS(m/z): [M + Na]+ calcd for $C_{24}H_{28}FO_9PNa$, 533.1353; found 533.0648

**2-Deoxy-2,2-di-fluoro-3,4-di-*O*-acetyl-β-1-(dibenzylphosphoryl)-L-fucopyranose (12)**

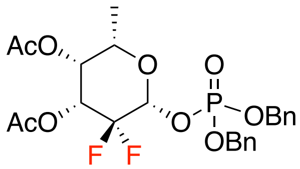


Compound **11** (50 mg, 0.2 mmol) was dissolved in $MeNO_2/H_2O$ (4:1, 1 mL), Selectfluor (114 mg, 0.3 mmol) was added, and the resulting reaction mixture was placed in microwave reactor and stirred for 5 min at 100 °C, after which it was allowed to cool down to RT The mixture was diluted with ethyl acetate and washed with water, dried ($Na_2SO_4$), filtered and the filtrate concentrated *in vacuo*. The residue was dissolved in 1,2-dichloroethane (2 mL), caesium carbonate (169 mg, 0.52 mmol) and tetrabenzyl pyrophosphate (173.4 mg, 0.32 mmol) were subsequently added. The reaction mixture was stirred at 40 °C for 3 h, and allowed then to cool down to RT, diluted with DCM and washed with brine. The organic layer was dried ($Na_2SO_4$) and filtered and the filtrate concentrated *in vacuo*. The residue was purified by silica gel column chromatography to afford **12** (31.8 mg, 28%) as colorless oil. $^{1}$H NMR (600 MHz, Chloroform-*d*) δ 7.35 (m, 10H, Ar), 5.41 (dd, $J_{H1\text{-}Fax}$ = 14.3 Hz, $J_{H1\text{-}P}$=8.2 Hz, 1H, H-1), 5.28 (d, *J* = 3.5 Hz, 1H, H-4), 5.20 (dt, $J_{H3\text{-}Fax}$ = 20.7 Hz, *J* =3.9 Hz, 1H, H-3), 5.16 – 5.07 (m, 4H, $CH_2$Ar), 4.01 (q, *J* = 6.3 Hz, 1H, H-5), 2.19 (s, 3H, $CH_3$-Ac), 2.12 (s, 3H, $CH_3$-Ac), 1.26 (d, *J* = 6.4 Hz, 3H, $CH_3$-Fuc). $^{13}$C NMR (151 MHz, Chloroform-*d*) δ 170.73, 169.51, 135.45, 135.40, 135.27, 135.22, 128.85, 128.77, 128.73, 128.68, 128.18, 128.16, 113.11 (t, $J_{C2\text{-}F}$ = 251.1 Hz, C-2), 94.04 (dd, $J_{C1\text{-}F}$ = 29.0, 19.0Hz, C-1), 70.77, 70.05 (dd, *J* = 18.0, 5.6 Hz), 69.24 (d, *J* = 6.7 Hz), 68.66 (dd, *J* = 21.0, 16.6 Hz), 20.67, 20.49, 15.81. $^{13}$C NMR (151 MHz, Chloroform-*d*) δ 170.73, 169.51, 135.45, 135.40, 135.27, 135.22, 128.85, 128.77, 128.73, 128.68, 128.18, 128.16, 113.11 (t, $J_{C2\text{-}F}$ = 251.1 Hz), 94.04 (dd, $J_{C1\text{-}F}$ = 29.0, 19.0 Hz), 70.77, 70.11 (d, $J_{CH2Ar\text{-}F}$= 5.6 Hz), 69.99 (d, $J_{CH2Ar\text{-}F}$ = 5.5 Hz), 69.24 (d, $J_{C4\text{-}F}$ = 6.7 Hz), 68.66 (dd, $J_{C3\text{-}F}$ = 21.0, 16.6 Hz), 20.67, 20.49, 15.81. $^{19}$F NMR (565 MHz, Chloroform-*d*) δ -121.77 (dt, $J_{C2\text{-}F}$ = 250.6 Hz,

*J*=4.5Hz, $F_{eq}$), -134.57 (ddd, $J_{C2-F}$ = 250.6 Hz, $J_{H3-Fax}$ = 20.7, $J_{H1-Fax}$ = 14.2 Hz, $F_{ax}$). ESI MS(m/z): [M + Na+] calcd for $C_{24}H_{27}F_2O_9PNa$, 551.1258; found 551.0592.

**2-deoxy-2,2-di-fluoro-β-fucopyranosyl phosphate (13)**

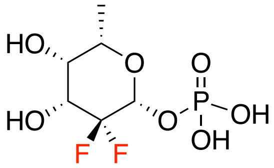


The mixture of compound **12** (55 mg, 0.1 mmol) in MeOH (2 mL) and 10% activated Pd/C (40 mg) was placed under an atmosphere of $H_2$. After 16 h, the reaction mixture was filtered, and the filtrate was concentrated *in vacuo*. The residue was dissolved in a mixture of MeOH:$H_2O$:$Et_3N$ (7:3:1, 3.3 mL) and stirred for 6 h at RT. The mixture was then concentrated in vacuo and purified by silica gel chromatography to afford **13** (24 mg, 88%) as white solid. $^1$H NMR (400 MHz, Deuterium Oxide) δ 5.21 (dd, $J_{H1-Fax}$ = 15.7 Hz, $J_{H1-P}$ = 9.2 Hz, 1H, H-1), 4.15 (dt, $J_{H3-Fax}$ = 23.2 Hz, $J_{H3-Feq}$ = 4.6 Hz, 1H, H-3), 3.99 (q, *J* = 6.7 Hz, 1H, H-5), 3.92 (t, *J* = 4.4 Hz, 1H, H-4), 3.24 (q, *J* = 7.3 Hz, 7H, $CH_2$-$Et_3N$), 1.33 (q, *J* = 7.3 Hz, 14H, $CH_3$-$Et_3N$, $CH_3$-Fuc). $^{13}$C NMR (151 MHz, Deuterium Oxide) δ 115,89 ( t, $J_{C2-F}$ = 249.1 Hz), 92.85 (dd, $J_{C1-F}$ = 29.2, 18.8 Hz), 71.61, 70.78 (d, *J* = 7.5 Hz, C-4), 69.18 (t, *J* = 18.4 Hz, C-3), 46.54, 15.10, 8.15. $^{19}$F NMR (376 MHz, Chloroform-*d*) δ -118.95 (dt, $J_{C2-F}$ = 249.6 Hz, $J_{H3-Feq}$ = 4.6Hz, $F_{eq}$), -133.94 (ddd, $J_{C2-F}$ = 249.0 Hz, $J_{H3-Fax}$ = 23.4 Hz, $J_{H1-Fax}$ = 15.6 Hz). ESI MS(m/z): $[M - H]^-$ calcd for $C_6H_{10}F_2O_7P$, 263.0132; found 263.0212.

**2-deoxy-2-fluoro-β-fucopyranosyl phosphate (10)**

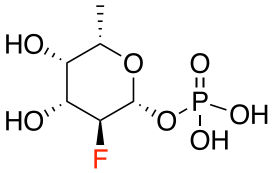


$^1$H NMR (400 MHz, Deuterium Oxide) δ 5.10 (td, *J* = 8.0, 3.7 Hz, 1H, H-1), 4.43 (dd, *J* = 9.6, 7.7 Hz, 0.5H, H-2), 4.30 (dd, *J* = 9.5, 7.7 Hz, 0.5H, H-2), 3.98 (m, 1H, H-3), 3.88 (q, *J* = 6.5 Hz, 1H, H-5), 3.83 (t, *J* = 3.3 Hz, 1H, H-4), 3.21 (q, *J* = 7.3 Hz, 9H, $CH_2$-$Et_3N$), 1.34 – 1.24 (m, 15H, $CH_3$-Fuc, $CH_3$-$Et_3N$). $^{13}$C NMR (101 MHz, Deuterium Oxide) δ 95.12 (d, *J* = 23.9 Hz), 90.80, 71.71 (d, *J* = 9.0 Hz), 71.50, 71.21 (d, *J* = 17.0 Hz), 46.60, 15.14, 8.15. ESI MS(m/z): $[2M - H]^-$ calcd for $C_{12}H_{23}F_2O_{14}P_2$, 491.0537; found 491.0350.

**GDP-β-2,2-di-fluoro-fucose (6)**

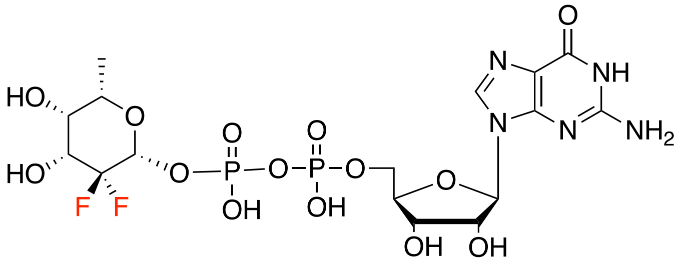


Compound **13** (8.5 mg, 0.03 mmol) and GMP-morpholidate (50 mg, 0.07mmol) were co-evaporated with pyridine and subsequently dried. 1H-Tetrazole (8 mg, 0.1 mmol) and pyridine were added to the mixture, and the solution was stirred at RT under an atmosphere od $N_2$. After 2 days, the mixture was diluted with water and applied to a Bio-Gel P2 column that was eluted with 50 mM $NH_4HCO_3$ solution to isolate compound **6** (3.7 mg, 18%) as white amorphous solid. $^1$H NMR (600 MHz, Deuterium Oxide) δ 8.13 (s, 1H), 5.91 (d, *J* = 6.2 Hz, 1H, $H_{Guo}$-1), 5.25 (dd, *J* = 15.5, 9.4 Hz, 1H, $H_{Fuc}$-1), 4.77 (m, 1H, $H_{Guo}$-2), 4.50 (t, *J* = 3.7 Hz, 1H, $H_{Guo}$-3), 4.33 (m, 1H, $H_{Guo}$-4), 4.19 (t, *J* = 4.3 Hz, 2H, $H_{Guo}$-5), 4.04 (dt, *J* = 23.4, 5.2 Hz, 1H, $H_{Fuc}$-3), 3.88 (q, *J* = 6.5 Hz, 1H, $H_{Fuc}$-5), 3.82 (d, *J* = 4.0 Hz, 1H, $H_{Fuc}$-4), 1.24 (d, *J* = 6.5 Hz, 3H, $CH_3$-Fuc). $^{13}$C NMR (101 MHz, Deuterium Oxide) δ 158.70, 153.87, 151.64, 137.49, 115.74 (t, $J_{C2-F}$ = 250.1 Hz, $C_{Fuc}$-2), 92.97 (dd, $J_{C1-F}$ = 29.4, 19.7 Hz, $C_{Fuc}$-1), 86.77 ($C_{Guo}$-1), 83.83 ($C_{Guo}$-4), 83.73, 73.54 ($C_{Guo}$-2), 71.90 ($C_{Fuc}$-5), 70.65 (d, *J* = 7.5 Hz, $C_{Fuc}$-4), 70.35 ($C_{Guo}$-3), 69.07 (t, *J* = 18.5 Hz, $C_{Fuc}$-3), 65.24 (d, *J* = 5.5 Hz, $C_{Guo}$-5), 15.01. ESI MS(m/z): [M - H]$^-$ calcd for $C_{16}H_{22}F_2N_5O_{14}P_2$, 608.0612; found 608.0384.

**GDP-β-2-fluoro-fucose (5)**

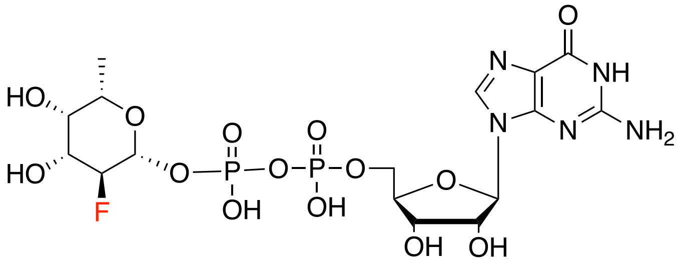


$^1$H NMR (400 MHz, Deuterium Oxide) δ 8.12 (s, 1H), 5.94 (d, *J* = 6.3 Hz, 1H, $H_{Guo}$-1), 5.18 (td, *J* = 7.9, 3.5 Hz, 1H, $H_{Fuc}$-1), 4.87 (m, 1H, $H_{Guo}$-2), 4.54 (dd, *J* = 5.2, 3.1 Hz, 1H, $H_{Guo}$-3), 4.44 (dd, *J* = 9.6, 7.5 Hz, 0.5H, $H_{Fuc}$-2), 4.36(m, 1H, $H_{Guo}$-4), 4.31 (dd, *J* = 9.5, 7.5 Hz, 0.5H, $H_{Fuc}$-2), 4.22 (dd, *J* = 5.4, 3.5 Hz, 2H, $H_{Guo}$-5), 3.92 (ddd, *J* = 14.4, 9.6, 3.6 Hz, 1H, $H_{Fuc}$-3), 3.82 (m, 1H, $H_{Fuc}$-5), 3.78 (t, *J* = 3.5 Hz, 1H, $H_{Fuc}$-4), 1.23 (d, *J* = 6.4 Hz, 3H, $CH_3$-Fuc). ESI MS(m/z): [M - H]$^-$ calcd for $C_{16}H_{23}FN_5O_{14}P_2$, 590.0706; found 590.0812.

**2-Deoxy-2,2-di-fluoro-3,4-di-*O*-acetyl-β-1-(dipivaloyloxymethylphosphoryl)-L-fucopyranose (4)**

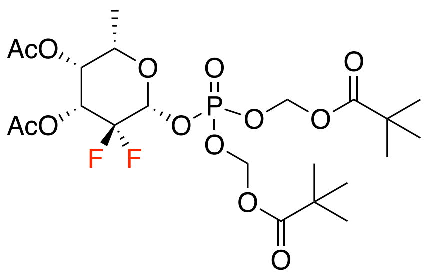

Compound **12** (55 mg, 0.1 mmol) and 10% activated Pd/C (30 mg) MeOH (2 mL) was placed under an $H_2$ atmosphere. After 12 h, the mixture was filtered, and the filtrate was concentrated *in vacuo*. The residue was dissolved in acetonitrile and $Ag_2CO_3$ (177 mg, 0.6 mmol) was added. The reaction mixture was stirred at RT for 2 h, and then pivaloyloxymethyl chloride (200 μL, 1.5 mmol) was added. After stirring for an additional 18 h at 60 C°, the reaction was cooled to RT, diluted with ethyl acetate and washed with brine. The organic layer was dried ($Na_2SO_4$), filtered, and the filtrate concentrated *in vacuo*. The residue was purified by silica gel column chromatography to afford **4** (38 mg, 66.6%) as colorless oil. $^1$H NMR (600 MHz, Chloroform-*d*) δ 5.74 – 5.63 (m, 4H, $CH_2$-POM), 5.40 (dd, *J* = 14.0, 7.6 Hz, 1H, H-1), 5.27 (d, *J* = 3.8 Hz, 1H, H-4), 5.20 (dt, *J* = 21.5, 4.4 Hz, 1H, H-3), 4.12 – 3.95 (m, 1H, H-5), 2.17 (s, 3H, $CH_3$-Ac), 2.12 (s, 3H, $CH_3$-Ac) 1.29 (d, *J* = 6.4 Hz, 3H, $CH_3$-Fuc), 1.24 (m, 18H, $C(CH_3)_3$-POM). $^{13}$C NMR (151 MHz, Chloroform-*d*) δ 176.59, 170.55, 169.34, 94.07 (m, C-1), 82.90 ($CH_2$), 70.80 (C-5), 69.02 (C-4), 68.64 – 68.26 (m, C-3), 38.73, 26.79, 26.74, 20.49, 20.34, 15.66. $^{19}$F NMR (565 MHz, Chloroform-*d*) δ -121.87 (d, *J* = 246.1 Hz, $F_{eq}$), -134.37 (ddd, *J* = 245.8, 21.8, 14.2 Hz, $F_{ax}$). ESI MS(m/z): $[M + Na]^+$ calcd for $C_{22}H_{35}F_2O_{13}PNa$, 599.1681; found 599.0009.

**2-Deoxy-2-fluoro-3,4-di-*O*-acetyl-β-1-(dipivaloyloxymethylphosphoryl)-L-fucopyranose (13)**

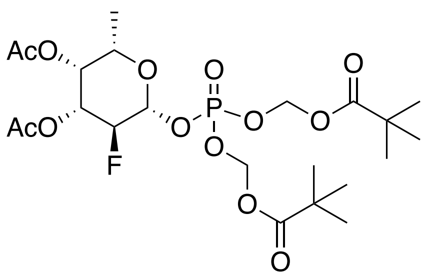

$^1$H NMR (600 MHz, Chloroform-*d*) δ 5.73 – 5.62 (m, 4H, $CH_2$-POM), 5.36 (m, 1H, H-1), 5.27 (m, 1H, H-4), 5.11 (ddd, *J* = 13.1, 9.9, 3.6 Hz, 1H, H-3), 4.61 (dd, *J* = 9.9, 7.6 Hz, 0.5H, H-2), 4.52 (dd, *J* = 9.9, 7.6 Hz, 0.5H, H-2), 3.95 (m, 1H, H-5), 2.16 (s, 3H, $CH_3$-Ac), 2.05 (s, 3H, $CH_3$-Ac), 1.23 (m, 21H, $CH_3$-Fuc, $C(CH_3)_3$-POM). $^{13}$C NMR (151 MHz, Chloroform-*d*) δ 176.73, 170.39, 169.95, 96.77 (dd, *J* = 24.2, 4.6 Hz, C-1), 87.72 (dd, *J* = 189.5, 9.2 Hz, C-2), 82.92, 71.02 (d, *J* = 18.6 Hz, C-3), 70.62 (C-5), 70.42 (d, *J* = 8.3 Hz, C-3), 38.86, 26.93, 26.90, 20.70, 20.66, 15.88. $^{19}$F NMR (565 MHz, Chloroform-*d*) δ -207.86 (dd,

$J$ = 51.8, 12.8 Hz). ESI MS(m/z): $[M + Na]^+$ calcd for $C_{22}H_{36}FO_{13}PNa$, 581.1775; found 581.0457.

**Methyl 4,6-Benzylidene-2-deoxy-2-N-phthalimido-β-D-glucopyranoside (17)**

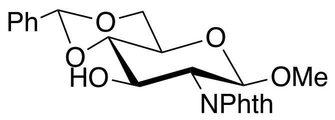

Compound **16** (1.3 g, 4 mmol) was dissolved in $CH_3CN$ (10 mL), Benzaldehyde dimethyl acetal (0.4 mL, 6 mmol) and p-TsOH·$H_2O$ (38 mg, 0.2 mmol) were then added. The reaction mixture was stirred at RT for 4 h. Then triethylamine was added to quench the reaction. The solution was dried and chromatographed to give compound **17** (1.6 g, 100%) as white solid. $^1$H NMR (600 MHz, Chloroform-*d*) δ 7.86 (m, 2H, Ar), 7.73 (m, 2H, Ar), 7.50 (m, 2H, Ar), 7.38 (m, 3H, Ar), 5.57 (d, $J$ = 5.5 Hz, 1H, CHPh), 5.20 (dd, $J$ = 8.5, 5.7 Hz, 1H, H-1), 4.66 – 4.59 (m, 1H, H-3), 4.41 (dt, $J$ = 10.7, 5.2 Hz, 1H, H-6a), 4.27 – 4.19 (m, 1H, H-2), 3.84 (td, $J$ = 10.1, 5.3 Hz, 1H, H-6b), 3.69 – 3.57 (m, 2H, H-4, H-5), 3.44 (d, $J$ = 5.9 Hz, 3H, OMe). $^{13}$C NMR (151 MHz, Chloroform-*d*) δ 137.08, 134.28, 131.86, 129.54, 128.55, 126.44, 123.71, 102.12, 99.97, 82.43, 68.85, 68.77, 66.25, 57.26, 56.56.

**Methyl 4,6-benzylidene-3-O-benzyl-2-deoxy-2-N-Phthalimido-β-D-glucopyranoside (18)**

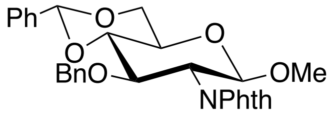

Compound **17** (1.6 g, 4 mmol) was dissolved in DMF (20 mL) and cooled to 0 °C. Then, sodium hydride (60% dispersion in mineral oil, 0.4 g, 10 mmol) was added in portions. After stirring for 20 min, benzyl bromide (2 mL, 16 mmol) was added dropwise. Then the reaction mixture was allowed to warm back to RT and stirred for 1 h. Afterward, MeOH was added to quench the reaction. The reaction mixture was diluted with ethyl acetate and washed with brine. The organic layer was collected and dried. Purification by silica gel chromatography afforded **18** (1.1 g, 50%) as white solid. $^1$H NMR (600 MHz, Chloroform-*d*) δ 7.71 (m, 3H, Ar), 7.53 (m, 2H, Ar), 7.43 – 7.33 (m, 4H, Ar), 7.03 – 6.97 (m, 2H, Ar), 6.93 (m, 1H, Ar), 6.88 (m, 2H, Ar), 5.63 (s, 1H, CHPh), 5.13 (d, $J$ = 8.5 Hz, 1H, H-1), 4.80 (d, $J$ = 12.3 Hz, 1H, $CH_2$-Bn), 4.50 (d, $J$ = 12.4 Hz, 1H, $CH_2$-Bn), 4.42 (ddd, $J$ = 10.4, 6.8, 1.9 Hz, 2H, H-6a, H-3), 4.21 (dd, $J$ = 10.4, 8.5 Hz, 1H, H-2), 3.87 (t, $J$ = 10.3 Hz, 1H, H-6b), 3.82 (t, $J$ = 9.2 Hz, 1H, H-4), 3.65 (td, $J$ = 9.8, 5.0 Hz, 1H, H-5), 3.40 (s, 3H, OMe). $^{13}$C NMR (151 MHz, Chloroform-*d*) δ 138.02, 137.45, 133.98, 131.80, 129.18, 128.45, 128.17, 127.53, 126.19, 123.50, 101.46, 99.97, 83.27, 74.70, 74.23, 68.94, 66.18, 57.16, 55.83.

**Methyl 3,6-O-dibenzyl-2-deoxy-2-N-Phthalimido-β-D-glucopyranoside (19)**

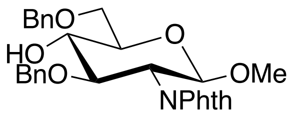

Compound **18** (1 g, 2 mmol) was dissolved in DCM (10 mL) and cooled to 0 °C. Triethyl silane (3.3 mL, 20 mmol) and trifluoroacetic acid (1.5 mL, 20 mmol) were added sequentially. The reaction mixture was stirred at 0 °C for 1 h. Next, the solution was washed with water, sat. aq. $NaHCO_3$ and brine. The organic layer was collected, dried ($Na_2SO_4$,), filtered and the filtrate concentrated, and the residue purified by silica column chromatography to give compound **19** (0.46 g, 44%) as colorless oil. $^1$H NMR (600 MHz, Chloroform-*d*) δ 7.69 (m, 4H, Ar), 7.42 – 7.29 (m, 5H, Ar), 7.09 – 7.02 (m, 2H, Ar), 7.00 – 6.90 (m, 3H, Ar), 5.07 (d, *J* = 7.8 Hz, 1H, H-1), 4.74 (d, *J* = 12.2 Hz, 1H, $CH_2$-Bn), 4.66 (d, *J* = 11.9 Hz, 1H, $CH_2$-Bn), 4.60 (d, *J* = 11.9 Hz, 1H, $CH_2$-Bn), 4.53 (d, *J* = 12.2 Hz, 1H, $CH_2$-Bn), 4.23 (dd, *J* = 10.7, 8.5 Hz, 1H, H-3), 4.14 (dd, *J* = 10.8, 8.4 Hz, 1H, H-2), 3.87 – 3.78 (m, 3H, H-6, H-4), 3.65 (dt, *J* = 9.8, 5.0 Hz, 1H, H-5), 3.38 (s, 3H, OMe), 2.89 (d, *J* = 2.5 Hz, 1H, OH).

**Methyl [peracetyl-β-D-galactopyranosyl]-(1→4)-3,6-O-dibenzyl-2-deoxy-2-N-Phthalimido-β-D-glucopyranoside (20)**

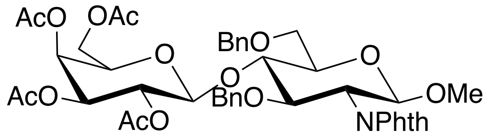

Compound **19** (0.46 g, 1 mmol) and 17 (0.67 g, 1.4 mmol) were dissolved in DCM (5 mL), and 4 Å molecular sieve (500 mg) was added. The reaction mixture was stirred at RT for 10 min and then cooled (-20 °C). Trimethylsilyl trifluoromethanesulfonate (33 µL, 0.2 mmol) was added and stirring was continued for 15 min. Next, the reaction was quenched with triethylamine and the mixture filtered through a pad of celite. The solution was concentrated in vacuo and the residue purification by silica gel column chromatography afforded **20** (0.5 g, 55%) as colorless oil. $^1$H NMR (600 MHz, Chloroform-*d*) δ 7.78 – 7.59 (m, 4H, Ar), 7.41 (d, *J* = 6.1 Hz, 4H, Ar), 7.34 (m, 1H, Ar), 7.01 (d, *J* = 6.5 Hz, 2H, Ar), 6.87 (m, 3H, Ar), 5.26 (d, *J* = 3.5 Hz, 1H, $H_{Gal}$-4), 5.14 (dd, *J* = 10.4, 8.0 Hz, 1H, $H_{Gal}$-2), 5.01 (d, *J* = 8.5 Hz, 1H, $H_{Glcl}$-1), 4.86 – 4.81 (m, 2H, $H_{Gal}$-3, $CH_2$-Bn), 4.79 (d, *J* = 12.3 Hz, 1H, $CH_2$-Bn), 4.58 (d, *J* = 8.0 Hz, 1H, $H_{Gal}$-1), 4.50 (d, *J* = 12.1 Hz, 1H, $CH_2$-Bn), 4.42 (d, *J* = 12.3 Hz, 1H, $CH_2$-Bn), 4.24 (dd, *J* = 10.7, 8.5 Hz, 1H, $H_{Glcl}$-3), 4.13 (dd, *J* = 10.8, 8.5 Hz, 1H, $H_{Glcl}$-2), 4.10 – 4.05 (m, 1H, $H_{Glcl}$-4), 4.00 – 3.91 (m, 2H, $H_{Gal}$-6), 3.82 – 3.76 (m, 2H, $H_{Glcl}$-6), 3.63 (t, *J* = 6.9 Hz, 1H, $H_{Gal}$-5), 3.53 (dt, *J* = 9.9, 2.5 Hz, 1H, $H_{Glcl}$-5), 3.38 (s, 3H, OMe), 2.06 (s, 3H, $CH_3$-Ac), 2.02 (s, 6H, $CH_3$-Ac), 1.97 (s, 3H, $CH_3$-Ac). $^{13}$C NMR (151 MHz, Chloroform-*d*) δ 170.48, 170.39, 170.21, 169.35, 138.67, 138.00, 128.77, 128.25, 128.22, 128.04, 127.96, 127.22, 100.46, 99.40, 78.15, 76.82, 74.91, 74.57, 73.79, 71.13, 70.57, 69.65,

67.59, 67.03, 60.85, 56.82, 55.64, 20.96, 20.83, 20.77, 20.73. ESI MS (m/z): [M + Na]+ calcd for $C_{43}H_{47}NO_{16}Na$, 856.2793; found 856.4816.

**Methyl [β-D-galactopyranosyl]-(1→4)- 2-deoxy-2-acetamido-β-D-glucopyranoside (8)**

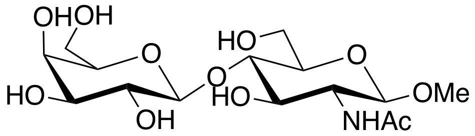


Compound **20** (0.5 g, 0.6 mmol) was dissolved in MeOH (10 mL), and freshly prepared sodium methoxide was added dropwise until the pH reached approximately 10. The reaction mixture was stirred at RT for 30 min, and then it was neutralized with $H^+$ resin. The solution was filtered through celite and concentrated under vacuum. The resulting residue was dissolved in ethanol (40 mL) and ethylenediamine (1.3 mL, 20 mmol) was added, the reaction mixture was refluxed for 5 h. Afterward, the solution was concentrated in vacuo. The residue was dissolved in methanol (10 mL) and acetic anhydride (0.5 mL, 5 mmol) was added. The reaction mixture was stirred at RT for 6 h. Next, the reaction mixture was concentrated in vacuo. The resulting residue was dissolved in a mixture of methanol (5 mL) and water (5 mL) and hydrogenated under an $H_2$ atmosphere with 10% activated $Pd(OH)_2/C$ (30 mg) as the catalyst for 3 h. The solution was filtered through a pad of celite and concentrated under vacuo. The residie was purified by silica gel column chromatography to afford **8** (252 mg, 95%) as white powder. $^1$H NMR (600 MHz, Methanol-*d*$_4$) δ 4.38 (d, *J* = 7.6 Hz, 1H, $H_{Gal}$-1), 4.31 (d, *J* = 8.4 Hz, 1H, $H_{Glc}$-4), 3.92 (dd, *J* = 12.2, 2.5 Hz, 1H, $H_{Glc}$-6a), 3.86 (dd, *J* = 12.1, 4.3 Hz, 1H, $H_{Glc}$-6b), 3.81 (d, *J* = 3.3 Hz, 1H, $H_{Gal}$-4), 3.76 (dd, *J* = 11.5, 7.6 Hz, 1H, $H_{Gal}$-6a), 3.72 (dd, *J* = 10.1, 8.4 Hz, 1H, $H_{Glc}$-2), 3.68 (dd, *J* = 11.6, 4.5 Hz, 1H, $H_{Gal}$-6b), 3.62 – 3.60 (m, 2H, $H_{Glc}$-3, $H_{Glc}$-4), 3.58 (dd, *J* = 7.8, 4.4 Hz, 1H, $H_{Gal}$-5), 3.53 (dd, *J* = 9.7, 7.6 Hz, 1H, $H_{Gal}$-2), 3.48 (dd, *J* = 9.7, 3.3 Hz, 1H, $H_{Gal}$-3), 3.46 (s, 3H, OMe), 3.44 – 3.38 (m, 1H, $H_{Glc}$-5), 1.97 (s, 3H, NHAc). $^{13}$C NMR (151 MHz, Methanol-*d*$_4$) δ 173.61, 105.09, 103.63, 80.83, 77.17, 76.60, 74.83, 74.34, 72.61, 70.34, 62.53, 61.92, 57.02, 56.51, 22.91. ESI MS(m/z): [M + Na]+ calcd for $C_{15}H_{27}NO_{11}Na$, 420.1482; found 419.6008.

**Preparation of substrate 7**

SGP (50 mg, 0.02 mmol) was dissolved in a sodium acetate buffer (PH 5.5, 50mM), containing calcium chloride (10 mM). Neuraminidase was added to the mixture, and the reaction was incubated at 37 °C with shaking until ESI indicated complete removal of sialic acid. The pH of the reaction mixture was then adjusted to 4.5 using acetic acid. Subsequently, BSA and β-galactosidase were introduced into the mixture, and the reaction was incubated at 37 °C with shaking until ESI indicated that all galactose residues had been removed. The protein in the reaction mixture was separated by centrifugation using a 10K MWCO filter. The filtrate was lyophilized and subjected to multiple rounds of purification via size exclusion chromatography (Bio-Rad P2 gel and P4 gel), eluting with a 0.1 M ammonium bicarbonate solution. Fractions containing the pure products were collected and lyophilized, resulting in the isolation of compound **7** (5.3 mg, 15.5%) as white powder. ESI MS(m/z): $[M - 2H]^{2-}$ calcd for $C_{78}H_{133}N_{13}O_{44}^{2-}$, 977.9290; found 977.9101.

**Kinetic studies**

The recombinant human fucosyltransferases were expressed following the reported procedures.[2] Ki values were determined using the GDP-Glo™ Glycosyltransferase Assay Kit (Promega). Kinetic parameters were determined by maintaining saturating concentration of acceptors while varying concentrations of GDP-Fuc (3-100 μM), and fluorinated GDP-Fuc (3-300 μM). Transfer reactions involving inhibitors were performed in a 25 μL reaction volume in PBS buffer, containing appropriate amounts of FUTs, acceptor, inhibitor and GDP-Fuc. Enzyme assay mixtures were incubated for 1 h at 37 °C and the reaction was stopped by adding 25 μL of GDP detection reagent. The assay plate was mixed with a plate shaker for 30 sec. and incubated at RT for 60 min. Luminescence was measured using a plate-reader. A control group, which included all reaction components except FUTs, was used as a blank. All experiments were performed in triplicate.

**Cell culture**

HL60, HEK293, and U-2 OS cells were obtained from ATTC. HL-60 cells were cultured in RPMI 1640 medium supplemented with fetal bovine serum (FBS; 10% v/v) and penicillin-streptomycin (1%). HEK293 cells were cultured in DMEM medium supplemented with FBS (10% v/v) and penicillin-streptomycin (1%). U-2 OS cells were cultured in McCoy's 5A medium supplemented with FBS (10% v/v) and penicillin-streptomycin (1%). All cell lines were maintained at 37 °C, 5% $CO_2$ in a humidified atmosphere.

**Reagents (antibodies and lectins)**

The sources of the antibodies and lectins used for flow cytometry experiments are as follows: biotinylated AAL (Vector Labs, B1395), biotinylated LCA (Vector Labs, B-1045), purified mouse anti-human CD15s (BD Pharmingen; 551344), streptavidin Alexa Fluor-488

(Invitrogen, S11223), R-PE anti–mouse IgM (Jackson ImmunoResearch, 115-116-075), FITC anti-human CD15 (SSEA-1) antibody (Biolegend, 301904).

**Flow cytometry**

For cell surface staining experiment, cells were seeded in a 96-well plate with 100 μL medium, after allowing the cells to settle for several hours, 100 μL of medium containing the inhibitor at the desired concentration was added. An equivalent concentration of DMSO was added to the control group, as all inhibitor stock solutions were prepared in DMSO. Cells were grown for 72 h at 37 °C in a humidified incubator. After this incubation period, cells were harvested and washed with FACS buffer (BSA/PBS, 1%, w/v). All staining procedures was performed in this buffer on ice for 30 min. Following staining, cells were washed 3 times with FACS buffer and then resuspended for analysis by flow cytometry (BD FACS Canto II, BD Biosciences). The staining solution were prepared with the following concentrations: For AAL, 1 μg/mL AAL-biotin was used, followed by 1 μg/mL streptavidin AF-488. For LCA, 1 μg/mL LCA-biotin was used, followed by 1μg/mL streptavidin AF-488. For SLe$^x$ detection, 0.1 μg/mL primary antibody and 10 μg/mL R-PE anti-mouse IgM were used. Experiments were performed in triplicate and repeated at least 3 times. The data were normalized to the control cells treated with DMSO only (100%) and unstained cells (0%).

**Cell proliferation and viability**

Cell viability was assessed using trypan blue staining, while cell proliferation was determined with CellTiter 96 Aqueous One Solution Cell Proliferation Assay kit (MTS assay). Briefly, cells were incubated with desired concentrations of inhibitors for 3 days, after which the assay solutions were added. The assays were performed following the enclosed manual. Data were recorded using a Promega plate reader.

**Sugar nucleotide analysis**

HL-60 cells were treated for 3 days with either DMSO only, **1**, **3**, or **4** at concentrations of 8, 32, and 64 μM. After 72 h, the cells were pelleted and washed twice with cold washing buffer (75 mM ammonium carbonate in Milli-Q $H_2O$, adjusted to pH 7.4 with acetic acid). To each sample, 700 μL of cold extraction buffer (acetonitrile:methanol:water as 40:40:20) was added for 2 min. The supernatant was then transferred to a separate vial. This extraction process was repeated with another 700 μL cold extraction solvent for 3 min. The two extracts were pooled and centrifuge at 13,000 rpm for 3 min. in a pre-cooled centrifuge. The resulting supernatant was transferred to a 1.5 mL eppendorf vial and the samples was dried in by *vacuo* centrifugation at RT. The dried samples were stored at -80 °C until they were analyzed by mass spectrometry. GDP-Fuc, GDP-2-F-Fuc, and GDP-2,2-diF-Fuc were used as internal standards.

**Glycan profiling**[3]

HL-60 cells were cultured in medium containing either DMSO, 8 or 32 μM of each inhibitor. After 3 days, the cells were pelleted and washed three times with PBS. Cell pellets were lysed with RIPA lysis buffer on ice for 0.5 h, followed by centrifugation at 16,500 rcf to collect the cell lysate supernatant. To prepare the lysates for enzymatic *N*-glycan release, proteins were denatured and reduced in DTT/SDS (40 mM DTT, 0.5% v/v SDS) for 8 min at 95°C. NP-40 (1% v/v) and sodium phosphate buffer (50 mM, pH 7.5) were added after cooling to RT. Finally, PNGase F was added and incubated at 37 °C overnight. The released glycan solution was purified by C18 SPE (Avantor™ 7020-02 BAKERBOND™ SPE Octadecyl). The cartridges were conditioned with 1 mL MeCN, followed by 1 mL Milli-Q $H_2O$. The glycan solution was loaded onto the cartridge, collected, and then residual glycans were eluted with 1 mL 5% MeCN 0.05% TFA in Milli-Q $H_2O$. The C18 load and elution fractions were partially evaporated under nitrogen gas for 1 h, and then pooled to be applied to PGC SPE (Thermo Scientific™ HyperSep™ Hypercarb™ SPE cartridges). For PGC SPE cartridges, the conditioning steps were identical to C18 cartridges. After loading the pooled C18 fractions, three washing steps were performed: first, 1 mL of 0.05% TFA in Milli-Q $H_2O$, then 1 mL 5% MeCN and 0.05% TFA in Milli-Q $H_2O$, and finally elution with 1 mL 50% MeCN and 0.1% TFA in Milli-Q $H_2O$. The elution fraction was dried completely under a stream of nitrogen gas. The resulting samples (free reducing end *N*-glycans) were labelled with procainamide as follows: samples were dissolved in 30 μL Milli-Q $H_2O$ and 6 μL glacial acetic acid, to which 10 μL procainamide and 2-picoline borane complex (2-PB) mix was added (53 mg/mL procainamide HCl, 54 mg/mL 2-PB in DMSO). The labelling reaction was kept at 65°C for 2 h and then a liquid-liquid extraction was performed to remove residual procainamide. 300 μL Milli-Q $H_2O$ was added to the labeling mixture and adjusted to pH >10 with ammonia solution (25% w/v in $H_2O$). Washes with 500 μL DCM were performed three times. The DCM was discarded and the remaining aqueous phase was evaporated to dryness under a stream of nitrogen gas. Finally, the samples were desalted by reconstitution in 1 mL Milli-Q $H_2O$ and PGC SPE as described above.

The resulting dried samples were made ready for LC-MS analysis on a Agilent 1260 Infinity LC coupled to a 6560 IM-QTOF mass spectrometer (Agilent Technologies, Santa Clara, USA). The samples were reconstituted in 20 mL 7:3 MeCN:Milli-Q $H_2O$ with an injection volume of 15 µL. For LC-separation, a SeQuant ZIC-cHILIC column (3 µm, 100 Å; 150 × 2.1 mm) was used with a matching guard column (20 × 2.1 mm). The compartment temperature was 40 °C. The mobile phase was composed of eluent A: 10 mM ammonium formate with 10 mM formic acid in Milli-Q $H_2O$, and eluent B: MeCN. The initial eluent composition was 30% A at a flow rate of 0.2 mL/min, followed by a linear gradient to 50% A from 0 to 20 min. 50% A was held isocratically until 25 min.

The IMS-QTOF was set to positive ion mode with a capillary voltage of 3,500 V, nozzle voltage of 2,000 V, and a fragmentor voltage of 360 V. The drying gas temperature was 300 °C with a flow rate of 8 L/min and the sheath gas temperature was 300 °C at 11 L/min. The nebulizer pressure was set at 40 psi.

MS data were calibrated to reference signals of m/z 121.050873 and 922.009798 using the IM-MS reprocessor utility of the Agilent Masshunter software. The mass-calibrated data was then demultiplexed using the PNNL preprocessor software using a 5-point moving average smoothing and interpolation of 3 drift bins. To find potential glycan hits in the processed data, the "find features" (IMFE) option of the Agilent IM-MS browser with an m/z range of 300–3,200 and an abundance of over 500 (peak area of most abundant ion). The list of glycan features was manually matched to *N*-glycan compositions using the ExPASy GlycoMod tool with the following parameters: monoisotopic mass values, 5 ppm mass tolerance and monosaccharide residues: Hexose, HexNAc, Deoxyhexose, and NeuAc. The resulting *N*-glycan composition lists were sorted by relative abundance, which are visualized in Fig. 5.

### 4) NMR spectra

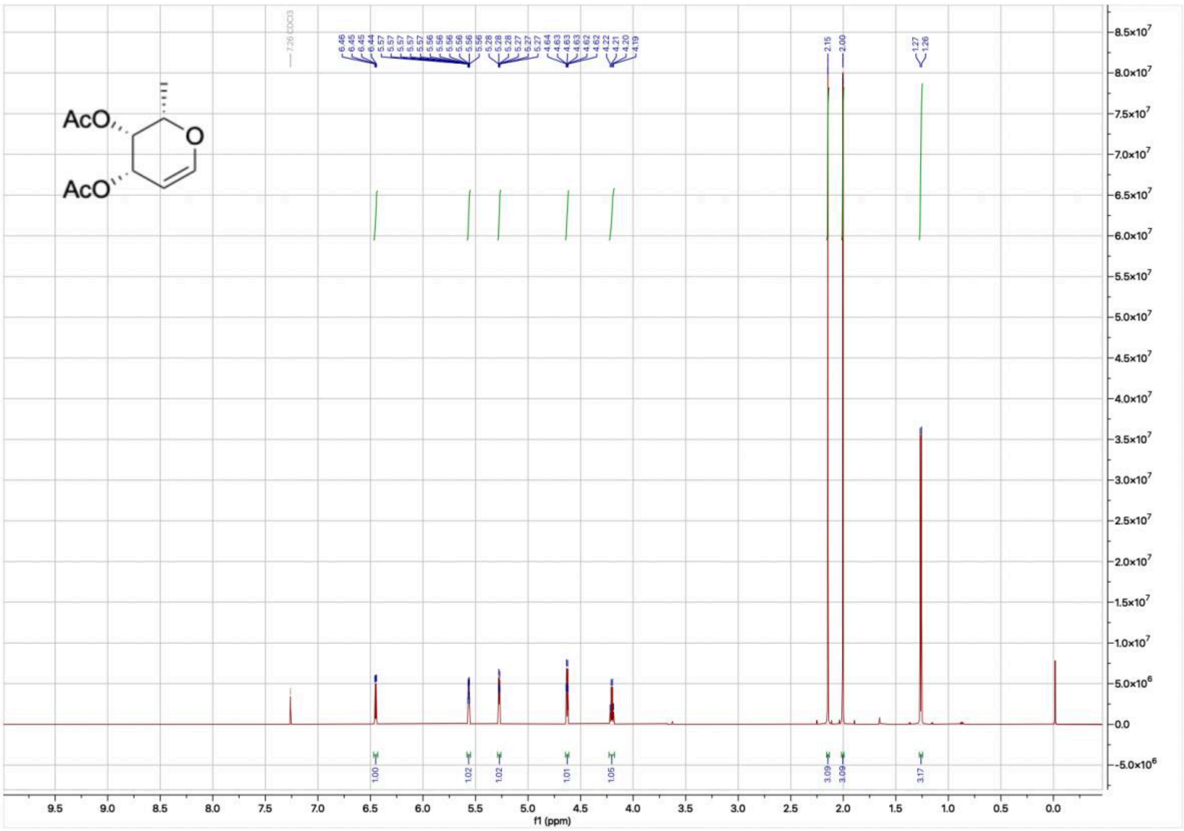


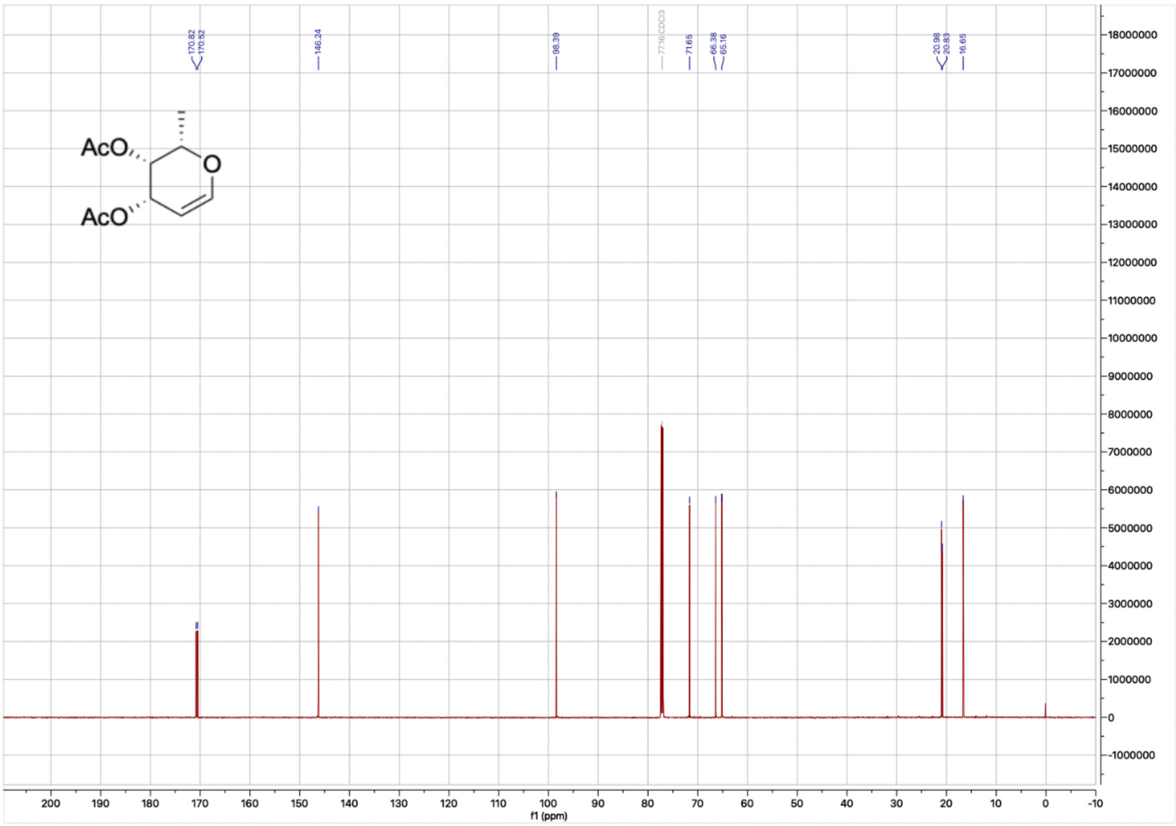

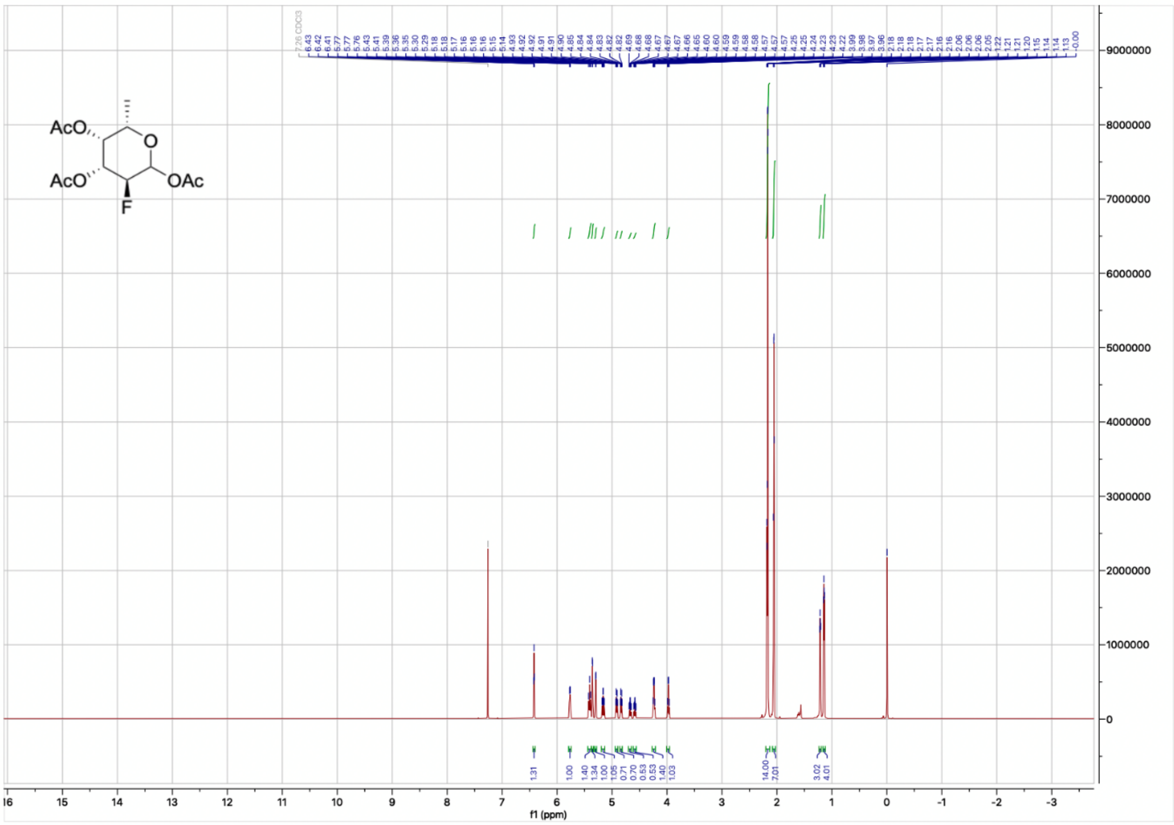
7.26 CDCl3
6.43 6.42 6.41 5.77 5.77 5.76 5.43 5.41 5.39 5.36 5.35 5.30 5.29 5.18 5.18 5.17 5.16 5.16 5.16 5.15 5.14 4.93 4.92 4.92 4.91 4.91 4.90 4.85 4.84 4.84 4.83 4.82 4.82 4.69 4.68 4.68 4.67 4.67 4.67 4.66 4.65 4.60 4.60 4.59 4.59 4.58 4.58 4.57 4.57 4.57 4.25 4.25 4.24 4.23 4.23 4.22 3.99 3.98 3.97 3.96 2.18 2.18 2.18 2.17 2.17 2.16 2.16 2.06 2.06 2.06 2.05 1.22 1.21 1.21 1.20 1.15 1.14 1.14 1.13 -0.00
AcO
AcO
O
OAc
F
1.31 1.00 1.40 1.34 1.00 1.05 0.71 0.70 0.53 0.53 1.40 1.03 14.00 7.01 3.02 4.01
f1 (ppm)

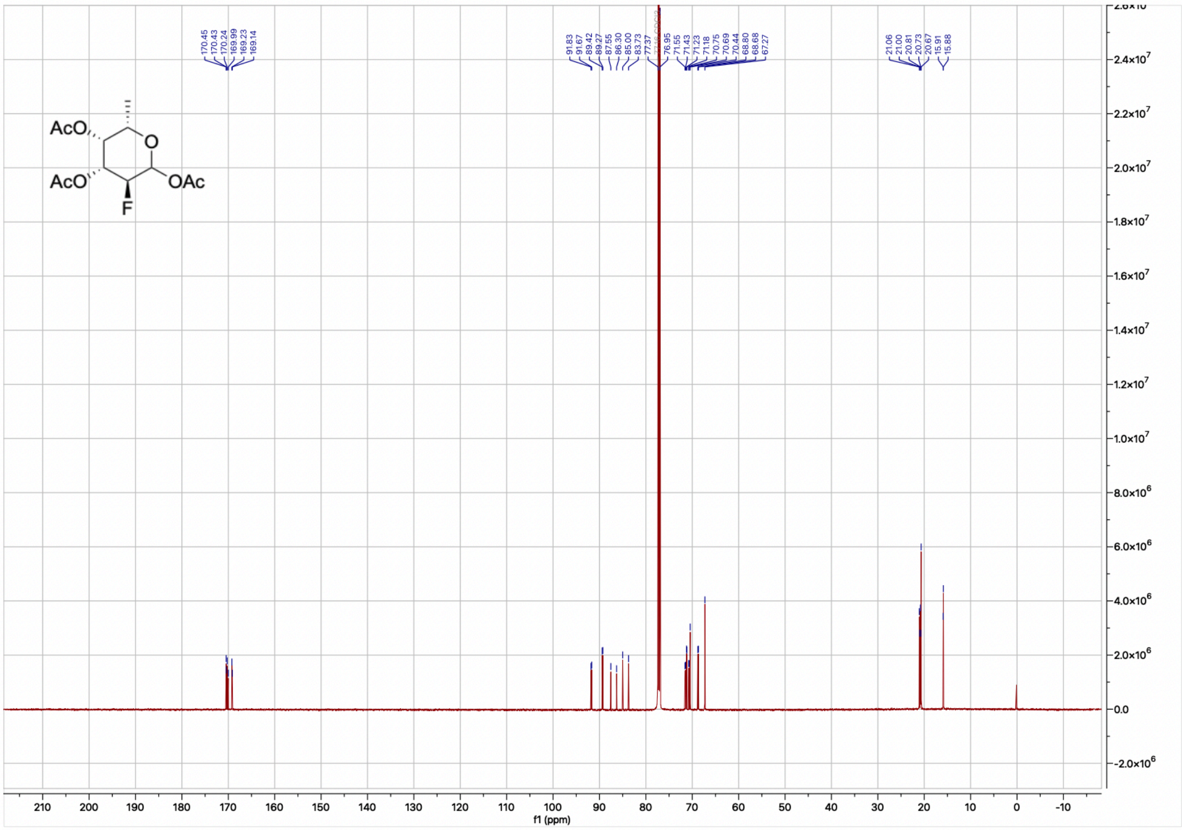
170.45 170.43 170.24 169.99 169.23 169.14
91.83 91.67 89.42 89.27 87.55 86.30 85.00 83.73 77.37 77.16 CDCl3 76.95 71.55 71.43 71.23 71.18 70.75 70.69 70.44 68.80 68.68 67.27
21.06 21.00 20.81 20.73 20.67 15.91 15.88
AcO
AcO
O
OAc
F
f1 (ppm)

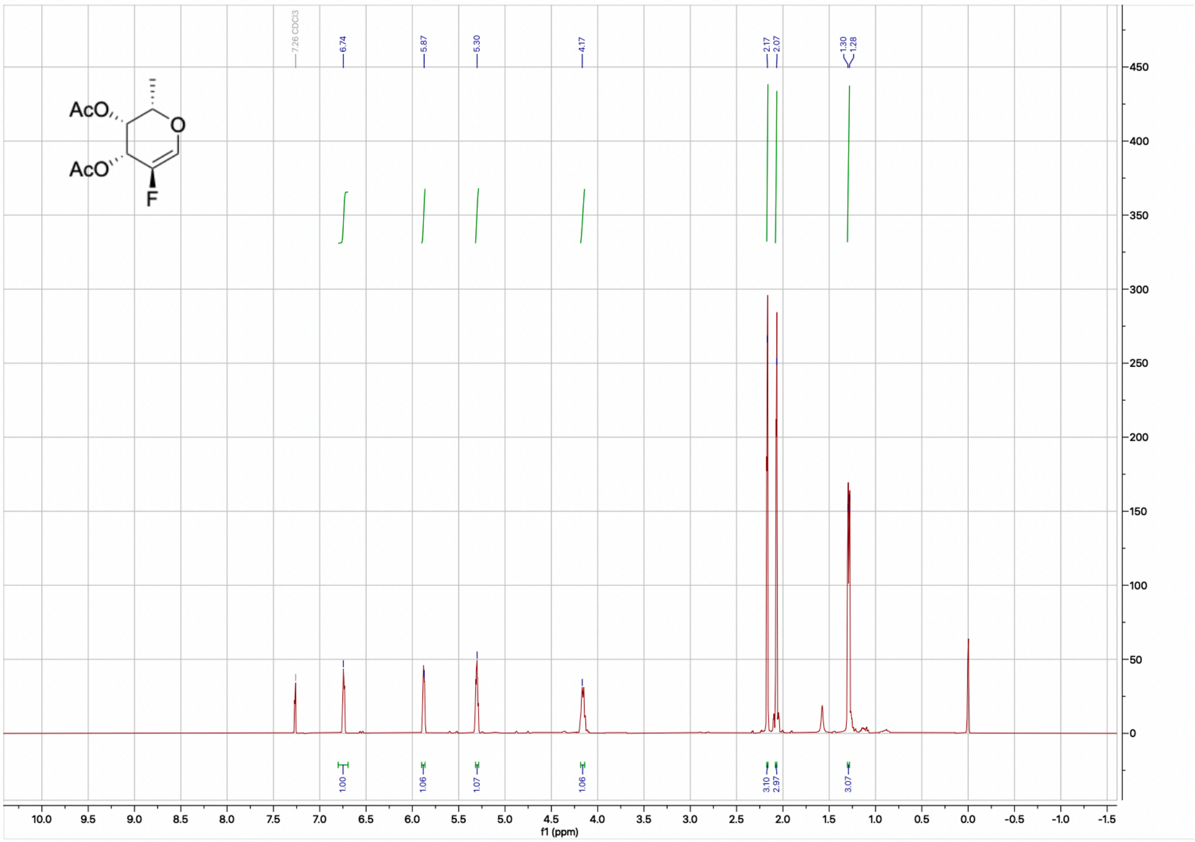
7.26 CDCl3
6.74
5.87
5.30
4.17
2.17
2.07
1.30
1.28
1.00
1.06
1.07
1.06
3.10
2.97
3.07
f1 (ppm)

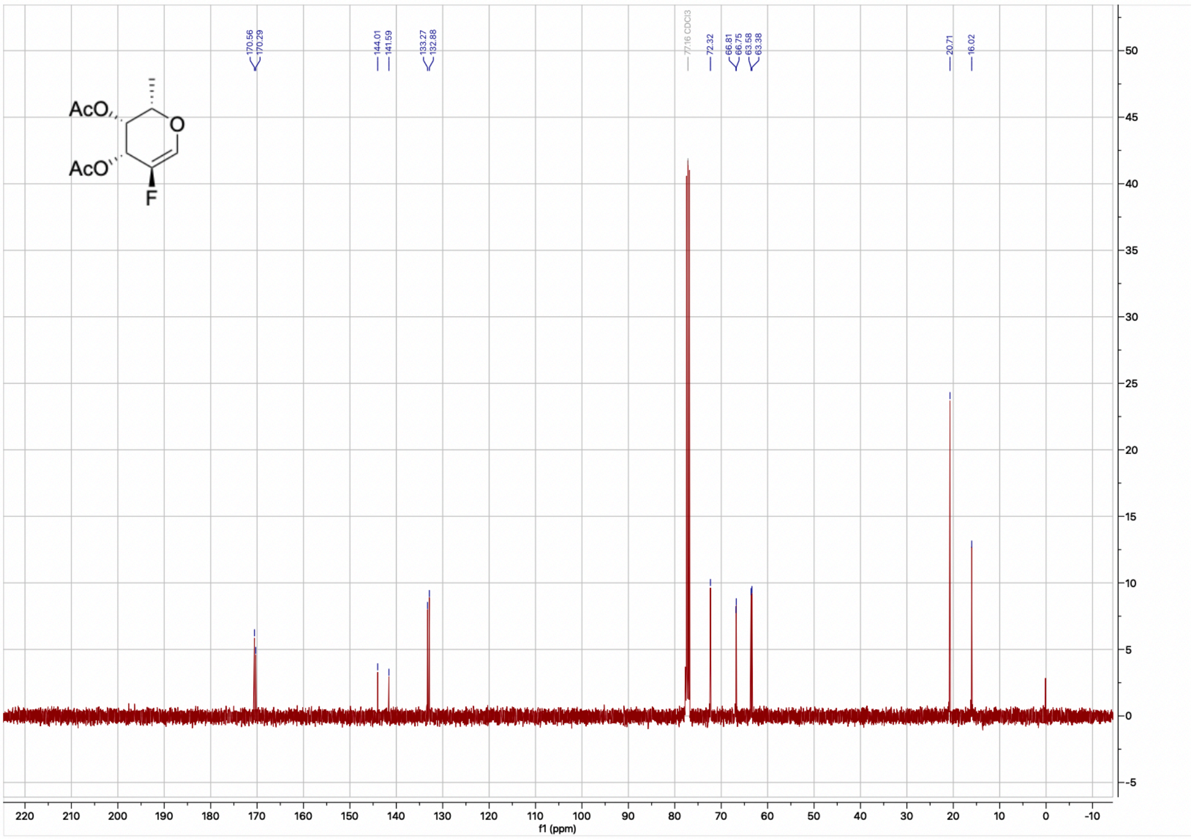
170.56
170.29
144.01
141.59
133.27
132.88
77.16 CDCl3
72.32
66.81
66.75
63.58
63.38
20.71
16.02
f1 (ppm)

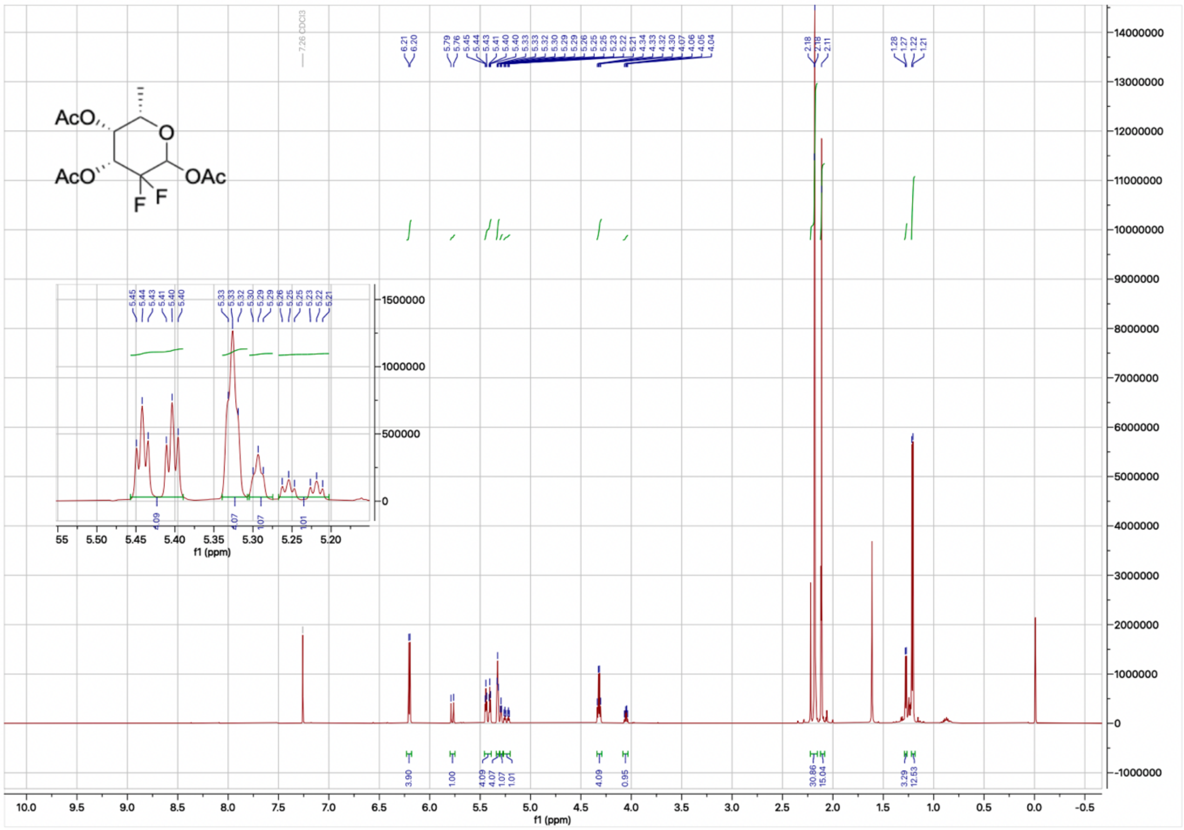
AcO
AcO
O
OAc
F F
f1 (ppm)

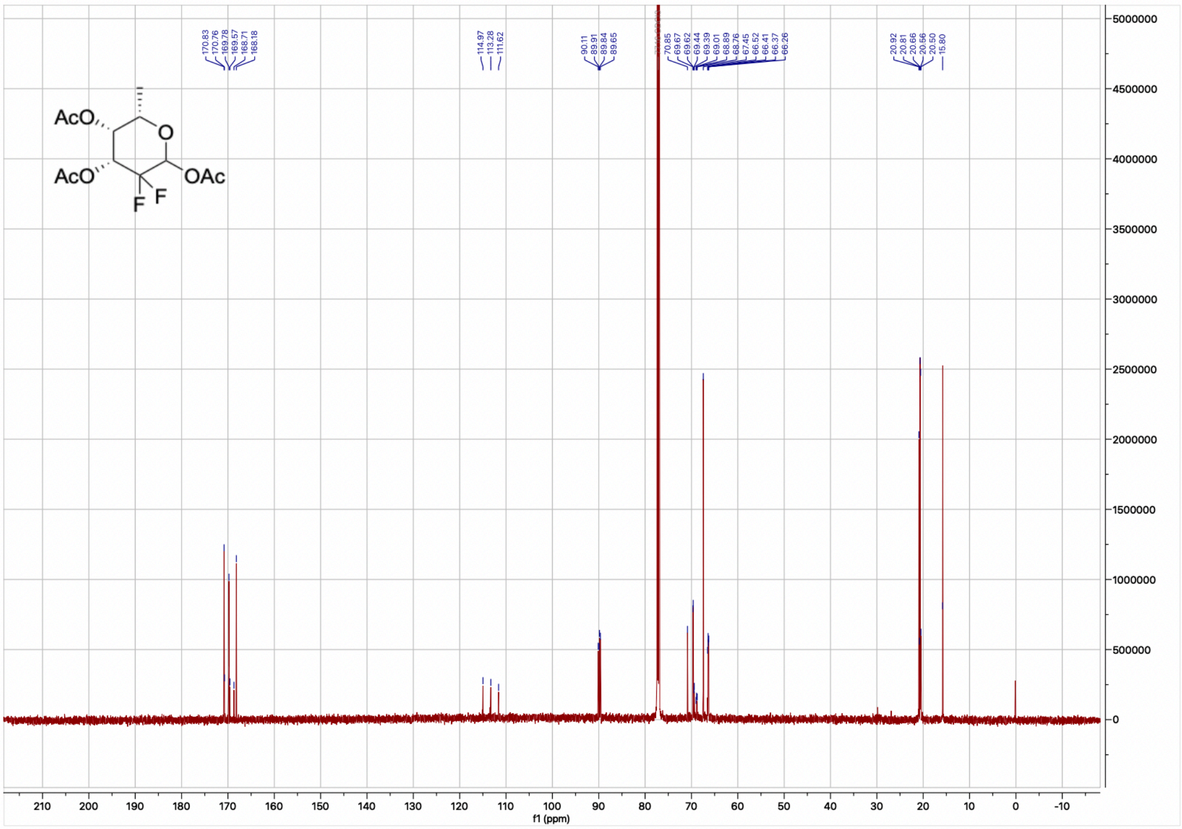
AcO
AcO
O
OAc
F F
f1 (ppm)

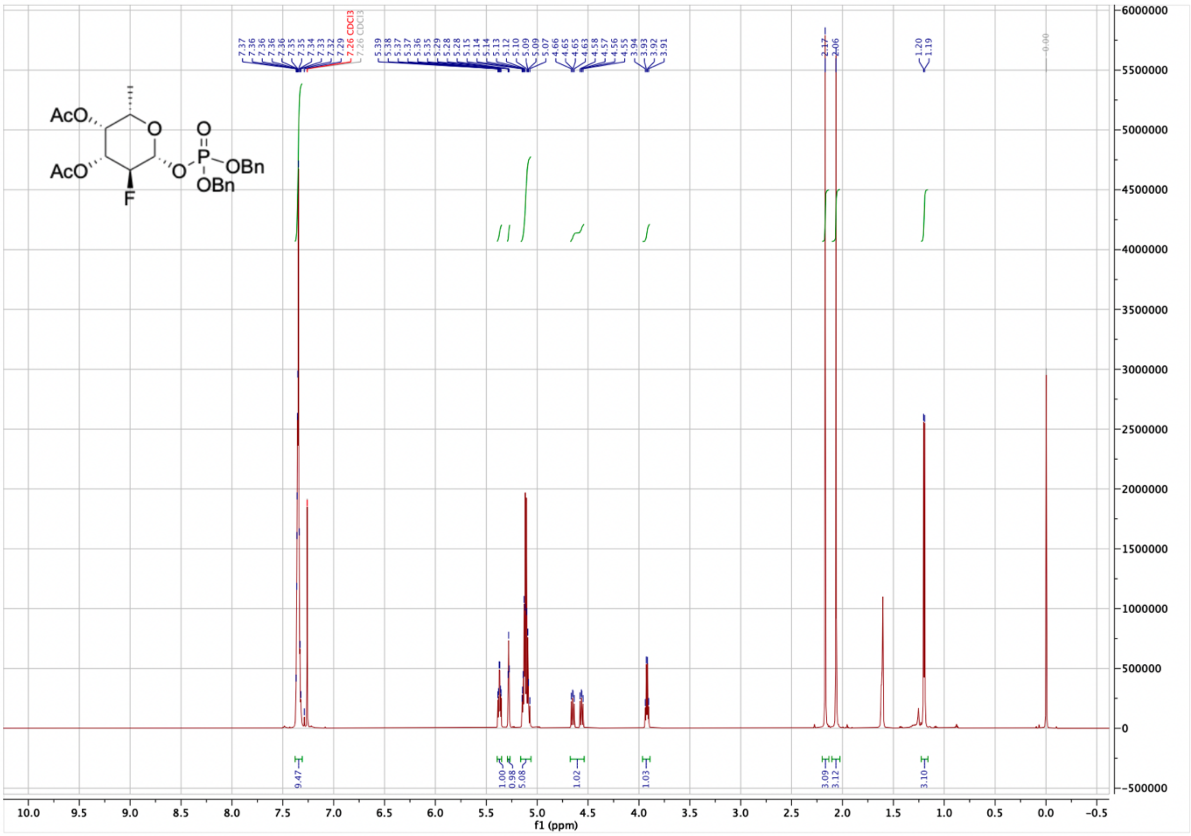

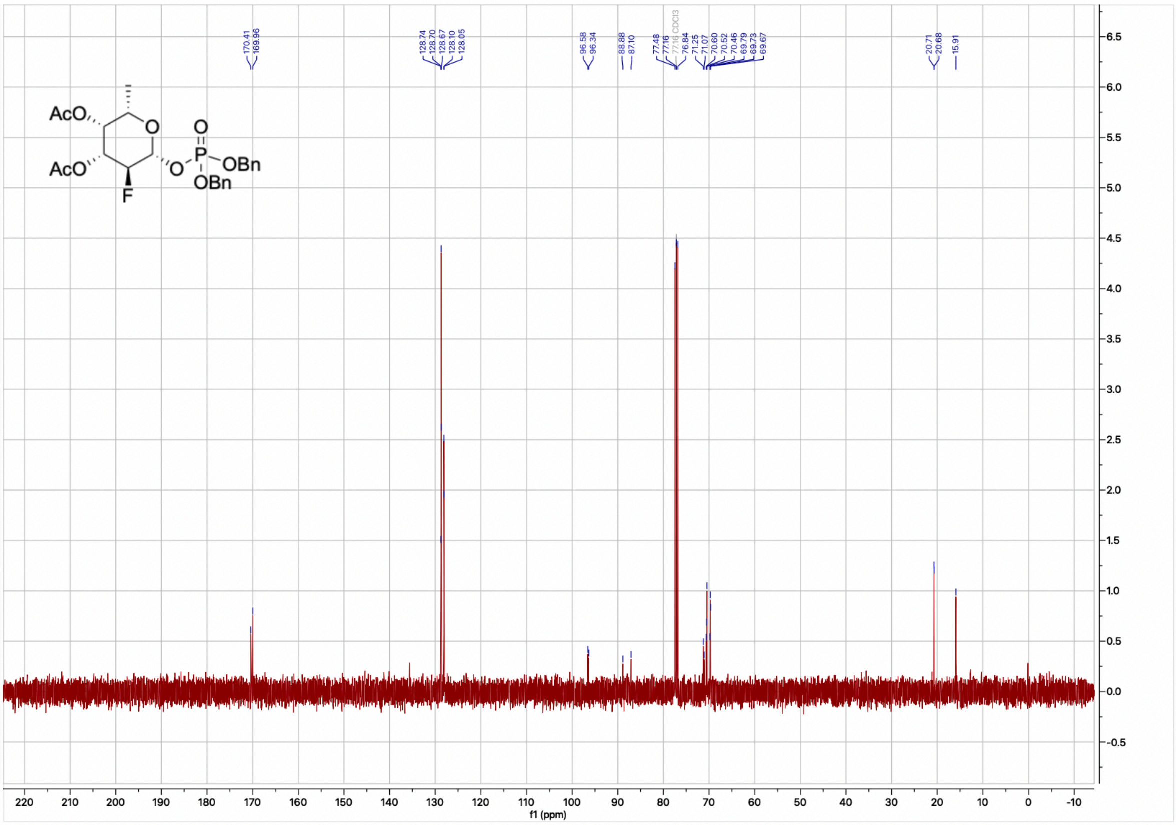

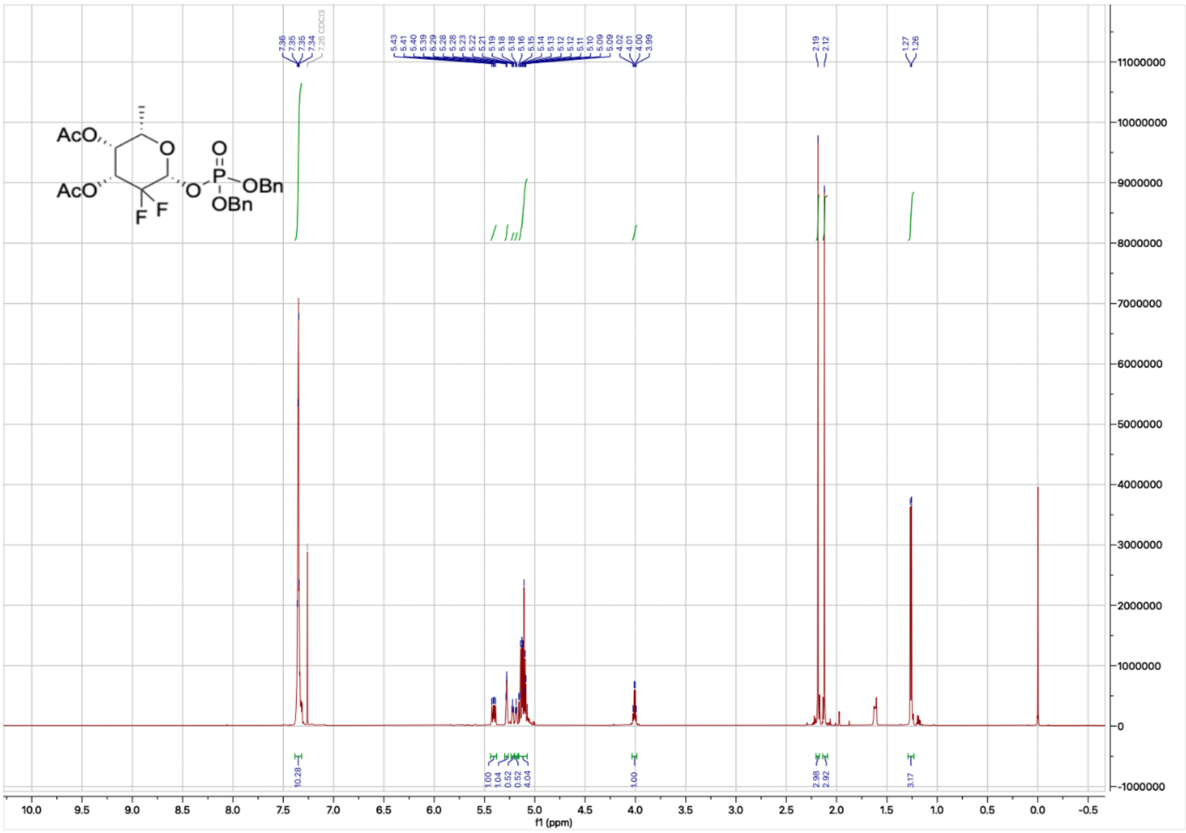
AcO
AcO
O
O
P
OBn
OBn
F F
7.36 7.35 7.35 7.34 7.26 CDCl3
4.02 4.01 4.00 3.99
2.19 2.12
1.27 1.26
10.28
1.00 1.04 0.52 0.52 4.04
1.00
2.98 2.92
3.17
f1 (ppm)

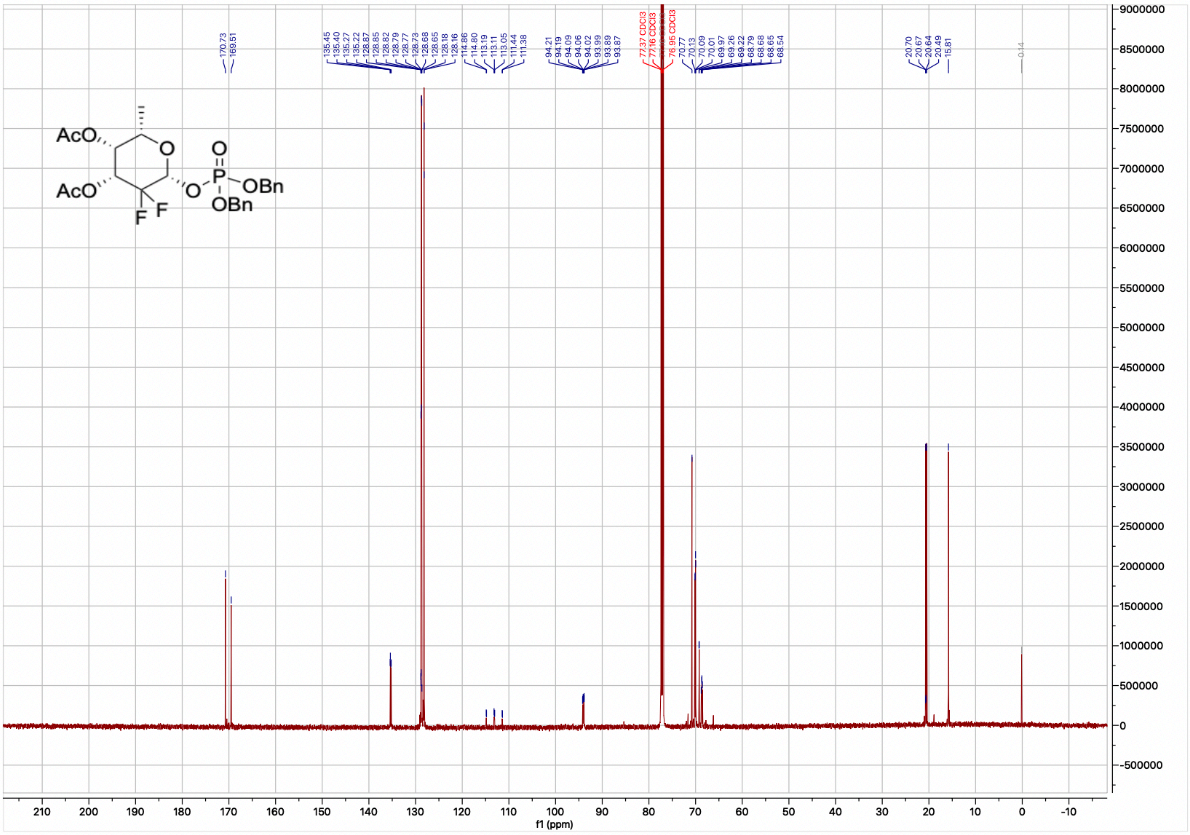
AcO
AcO
O
O
P
OBn
OBn
F F
170.73 169.51
77.37 CDCl3
77.16 CDCl3
76.95 CDCl3
20.70 20.67 20.64 20.49 15.81
f1 (ppm)

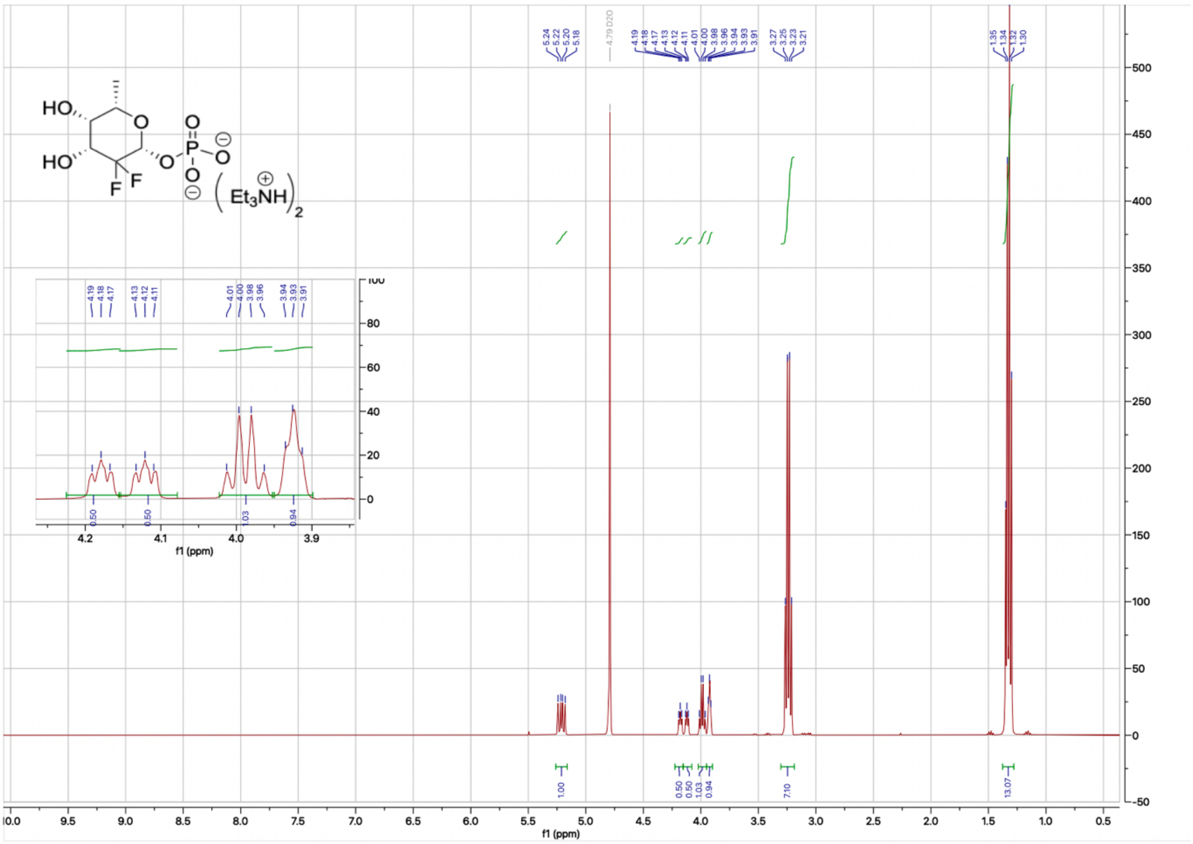
5.24
5.22
5.20
5.18
4.79 D2O
4.19
4.18
4.17
4.13
4.12
4.11
4.01
4.00
3.98
3.96
3.94
3.93
3.91
3.27
3.25
3.23
3.21
1.35
1.34
1.32
1.30
HO
HO
O
O
P
O
O
F
F
Et3NH
2
1.00
0.50
0.50
1.03
0.94
7.10
13.07
f1 (ppm)

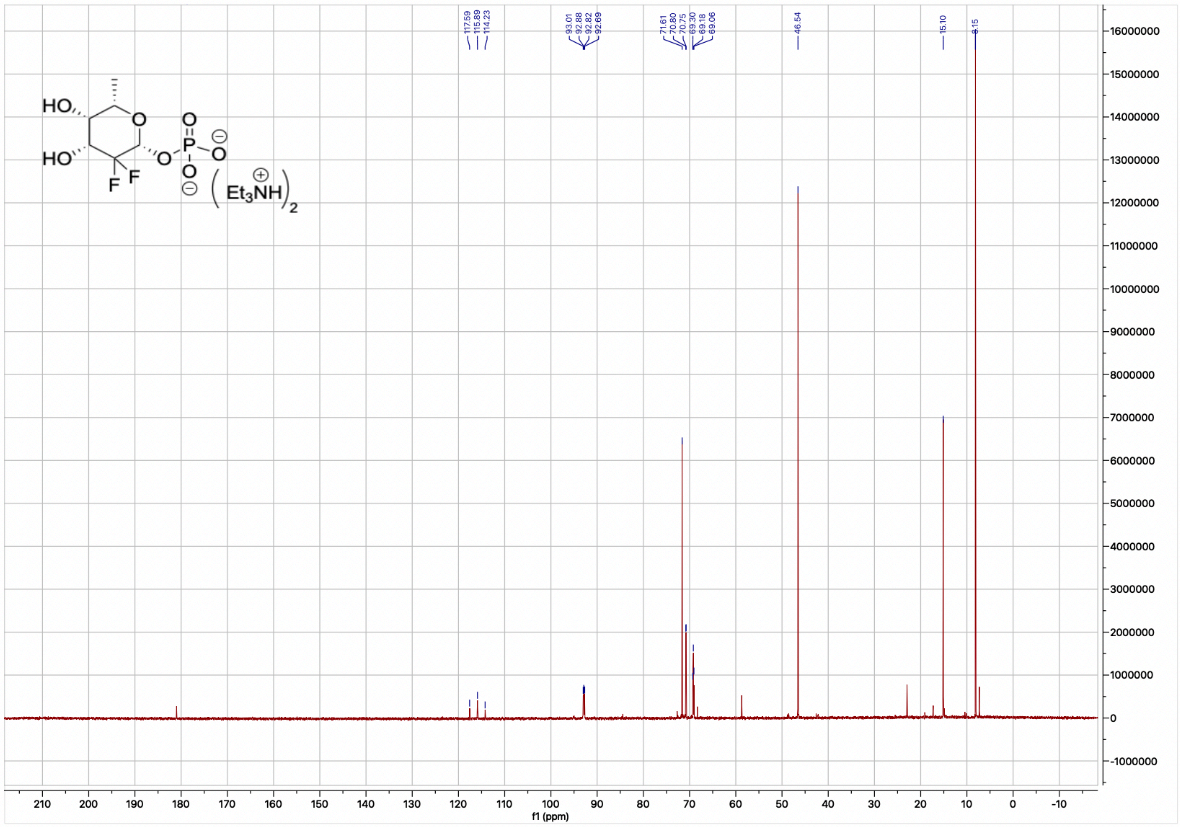
117.59
115.89
114.23
93.01
92.88
92.82
92.69
71.61
70.80
70.75
69.30
69.18
69.06
46.54
15.10
8.15
HO
HO
O
O
P
O
O
F
F
Et3NH
2
f1 (ppm)

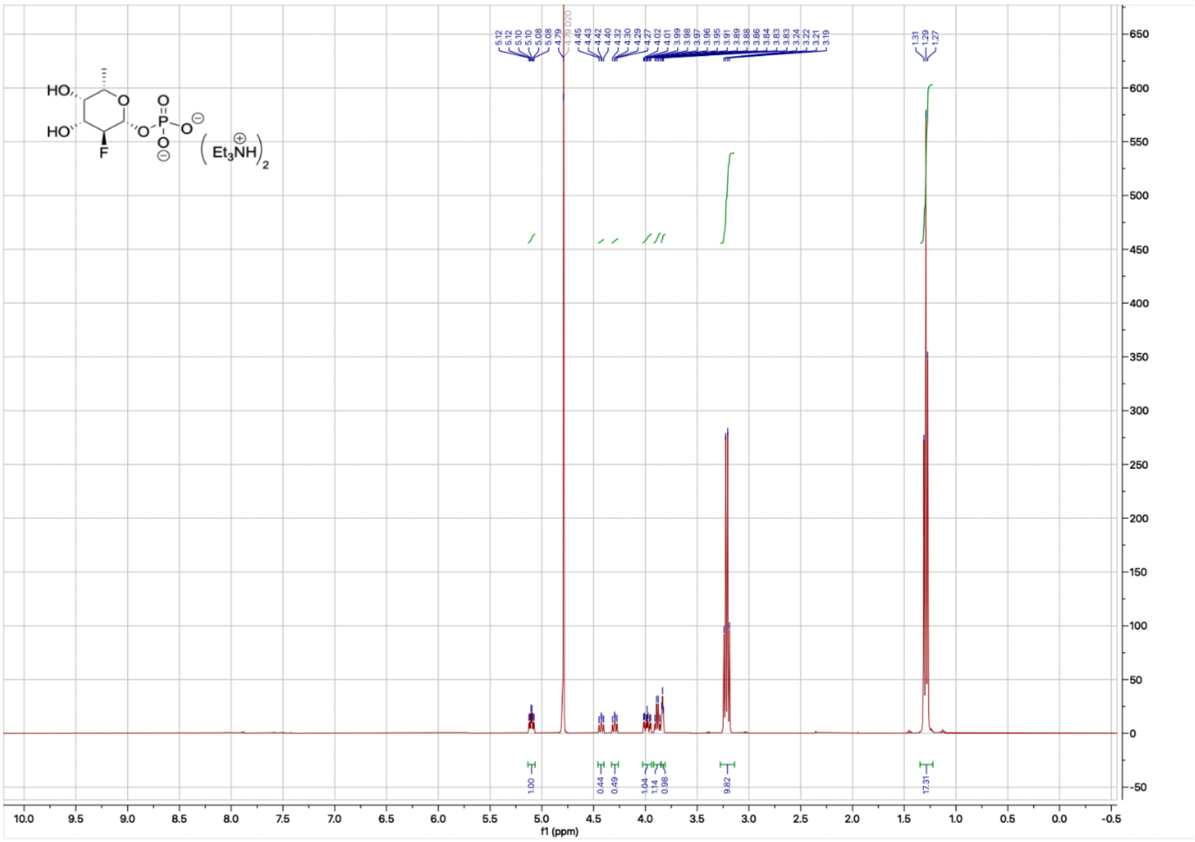

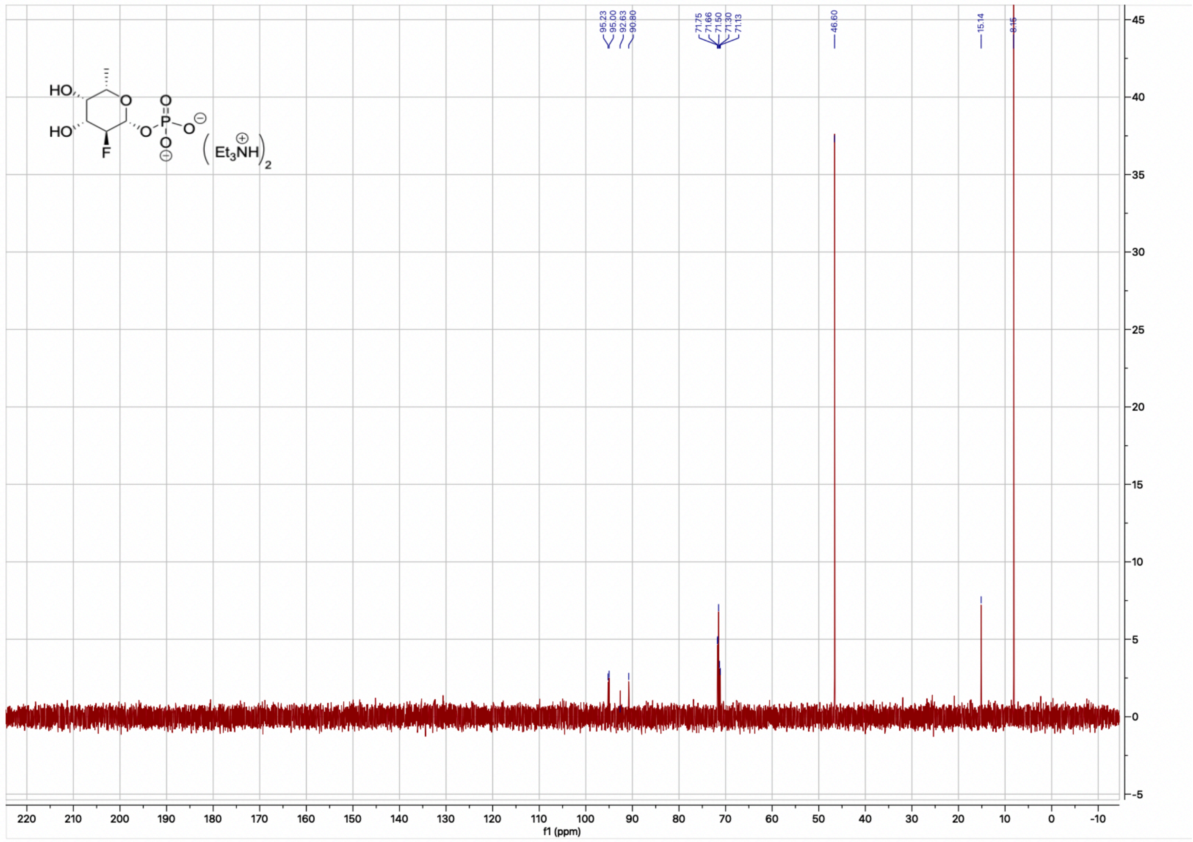

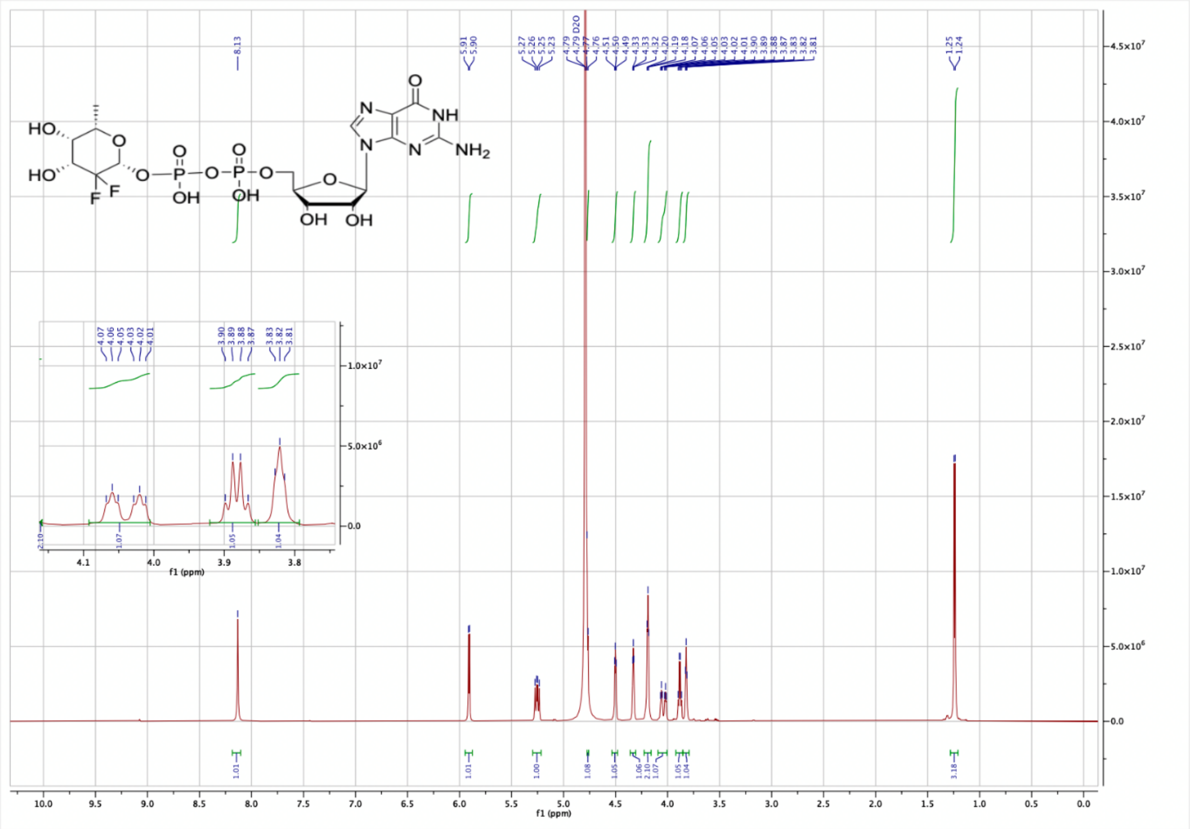

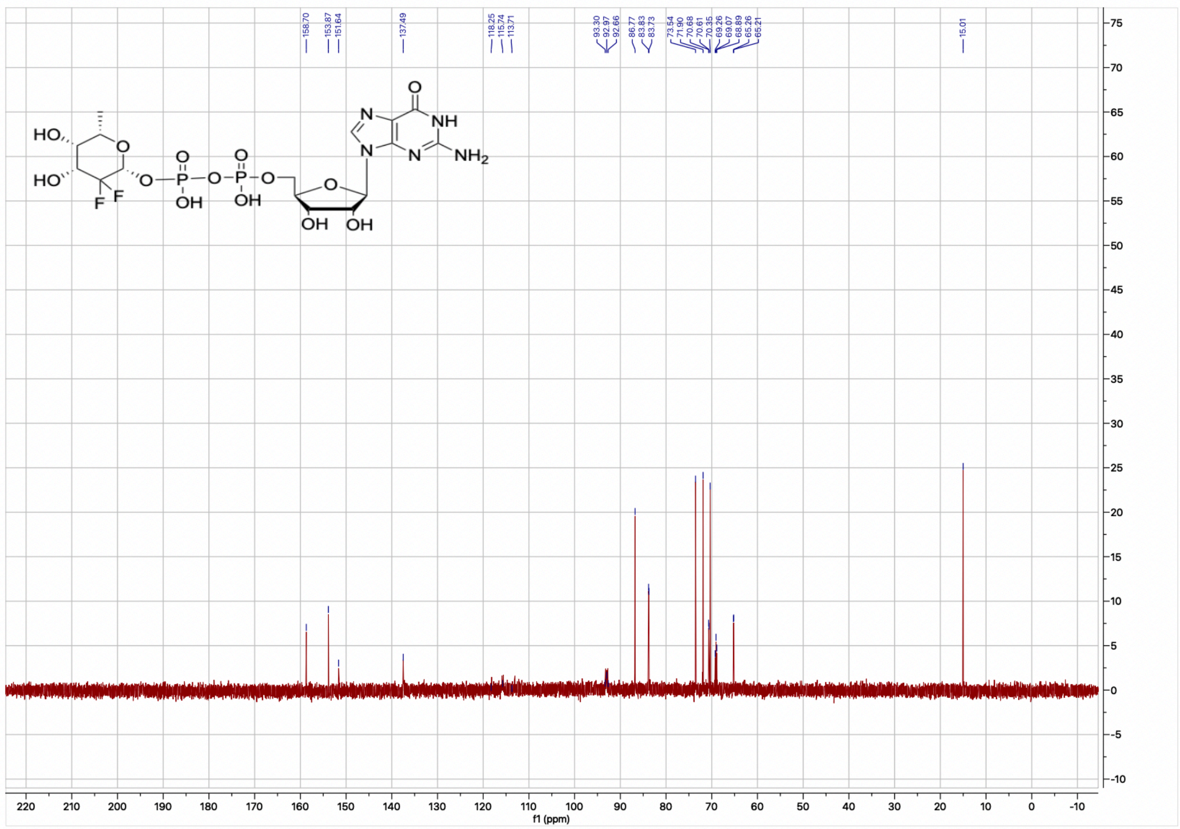

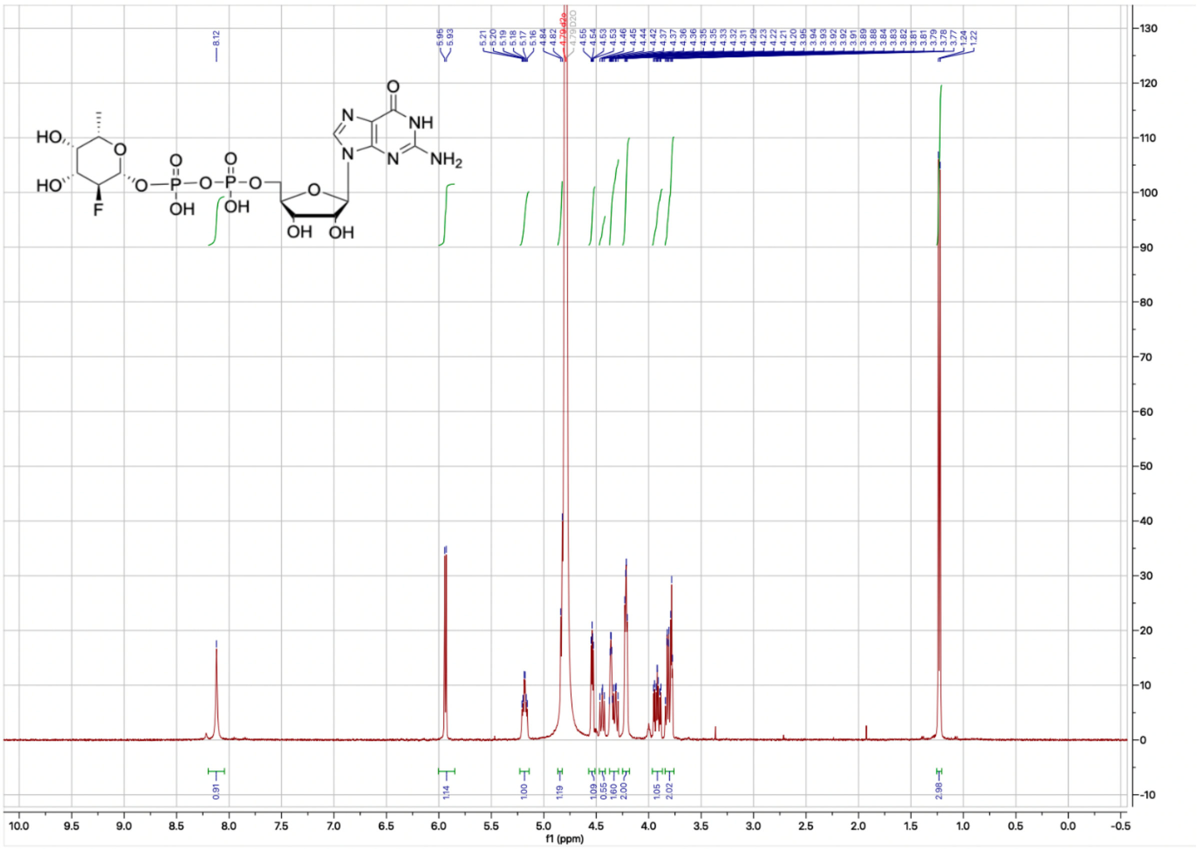

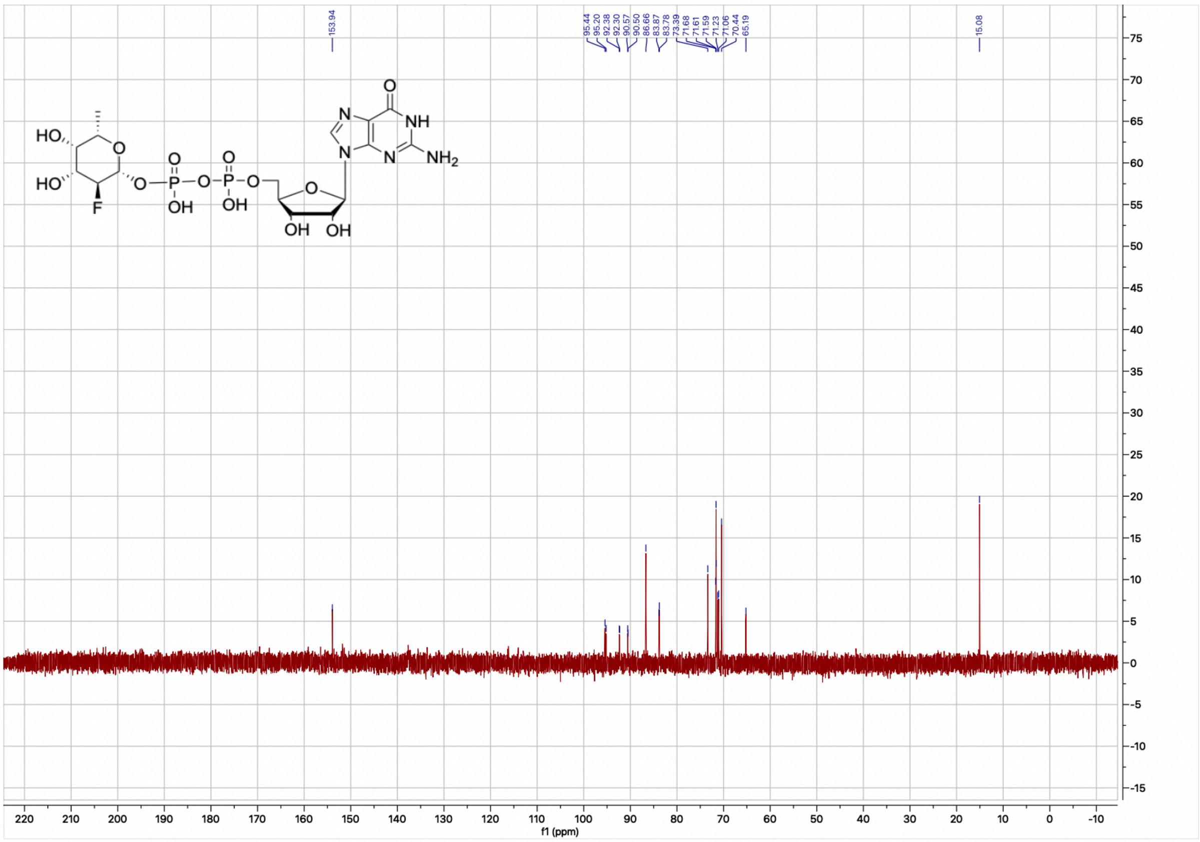

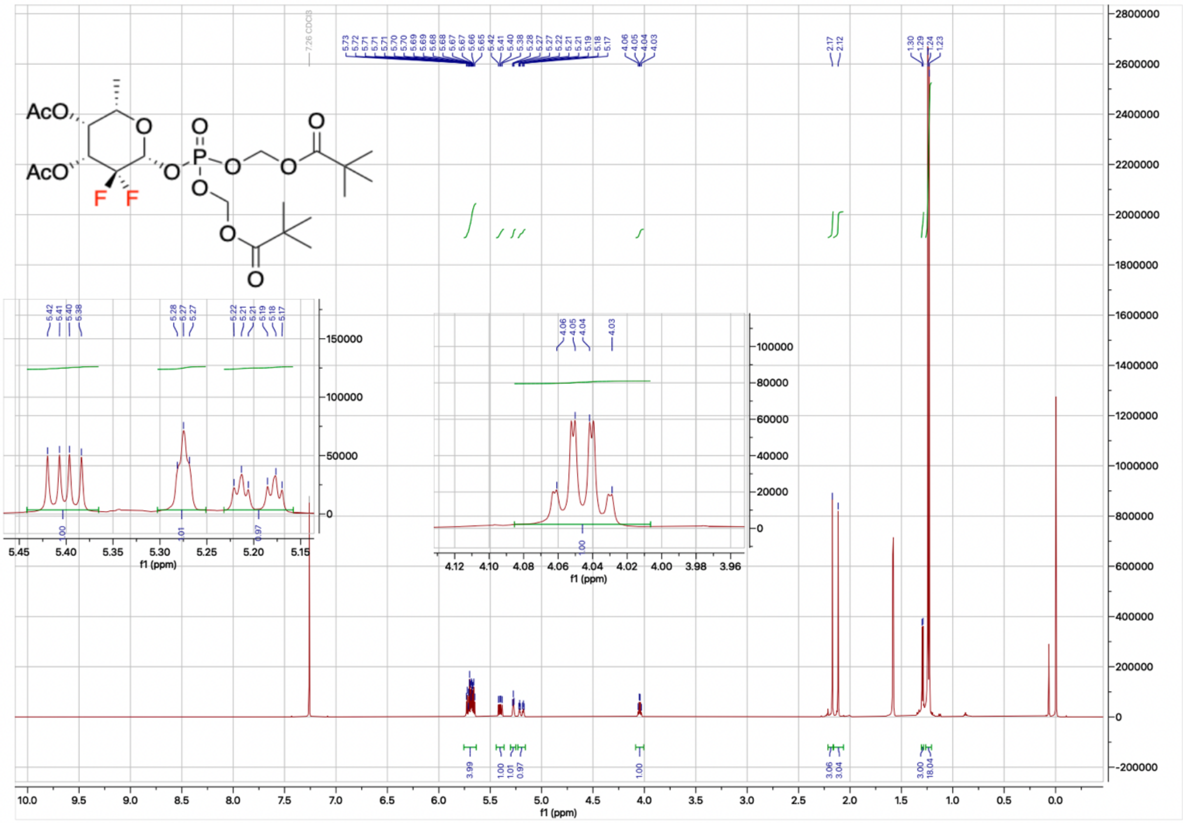

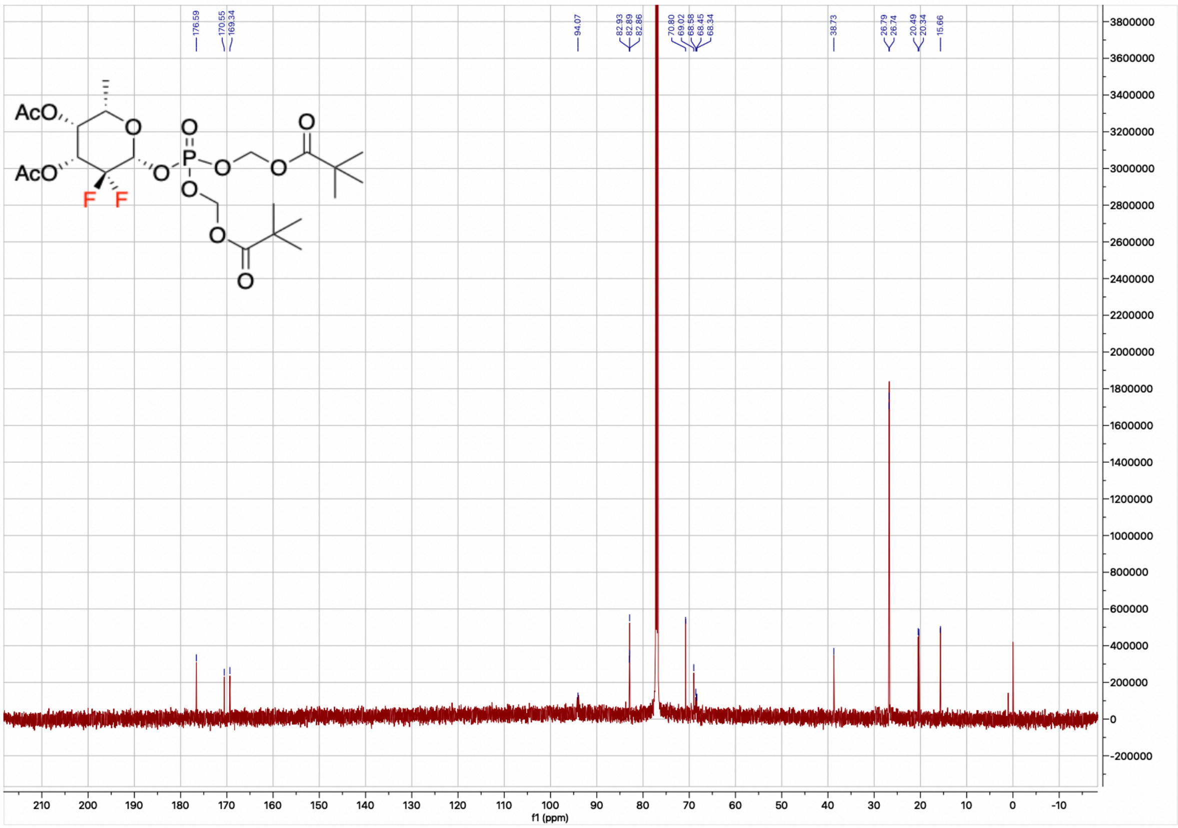

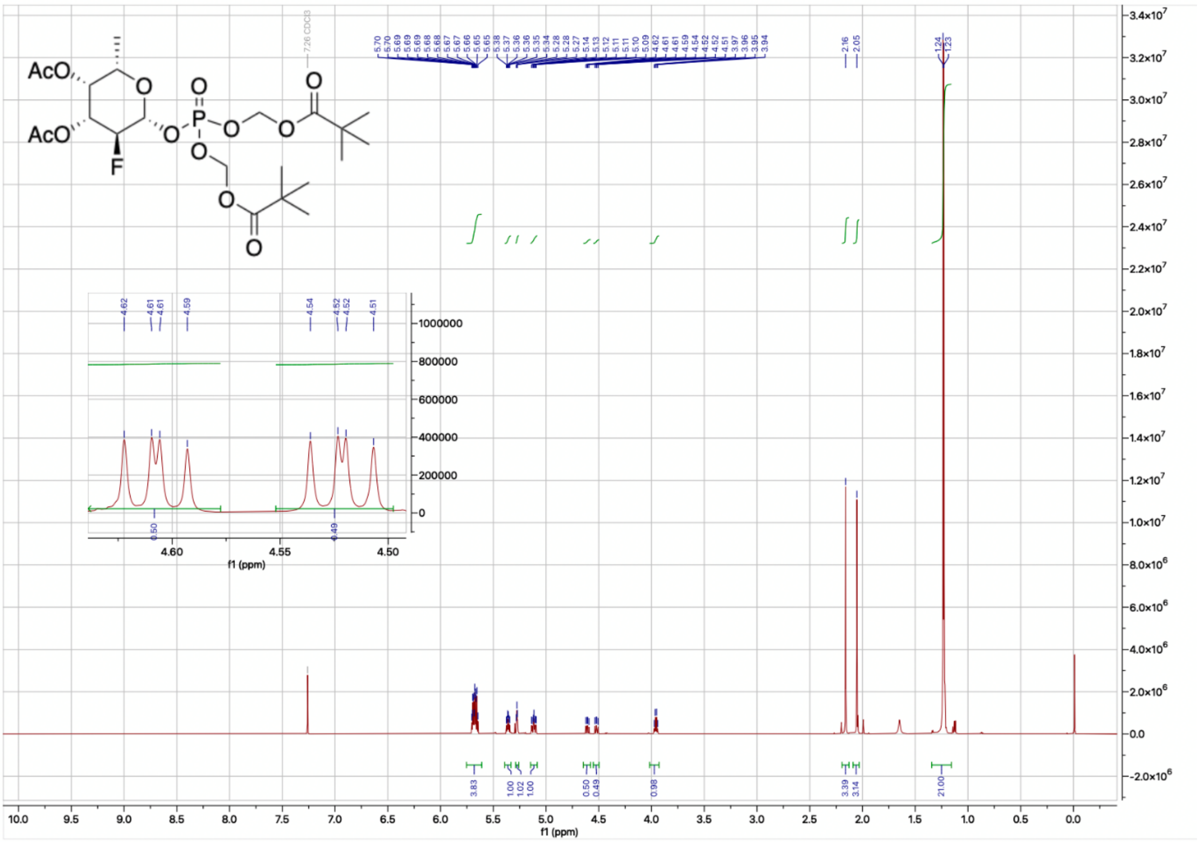

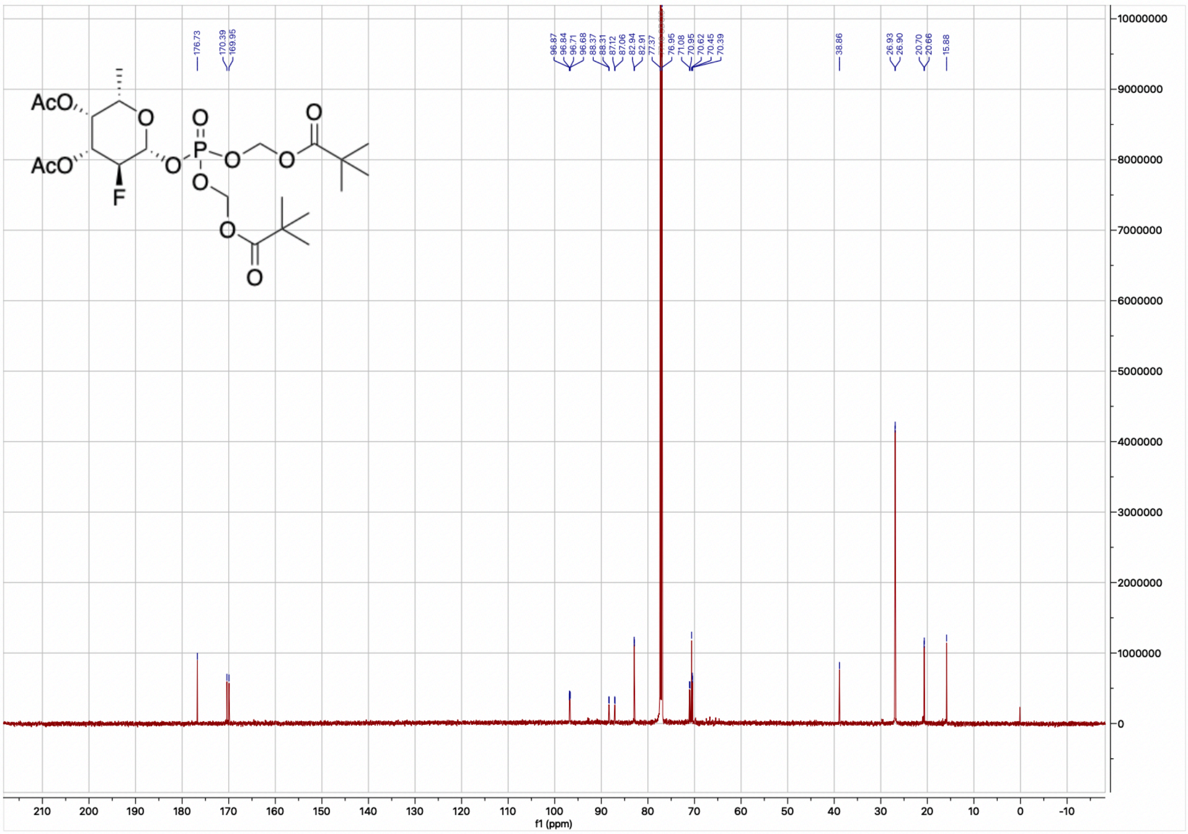

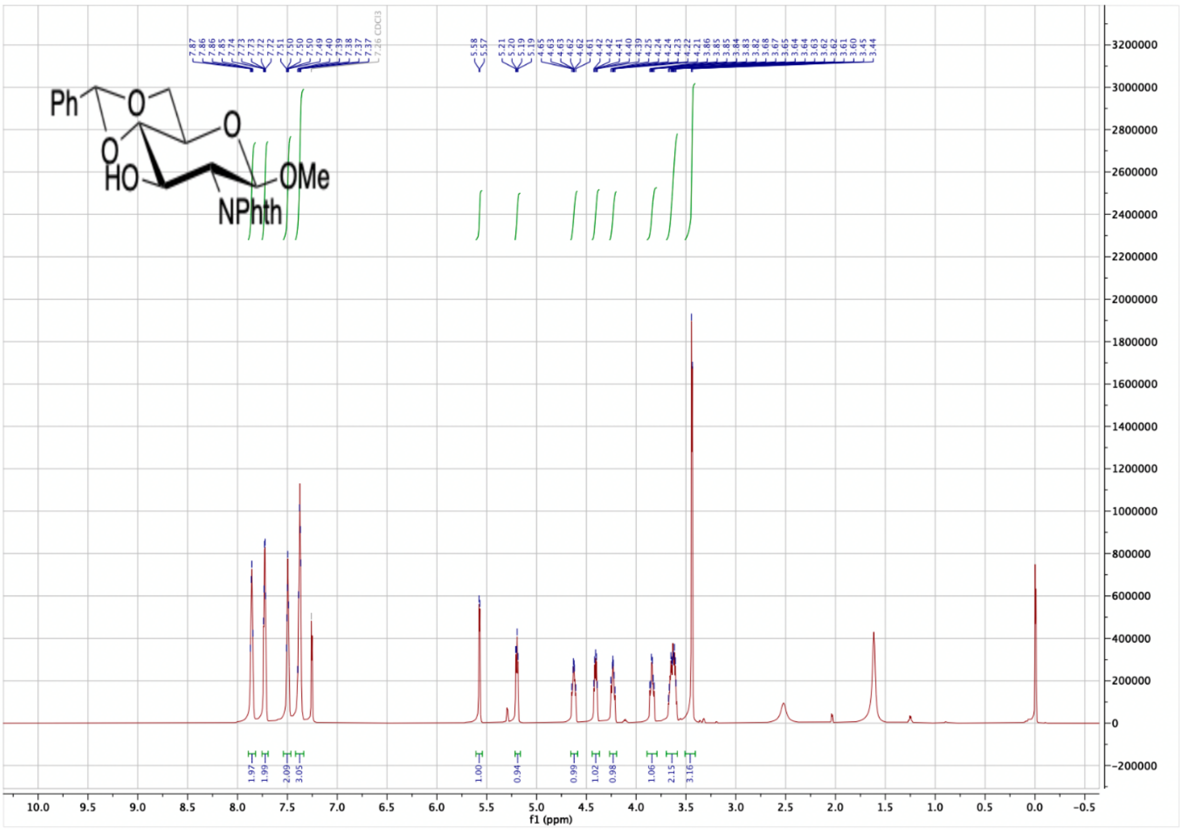

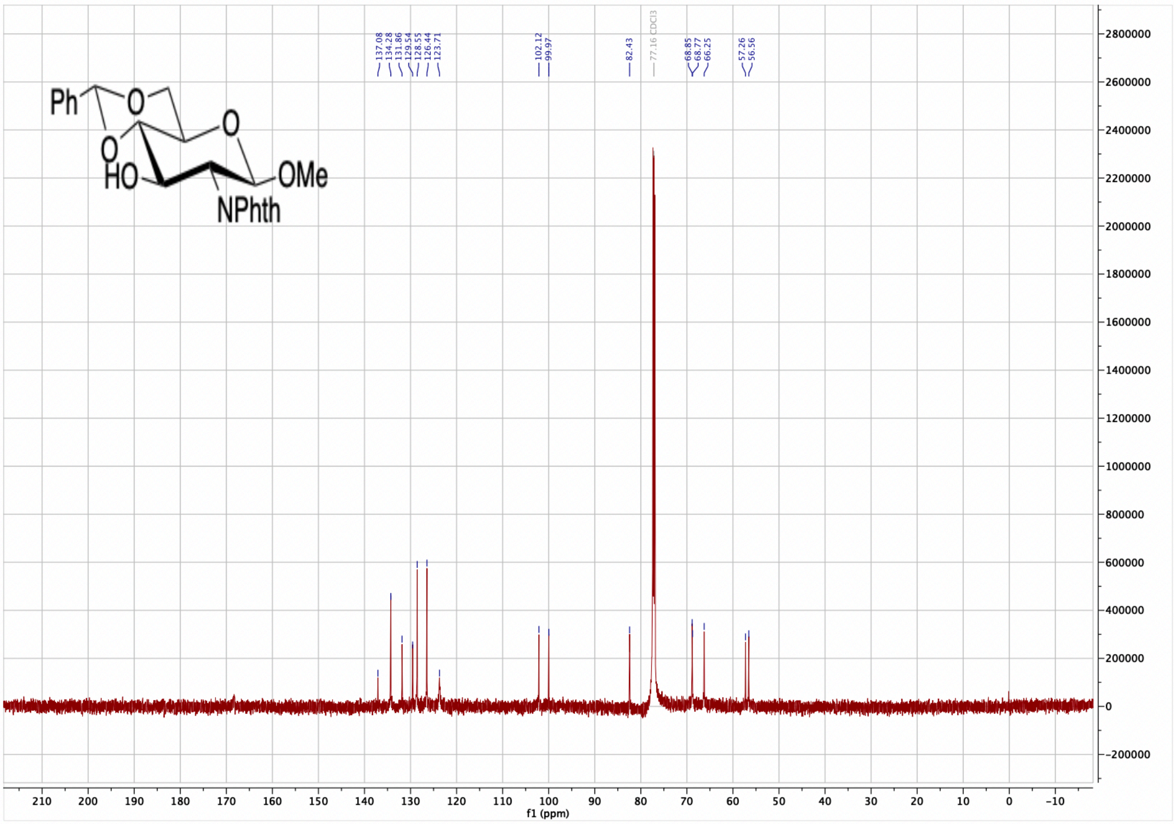

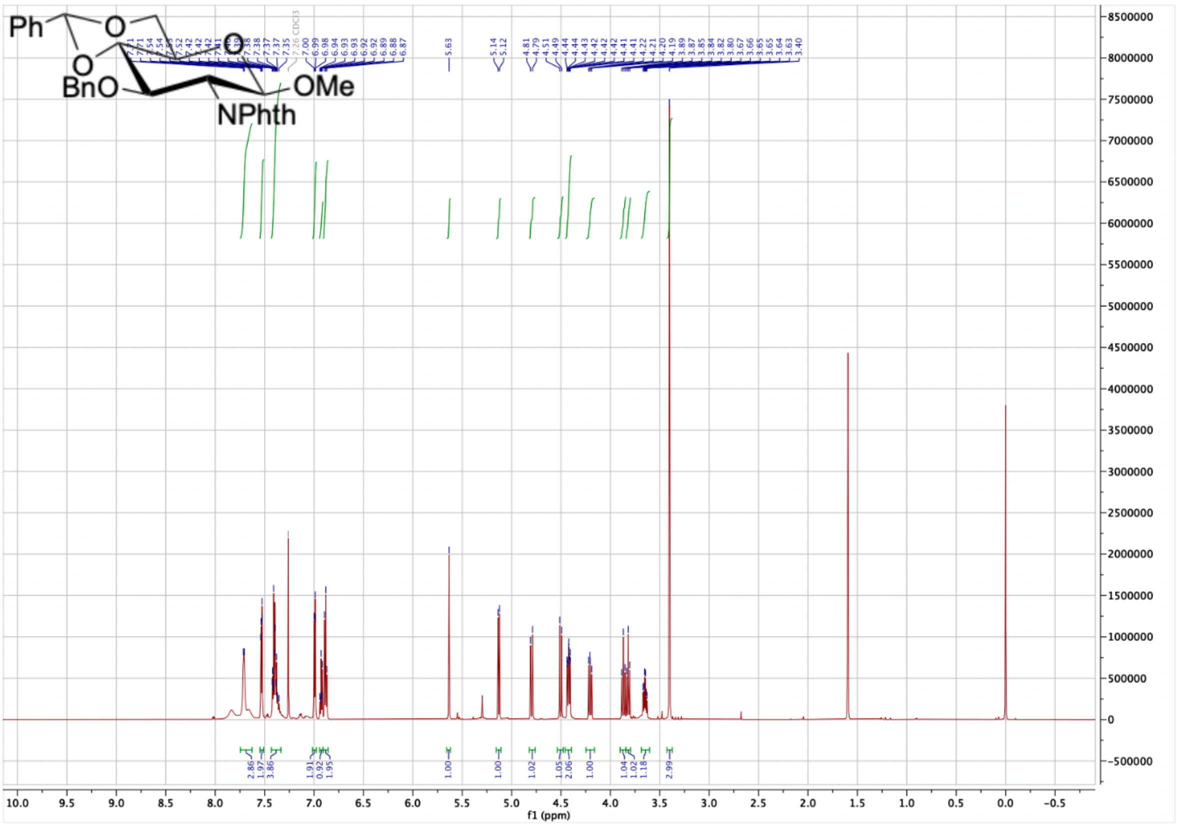

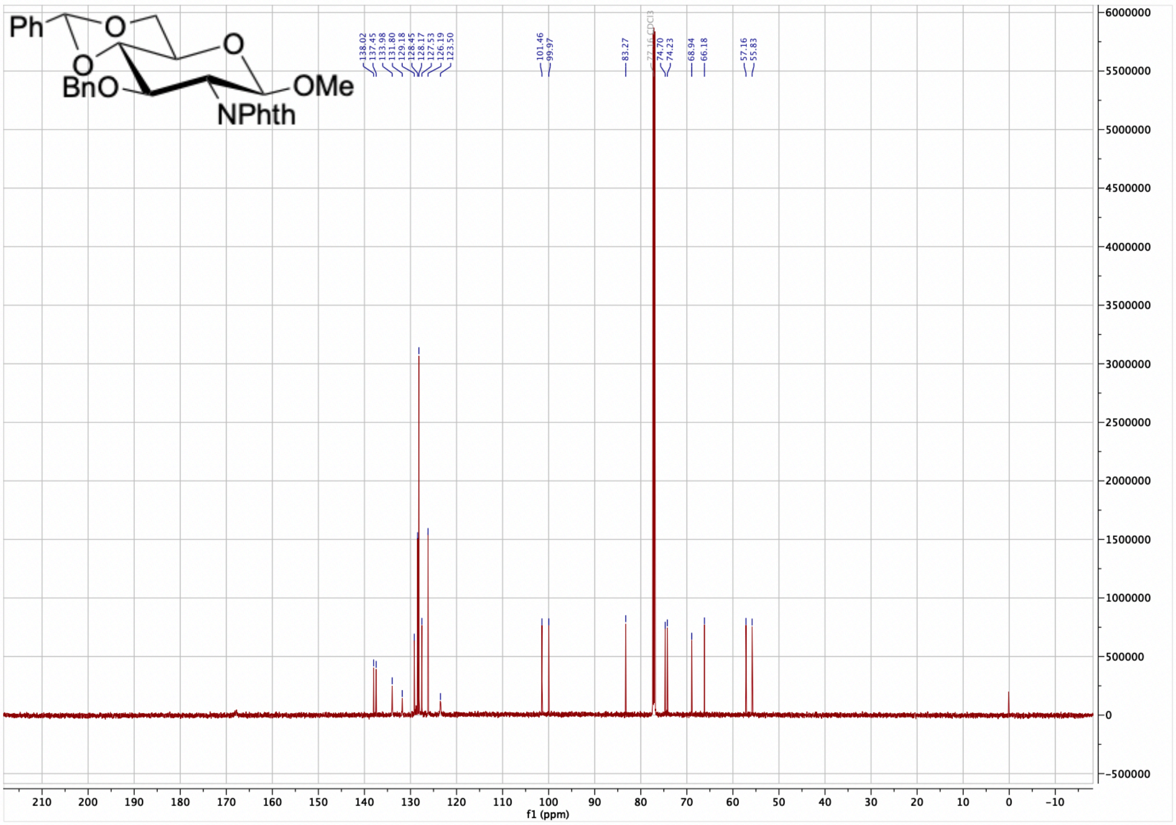

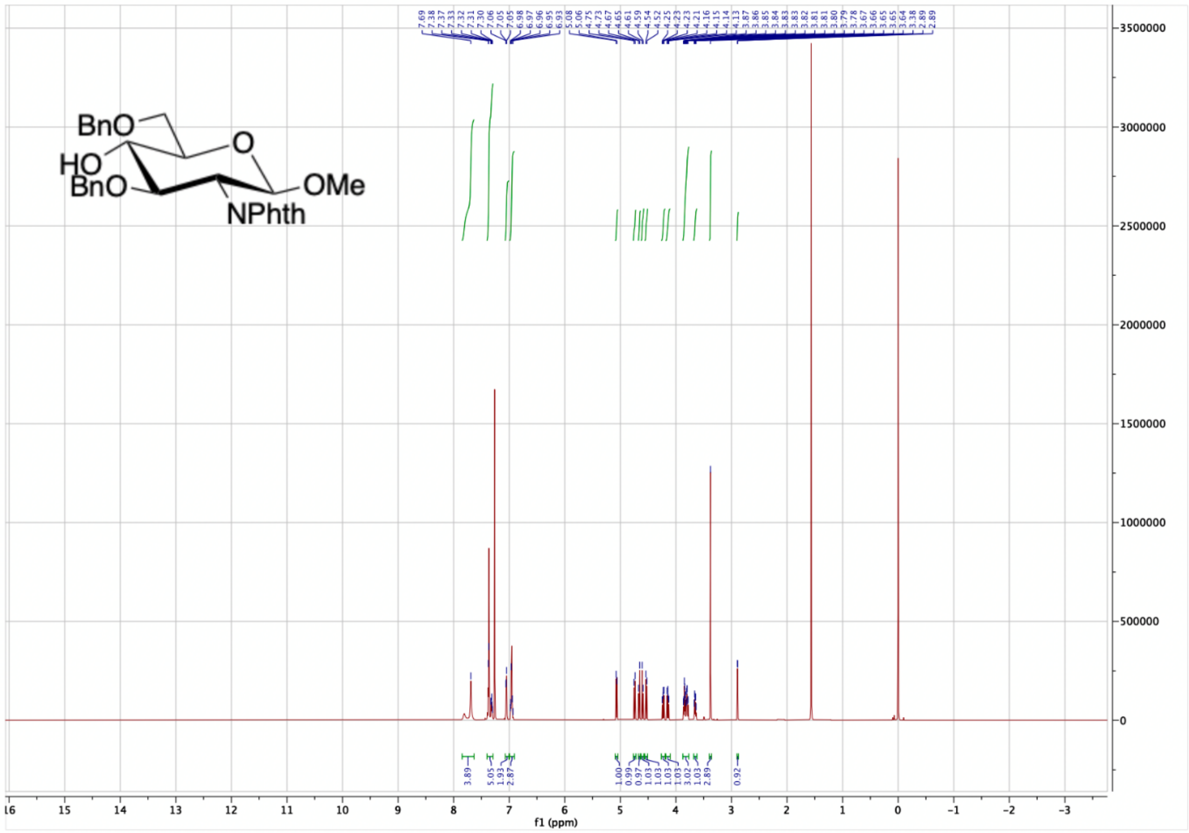

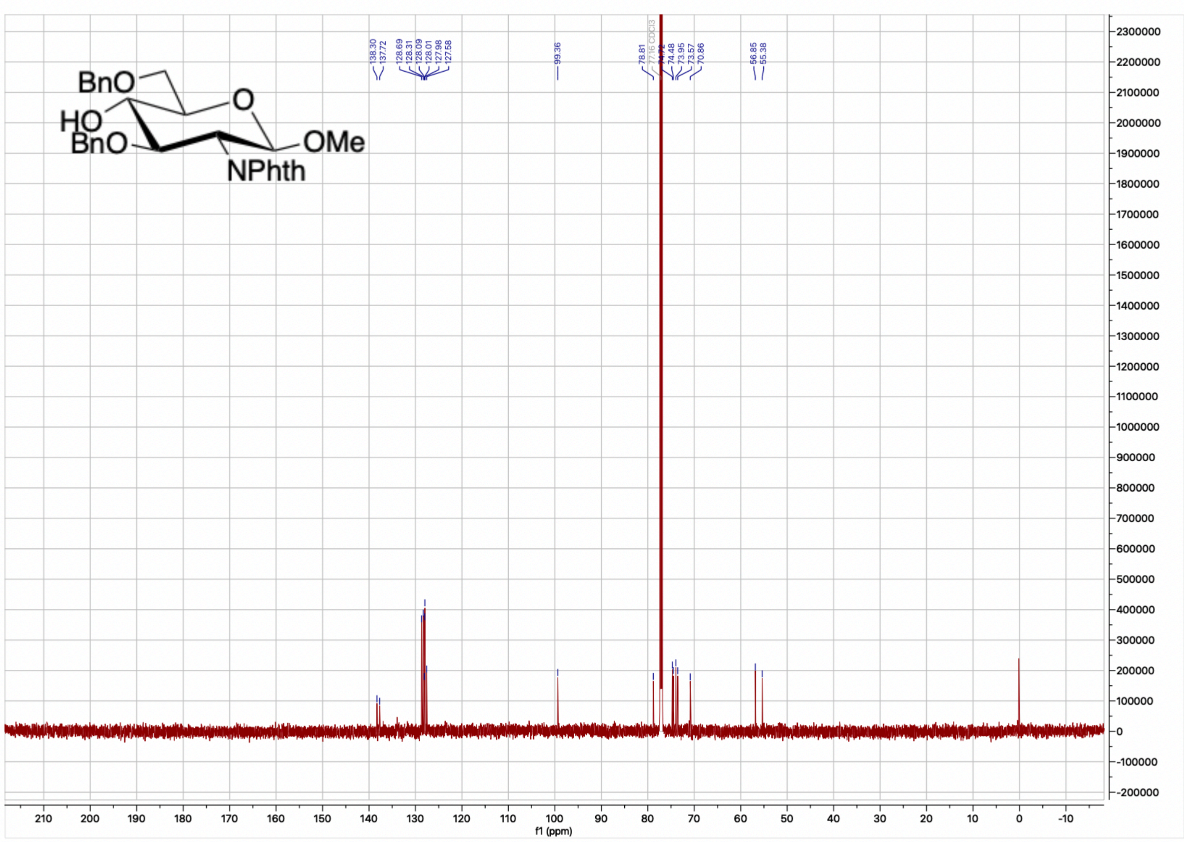

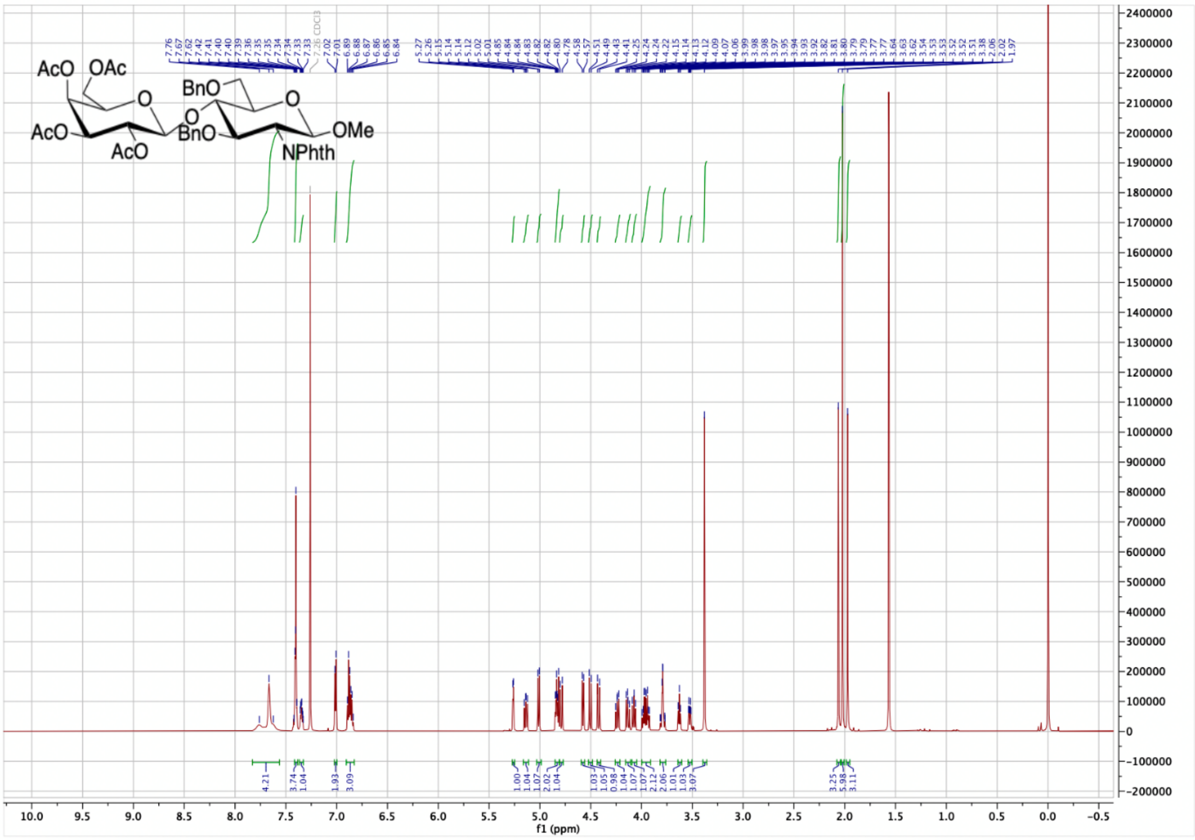

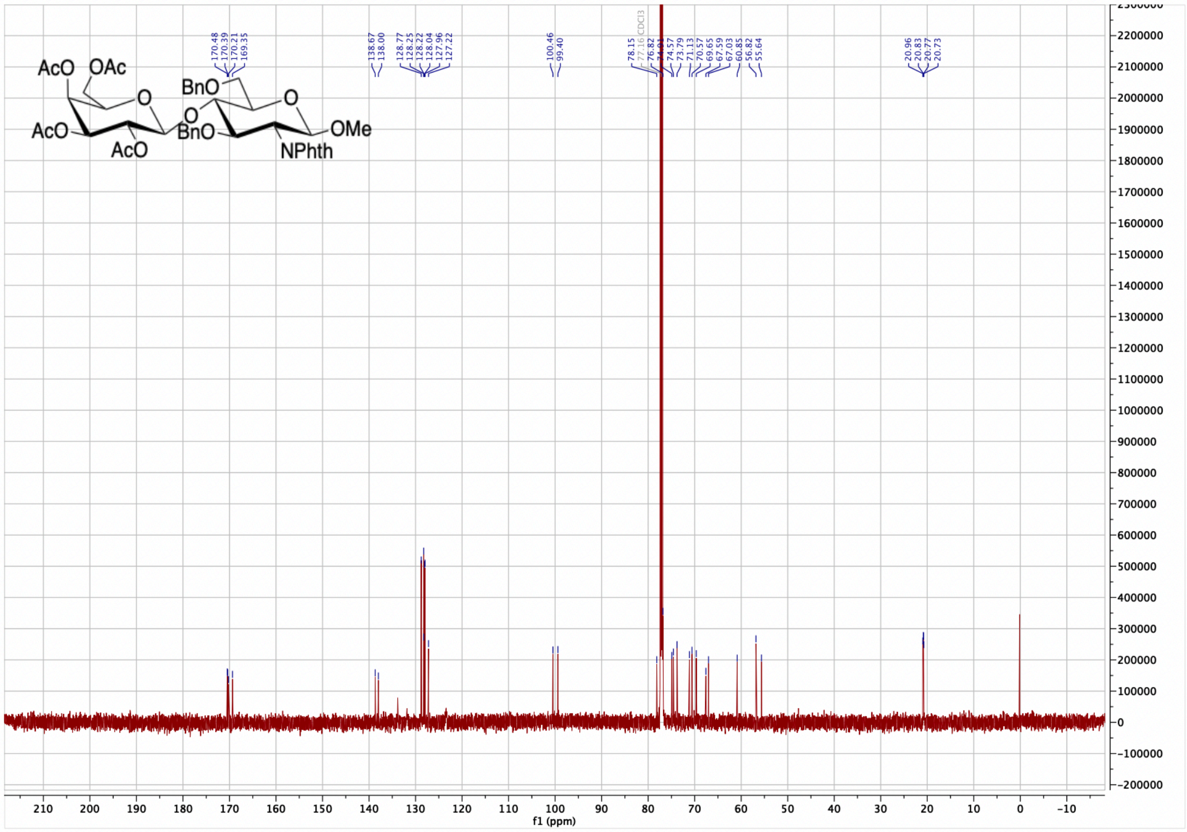

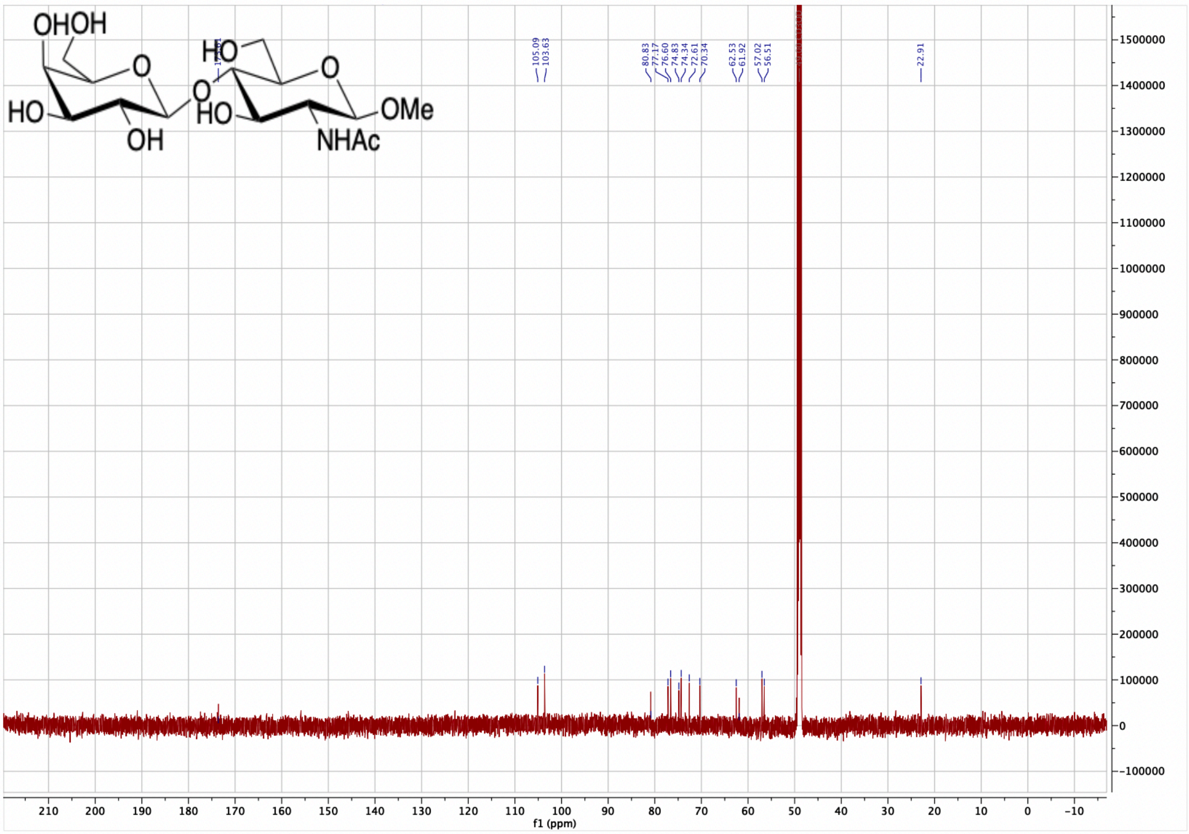

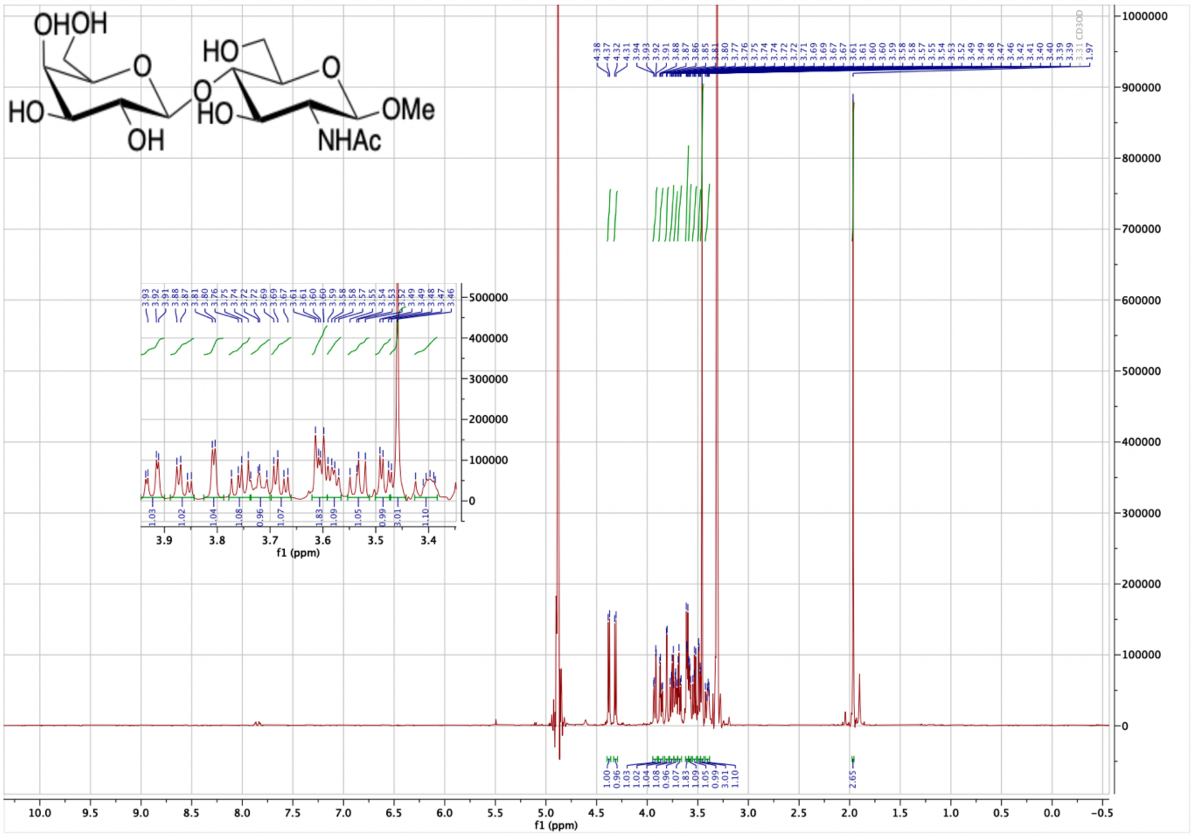
